# The Potent PHL4 Transcription Factor Effector Domain Contains Significant Disorder

**DOI:** 10.1101/2024.06.27.601048

**Authors:** Blake D. Fonda, Dylan T. Murray

**Author notes:** Corresponding Author.; Address: 91 N. Eagleville Road, Unit 3125, Storrs, CT 06269, USA. Description of Supplementary Material: Purification and sequence information for protein constructs; additional data and analysis from the denaturing assay; mass spectrometry data; additional NMR diffusion data and analysis; additional NMR spectra and analysis; disorder and structure predictions; tables of NMR data acquisition parameters, assigned NMR chemical shifts, and unassigned chemical shifts.

## Abstract

The phosphate-starvation response transcription-factor protein family is essential to plant response to low-levels of phosphate. Proteins in this transcription factor (TF) family act by altering various gene expression levels, such as increasing levels of the acid phosphatase proteins which catalyze the conversion of inorganic phosphates to bio-available compounds. There are few structural characterizations of proteins in this TF family, none of which address the potent TF activation domains. The phosphate-starvation response-like protein-4 (PHL4) protein from this family has garnered interest due to the unusually high TF activation activity of the N-terminal domain. Here, we demonstrate using solution nuclear magnetic resonance (NMR) measurements that the PHL4 N-terminal activating TF effector domain is mainly an intrinsically disordered domain of over 200 residues, and that the C-terminal region of PHL4 is also disordered. Additionally, we present evidence from size-exclusion chromatography, diffusion NMR measurements, and a cross-linking assay suggesting full-length PHL4 forms a tetrameric assembly. Together, the data indicate the N- and C-terminal disordered domains in PHL4 flank a central folded region that likely forms the ordered oligomer of PHL4. This work provides a foundation for future studies detailing how the conformations and molecular motions of PHL4 change as it acts as a potent activator of gene expression in phosphate metabolism. Such a detailed mechanistic understanding of TF function will benefit genetic engineering efforts that take advantage of this activity to boost transcriptional activation of genes across different organisms.

**Significance:** Transcription factor proteins upregulate genes and are essential to concerted biological response to environmental conditions like stress or low nutrient availability. In this work, we show the activating effector domain of the potent PHL4 transcription factor protein is primarily disordered, without well-defined secondary structure, and that the isolated effector domain behaves similarly in isolation as it does in the full-length protein. Our finding is consistent with protein transcription factors often having regions of disorder within their functional activator domains.

## Introduction

Despite their ubiquitous role in biological processes, a mechanistic molecular understanding of how DNA-binding TF proteins alter gene expression is elusive. The action of DNA-binding TF proteins is generally due to a two-component architecture consisting of a DNA binding domain and a separate effector domain (Staby et al., 2017). Effector domains, either activating or repressing transcription, have been shown to be functionally interchangeable between TFs (Hummel et al., 2023). For example, genetically engineered chimeric proteins with DNA binding and transcriptional effector domains from different proteins modulate transcription according to the nature of the effector domain for the target gene of the DNA binding domain (Hummel, 2023).

At the molecular level, it is unclear how these effector domains modulate transcription. Some proposals implicate proline rich, glutamine rich, or acidic regions within the effector domain. For instance, the presence of acidic regions activate gene transcription, while deletion of acidic regions decrease transcription rates (Sanborn et al., 2021; Sigler, 1988; Soto et al., 2022). These effector domains increase transcription via interaction with partner proteins such as mediator but also interact with other regulatory proteins that inhibit their effect on transcription (Sanborn et al., 2021). For favorable protein-protein interactions to be possible between a TF effector domain and several different proteins, the ability to populate several distinct conformations is likely beneficial. Indeed, conformational flexibility has been evolutionarily selected for on a genome wide scale for the effector domains of transcription factors (Soto, et al. 2022). Thus, intrinsically disordered regions within effector domains of TFs have been hypothesized to be causative of modulating gene transcription.

The Arabidopsis TF phosphate starvation response like protein-4 (PHL4) belongs to a family of homologous phosphate-starvation response TF proteins (Wang et al., 2018). The best characterized family member, and the one with the closest sequence similarity to PHL4, is the protein phosphate starvation response-1 (PHR1), believed to be the central TF that is most-essential to the plant response to low phosphate availability (Rubio et al., 2001). All family members (PHR1, PHL1, PHL2, PHL3, and PHL4) share highly conserved amino acid sequences in the DNA binding and coiled-coil domains (Bustos et al., 2010; Wang, et al., 2018). Like PHR1, although to a different degree depending on the gene target, PHL4 can increase the transcription of these same genes in the presence of low phosphate levels (Wang et al., 2018). PHL4 is therefore functionally similar to the central TF of the phosphate response, PHR1.

While the coiled-coil and DNA binding domains explain promoter binding and cooperative enhancement of gene expression, these domains do not fully explain the interaction of PHR1 and PHL4 with the transcriptional machinery. The first 227 amino acids of PHL4 are not predicted to adopt a folded conformation by AlphaFold 2.0 (Jumper et al., 2021; Varadi et al., 2022), yet have significant sequence similarity, 48%, with PHR1. PHL4 has an extra 37 residues at the N-terminus, which could be related to additional functional activity relative to PHR1 (Wang, 2018). These first 227 residues of PHL4, the effector domain, were among the most potent in an Arabidopsis-based screen of effector domains for 400 TF proteins, in some cases inducing gene expression more than the state-of-the-art bioengineering effector domain VP16 (Hummel et al., 2023). The N-terminal effector domain of PHL4 is therefore likely an important component of transcriptional activation. In addition to the native function of PHL4, it is also a promising candidate for creating artificial TF proteins in genetically modified organisms (Hummel et al., 2023). It is highly active in multiple eukaryotes including yeast and N. benthamiana (Hummel et al., 2023), promising broad applicability of the PHL4 effector domain as a molecular tool to increase gene expression in genetically engineered crops. Additionally, the PHL4 effector domain is an important case study, helping to build a consensus regarding the shared features of potent TF proteins across eukaryotic organisms.

To the best of our knowledge, despite the importance of PHL4 in phosphate response pathways and as an exceptional transcriptional activator, there is no published structural characterization of the protein. As a starting point for dissecting the protein-protein and protein-nucleic acid interactions of PHL4 as it carries out TF activity, we characterize the structure and conformational flexibility of the protein, with particular focus on the N-terminal effector domain. Fluorescence and circular dichroism measurements report on global structure and order. Cross-linking, chromatography, and diffusion NMR measurements probe higher order oligomeric assembly. Solution NMR measurements determine residue-specific conformations and timescales of motion. Together these results form a foundation for addressing changes in these properties as PHL4 carries out TF activity.

## Results

Pure preparations of full-length, wild-type PHL4 (herein referred to as PHL4_FL_) and the first 227 N-terminal residues (herein referred to as PHL4_effector_), were made using recombinant expression in E. coli. Each construct, illustrated in Supplementary Figure 1, contains a N-terminal 6x His tag with a tobacco etch virus (TEV) cleavage site that was removed before subsequent experiments. Following TEV cleavage, reverse phase affinity column chromatography yielded highly pure PHL4_FL_ and PHL4_effector_ as observed by SDS-PAGE in Supplementary Figure 1. The domain map in Figure 1a shows PHL4_FL_ follows the typical architecture of a TF, with an effector domain, DNA binding domain, and accessory domains (Staby et al., 2017).

**Figure 1.**
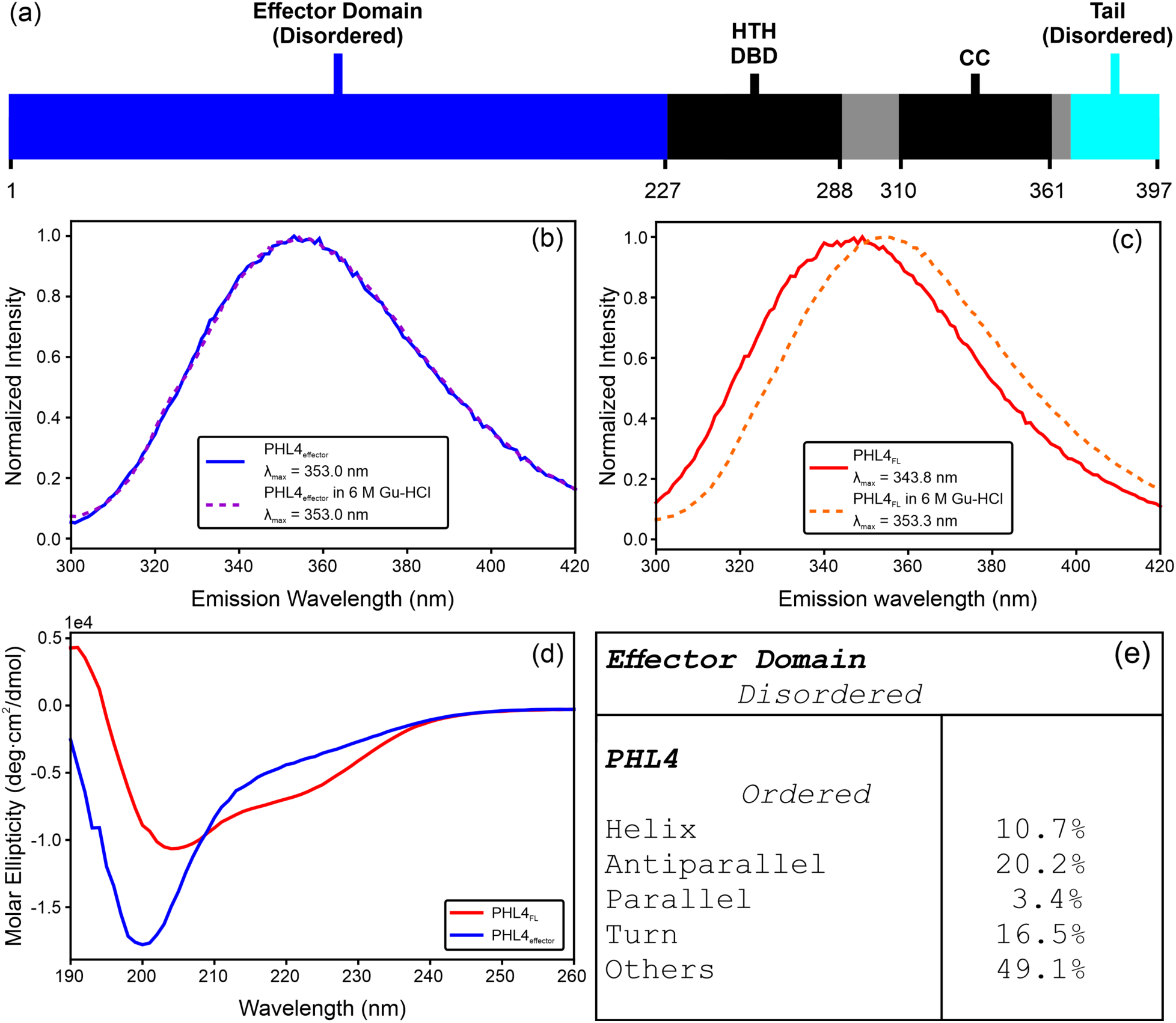
Bulk secondary structure content in PHL4. (a) A domain map for PHL4. HTH DBD is the homology predicted helix-turn-helix DNA binding domain, CC is the homology predicted coiled-coil domain, and residues 1–227 and 369–397 are to be shown in this work as primarily disordered. (b) Intrinsic Trp fluorescence emission spectra for PHL4_effector_ with and without 6 M Gu-HCl. (c) PHL4_FL_ intrinsic Trp fluorescence spectra with and without 6 M Gu-HCl. For both (c) and (d), the spectra are normalized so the maximal intensity is 1.0. (d) CD spectra of PHL4_FL_ in red and PHL4_effector_ in blue recorded in non-denaturing conditions. (e) A summary of the analysis of the CD spectra in (d) using BeStSel. “Others” in the BeStSel analysis encompasses disordered regions, 3_10_-helix, π-helix, β-bridge, bend, and loop regions.

As an initial probe of structure formation in PHL4, we compared intrinsic Trp fluorescence in native and denaturing conditions. Figure 1b shows spectra in non-denaturing and guanidine hydrochloride (Gu-HCl) denaturing conditions for PHL4_effector_. No change in the spectral line shape or position is observed and the intensity maximum (λ_max_) is 353.0 nm, close to that of a fully water exposed Trp of 355.0 nm (Royer, 2006). Thus, the Trp residues in the PHL4_effector_ are solvent exposed in both conditions. However, Figure 1c shows spectra of PHL4_FL_ revealing an intensity maximum shift from 343.8 to 353.3 nm when the buffer is changed from native-like to denaturing. This shift is consistent with a reduction in solvent accessibility for Trp residues in PHL4_FL_ in non-denaturing conditions. However, 343.8 nm is intermediate between the fully water exposed value (355 nm) and that of partially buried Trp (∼335 nm) (Royer, 2006), indicating not all Trp species in PHL4_FL_ are involved in structure formation in native-like buffer conditions. Of the three Trp residues in PHL4_FL_, two are in the effector domain at W113 and W149, and the third is residue W236 in the DNA binding region. The intrinsic Trp fluorescence spectra for PHL4_FL_ and PHL4_effector_ are therefore consistent with a structured conformation around W236, resulting in the partial burial away from solvent of this residue in PHL4_FL_, and residues W113 and W149 being solvent exposed and disordered in both PHL4_FL_ and PHL4_effector_.

An unfolding assay of PHL4_FL_ was then performed to estimate the ΔG_folding_. PHL4_FL_ was diluted into a series of buffered, near neutral pH, and increasingly denaturing Gu-HCl solutions and, after allowing the samples to equilibrate, intrinsic Trp fluorescence spectra were recorded. The mole fraction of folded protein at each denaturant concentration was calculated and fit to a sigmoidal relationship providing the apparent Gibbs energy change of folding, ΔG_folding_, values for PHL4_FL_ of −2.5 ± 0.3 kcal/mol based on intensity measurements or −2.3 ± 0.8 kcal/mol based on wavelength of maximum intensity (λ_max_) values following the analysis described in Monsellier and Bedouelle, 2005, and assuming a two state, folded to unfolded, model. The intrinsic Trp spectra and fitted curves are shown in Supplementary Figure 2. These ΔG_folding_ values are consistent with a weakly folded, or mostly disordered PHL4_FL_ with small regions of well-defined structure. In comparison, the protein lysozyme, for example, with roughly 250 *fewer* amino acid residues than PHL4, has a much more favorable ΔG_folding_ of −8.5 kcal/mol (Li-Blatter et al., 2019).

Circular dichroism (CD) was then used to investigate the secondary structure present in PHL4_FL_ and PHL4_effector_. CD uses the differential absorption of circularly polarized light by chiral protein structural motifs, like α-helix versus β-strand, to estimate global protein secondary structure content when compared with databases of CD spectra and known protein structures. Figure 1d shows the CD spectra of PHL4_effector_ and PHL4_FL_. Analysis of the spectra using BeStSel (Micsonai et al., 2022) with the “Disordered-Ordered” classification is summarized in Figure 1e. The prediction for the PHL4_effector_ spectrum is consistent with a disordered protein, while the prediction for PHL4_FL_ is consistent with a mixture of ordered secondary structure motifs accounting for approximately half of the protein. In this analysis, unfolded or regions with non-standard structure are indicated as “Other” regions and account for the remaining half of the PHL4_FL_ protein. The lack of secondary structure content determined by CD is consistent with the intrinsic Trp fluorescence results.

To characterize the oligomeric state of PHL4_effector_ and PHL4, mass-spectral analysis was performed with two different spectrometer types: electro-spray with an orbitrap detector (ESI-orbitrap) and matrix assisted laser desorption ionization with a time-of-flight detector (MALDI-TOF). Supplementary Figure 3 shows the mass spectra and analysis for both PHL4_FL_ and PHL4_effector_ which are consistent with primarily monomeric protein. However, these methods are subject to the caveat that the ionization methods used can break apart weakly associated oligomers or miss lowly populated states in the presence of a dominant monomeric species (Leney and Heck, 2017, Rogawski and Sharon, 2022).

Since PHL4_FL_ was shown by fluorescence and CD measurements to contain folded regions, a cross-linking SDS PAGE assay on PHL4_FL_ was ran to confirm the oligomeric state. Figure 2a demonstrates that the PHL4_FL_ monomer band runs on the gel as a 56 kDa mass, higher than the 44 kDa calculated from the protein primary sequence, which is also consistent with the SDS PAGE gel used to analyze the PHL4_FL_ purification shown in Supplementary Figure 1. Distinct, higher molecular weight bands are also observed in the crosslinking gel in Figure 2a. Dividing the mass of the higher molecular weight bands by the 56 kDa monomer mass allows for the determination of oligomeric species present in each sample. These oligomeric states are depicted in Figure 2b–d as dimer, tetramer, and higher oligomeric species. The lack of a band corresponding to a trimer indicates that PHL4_FL_ is likely not a monomer or a trimer in solution, but rather is an even numbered oligomer, consistent with a dimer, tetramer, or higher state. The strong monomer band on the gel is unsurprising given that the gel sample conditions (8% w/v sodium dodecyl sulfate) can break apart any weakly associated complexes not covalently linked by glutaraldehyde.

**Figure 2.**
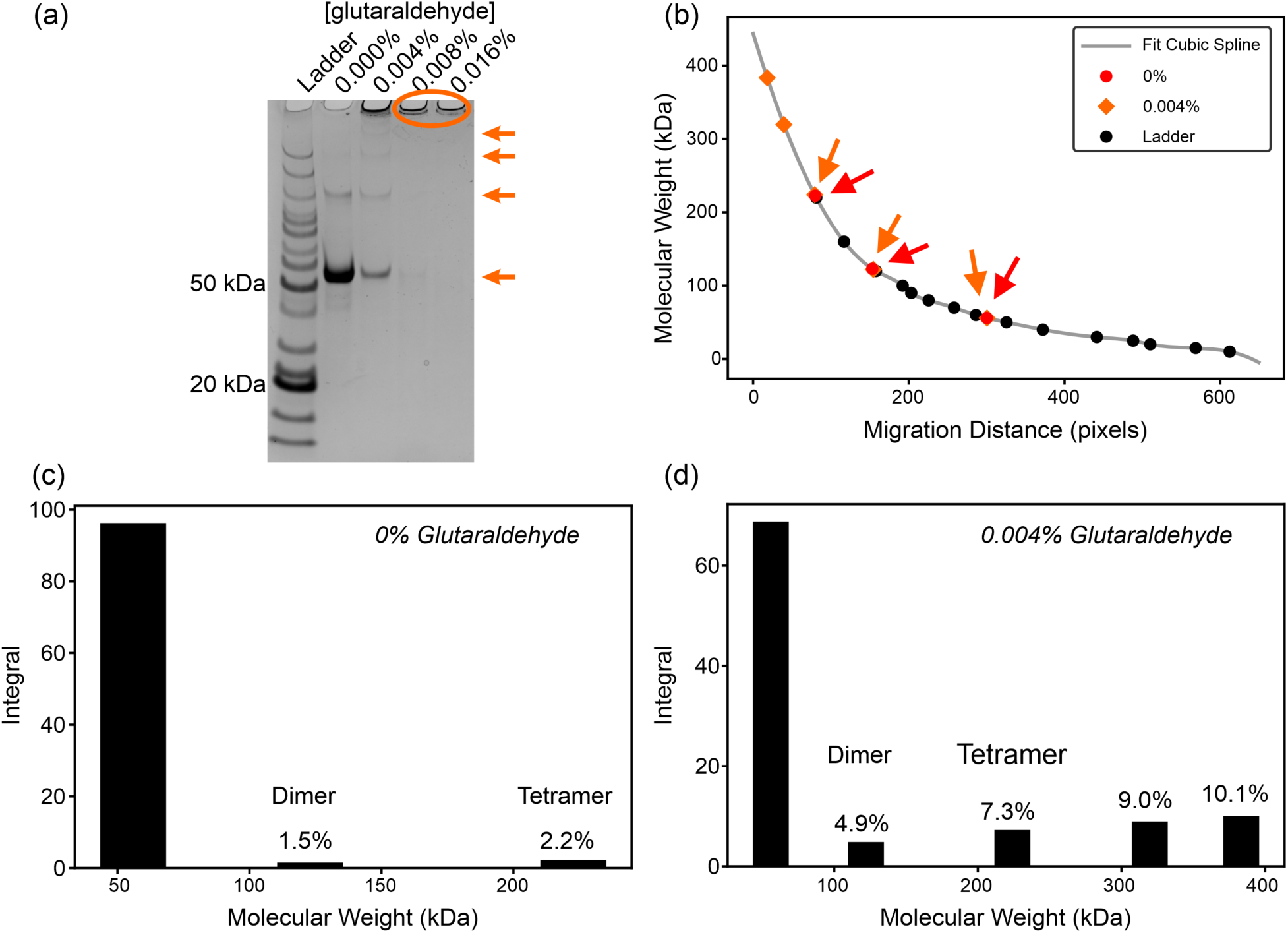
PHL4_FL_ cross-linking assay shows a lack of odd numbered oligomeric species. (a) PHL4_FL_ incubated with increasing percentages of glutaraldehyde, doubling the concentration of glutaraldehyde in each successive lane, run on a 4–10% Bis-acrylimide gradient SDS PAGE gel. Arrows point to oligomeric species detected in 0.004% glutaraldehyde and the circle highlights aggregated protein unable to enter the gel. (b) Plot of the molecular weights of the bands observed in the gel in panel (a), arrows point to bands observed in both the 0% and 0.004% lanes. (c) and (d) Bar plots of the integrated gel band intensity versus the molecular weight, measured from the cubic spline fit to the data in panel (b). Dimer and tetramer are based on the observed molecular weight divided by the intense monomer band at 56 kDa. The percentages normalized so that the sum of all band intensities equals 100%.

To further clarify the oligomeric state of PHL4_FL_, size exclusion chromatography (SEC) was performed in non-denaturing and pH 7.4 buffered solution. Figures 3a–b shows the chromatograms for PHL4_effector_ and PHL4_FL_ compared with protein molecular weight standards ran under identical buffer conditions. Since PHL4_effector_ and PHL4_FL_ contain disordered motifs as indicated by CD and fluorescence microscopy, the chromatograms were therefore used to calculate the radius of hydration, R_h_, not molecular weight. This approach helps account for larger than expected R_h_ values for a given protein molecular weight due to extended conformations of disordered domains. Figure 3c shows the R_h_ and retention time relationship used to estimate R_h_. For the protein standards, R_h_ values were obtained from the literature (Stetefeld et al., 2016; Talmard et al., 2007; Wilkins et al., 1999). PHL4_effector_ and PHL4_FL_ sizes measured by SEC are tabulated in Table 1.

**Figure 3.**
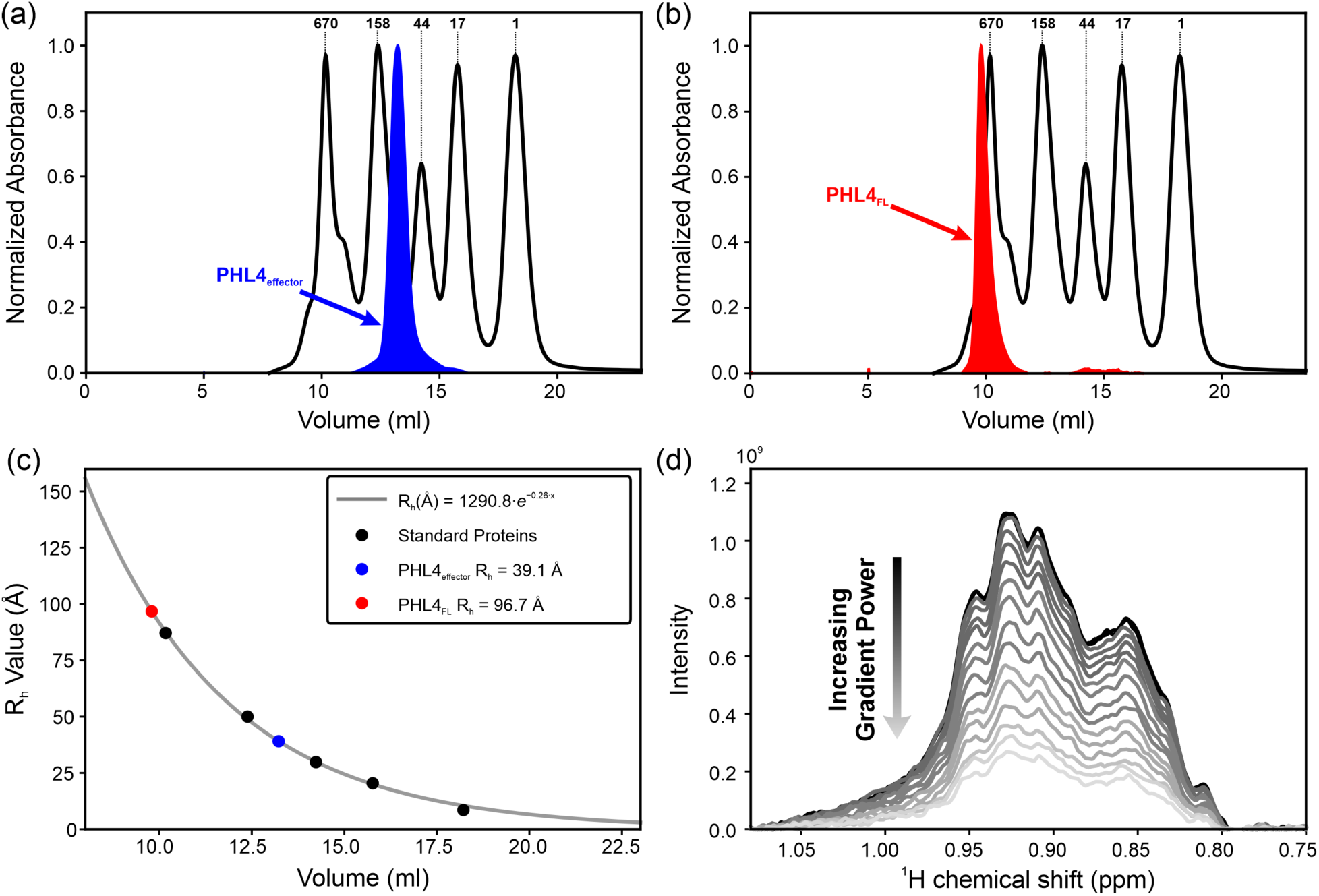
Radius of hydration measurement for PHL4_FL_ and PHL4_effector_. (a) SEC of PHL4_effector_ in blue, overlayed with a chromatogram of standard proteins ran in the same conditions in black, with the labels corresponding to the molecular weights specified by the manufacturer. (b) SEC of PHL4_FL_ in red versus standards ran in the same conditions. (c) Plot of the radius of hydration, R_h_, reported in the literature for the SEC standards (Stetefeld et al., 2016; Talmard et al., 2007; Wilkins et al., 1999) as a function of retention volume with a best-fit exponential decay function. The R_h_ values for PHL4_FL_ and PHL4_effector_, calculated from the best-fit function, are shown and red and blue circles. (d) Region of PHL4_effector_ diffusion NMR ^1^H NMR spectrum integrated to determine R_h_.

**Table 1.** Radius of hydration predicts size and folded state for each protein. The SEC result is the value calculated from the measured SEC retention time. The diffusion NMR NMR result is calculated from the gradient-strength based decay observed in the ^1^H NMR spectra. *^a^*Uncertainty is the standard deviation determined for the best-fit exponential for the retention times of the standard samples. *^b^*Uncertainty is the standard deviation from the R_h_ value calculated from three separate diffusion NMR measurements on each sample. ^c^from Dudás and Bodor, 2019.

| Protein Motif | SEC $R_h$ (nm)<sup>a</sup> | Diffusion NMR $R_h$ (nm)<sup>b</sup> | Predicted Folded to Unfolded $R_h$ Range<sup>c</sup> | Predicted Oligomeric State |
| --- | --- | --- | --- | --- |
| PHL4<sub>FL</sub> | $97.0 \pm 1.0 \text{ \AA}$ | $86.9 \pm 9.7 \text{ \AA}$ | **Monomer:** 33.5–59.5 $\text{\AA}$ | Partially folded timer to octamer |
| | | | **Dimer:** 43.7–83.7 $\text{\AA}$ | |
| | | | **Trimer:** 51.0–102.1 $\text{\AA}$ | |
| | | | **Tetramer:** 67.0–117.7 $\text{\AA}$ | |
| | | | **Octamer:** 74.2–165.5 $\text{\AA}$ | |
| PHL4<sub>effector</sub> | $39.1 \pm 0.9 \text{ \AA}$ | $40.4 \pm 0.1 \text{ \AA}$ | **Monomer:** 27.1–45.2 $\text{\AA}$ | Unfolded monomer |
| | | | **Dimer:** 35.3–63.6 $\text{\AA}$ | |

To corroborate the SEC-based R_h_ measurement, diffusion NMR spectroscopy measurements were made as an additional measure of R_h_. The diffusion NMR measurement uses a series of increasing strength gradient pulses during a ^1^H NMR experiment causing a signal decay. Figure 3c shows the decay observed for PHL4_effector_. The rate of decay is related to the diffusion of each molecular species whose nuclei give rise to NMR peaks in the spectrum. The rate of diffusion for the protein is calibrated using the ^1^H NMR signal decay for an internal 1,4-dioxane standard. The diffusion rate data is then used to calculate R_h_. Additional spectra and analysis are presented in Supplemental Figures 4–6. Diffusion NMR measured R_h_ values are also tabulated in Table 1. Data for horse myoglobin is included as a control with a known molecular weight and radius of hydration, and published diffusion NMR data.

The R_h_ measured using diffusion NMR and SEC are in agreement, in light of established literature showing that SEC and diffusion NMR R_h_ measurements likely carry uncertainties of a few Å for the small proteins lysozyme (20 ± 2 Å) and ovalbumin (30 ± 1.4 Å) (Dudás and Bodor., 2019; Parmar et al., 2009). Extrapolation of those uncertainties to the size of PHL4_FL_ measured here suggest that the uncertainty is on the order of ±6 Å (Dudás and Bodor, 2019; Parmar and Muschol, 2009). The slight difference in R_h_ calculated between diffusion NMR and SEC methods also makes sense given these R_h_ analyses rest on a spherical approximation of protein shape. Additionally, potential protein-column interactions during the flow of proteins through porous resin during SEC could alter the observed R_h_.

The R_h_ values from both the SEC and diffusion NMR analyses are compared to known relations between the size and number of residues in a protein. The upper and lower bounds in Table 1 come from the estimated R_h_ calculated from analytical relationships for completely folded proteins (lower bound) and for unfolded proteins (upper bound) (Dudás and Bodor, 2019). This calculation gives a range of R_h_ values based on the number of amino acid residues for each possible oligomeric state of PHL4_effector_ and PHL4_FL_. For PHL4_effector_, the R_h_ measured using both SEC and diffusion NMR is consistent with monomeric, unfolded protein. For PHL4, on the other hand, the measured R_h_ value consistent with an oligomer bigger than a dimer.

The molecular weight estimates from diffusion NMR and SEC, the CD and fluorescence analysis of structural disorder, the cross-linking assay showing that the predominate species is an even integer multiple, and the unfolding assay showing a small ΔG_folding_, suggest that PHL4_FL_ is to likely be between a tetramer and an octamer with a mixture of folded and unfolded regions. We tentatively propose that these results are consistent with a tetrameric state, which would be consistent with the result for the homologous PHR1 protein, for which the coiled-coil domain forms tetramers (Ried et al., 2021). The tetrameric state upper bound is near the value calculated for a completely unfolded PHL4_FL_, which is compatible with the domain structure of PHL4_FL_ in Figure 1a showing the structured coiled-coil and helix-turn-helix domains occupying only ∼28% of the PHL4_FL_ sequence and our solution NMR measurements showing the N- and C-termini are disordered (vide infra). For PHL4_effector_, the data robustly show that the effector domain in isolation is a monomer.

To gain structural insight at the level of individual amino acid residues, solution NMR was used to measure the chemical shifts of individual atoms in the protein. The NMR chemical shift values report on both amino acid type and protein secondary structure. Sequence specific assignments are necessary to use NMR to probe structural conformations present in our PHL4_FL_ and PHL4_effector_ samples. The NMR chemical shifts from each protein sample were recorded using a standard set of triple-resonance solution NMR experiments. Figure 4a–b shows ^1^H–^15^N HSQC spectra for PHL4_effector_ and PHL4_FL_ annotated with the sequence specific assignments. These experiments and the data acquisition parameters are listed in Supplementary Table 1. Sequence specific assignments of the NMR chemical shifts were made manually for PHL4_effector_ sequence, with 72% of the 227 residues in PHL4_effector_ assigned. For PHL4_FL_, 59% of the N-terminal effector domain was assigned and 100% of the C-terminal residues from G369–E397. Out of all observed signals for PHL4_effector_, 73% were assigned, whereas for PHL4_FL_, 62% of the signals were assigned. Many of these unassigned peaks have low signal-to-noise, as shown in Figure 4 and described in Supplementary Tables 3–6.

**Figure 4.**
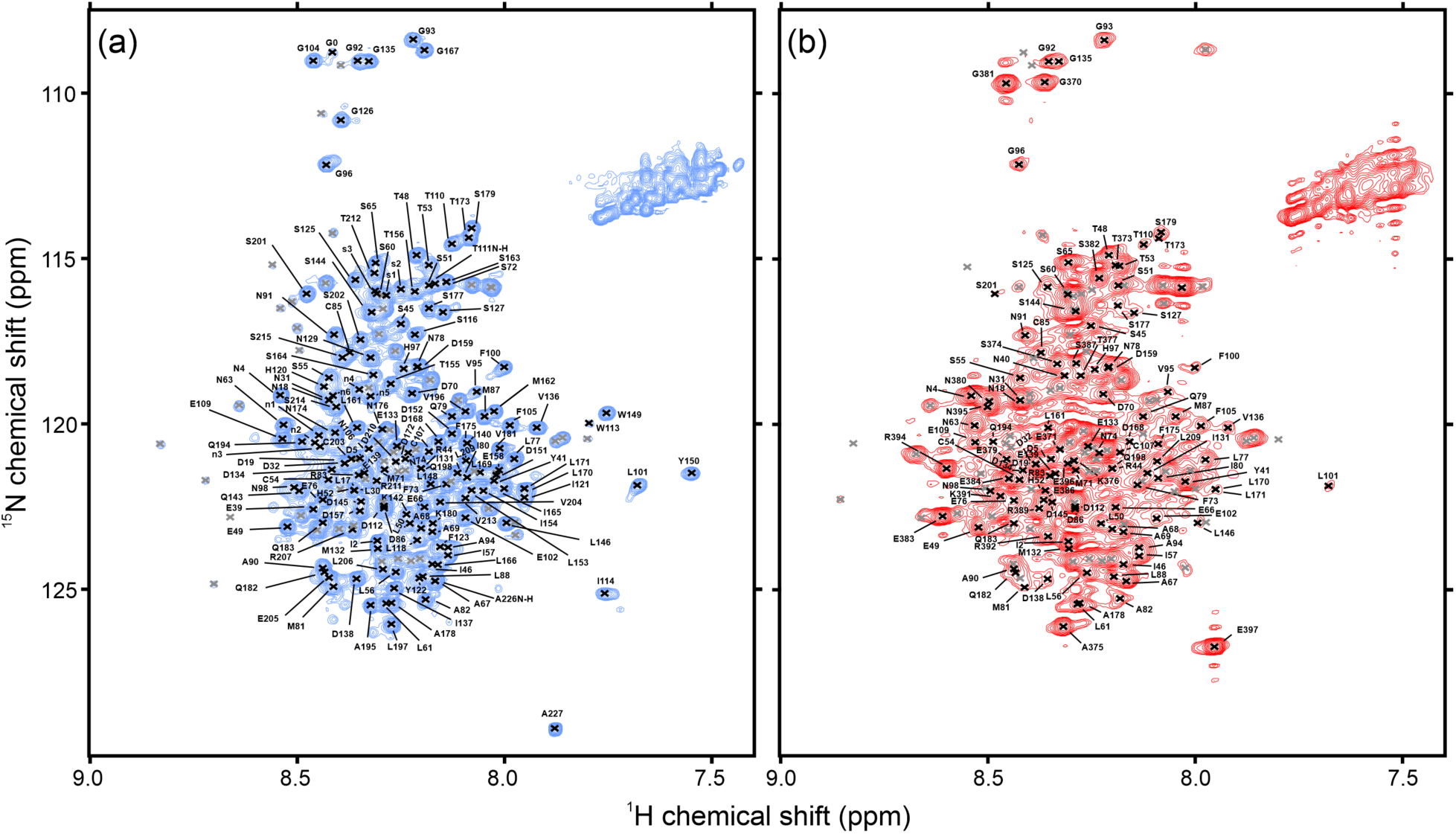
^1^H–^15^N HSQC spectra for PHL4_effector_ and PHL4_FL_. (a) ^1^H–^15^N HSQC NMR spectrum of PHL4_effector_. (b) ^1^H–^15^N HSQC NMR spectrum of the PHL4_FL_. Grey peaks represent unassigned peaks. “n” and “s” labels depict ambiguously assigned Asn and Ser peaks that have unique ^1^H and ^15^N shifts but non-unique “*i“* and “*i* − *1”* carbon chemical shifts.

For strong signals, the remaining unassigned NMR chemical shifts for PHL4_effector_ and PHL4_FL_ are primarily due to ambiguity in amino acid sequence. Supplemental Figures 1c–d shows the N-terminal region contains an acidic region with an almost identical 12–13 amino acid sequence repeated three times. Other unassigned NMR chemical shifts are likely due to the well folded regions in the PHL4_FL_ oligomer having large global rotational correlation times or slow local molecular motions, leading to reduced signal-to-noise in the NMR experiment and therefore a lack of connecting peaks observed in the spectra used for assignments.

The spectra in Figures 4a and 4b are highly similar to each other. The assigned NMR chemical shifts for PHL4_effector_ appear at a nearly identical NMR chemical shift values in the full-length PHL4_FL_ spectra for many residues. Supplementary Figure 7 quantifies the similarity in NMR chemical shifts via chemical shift perturbations, CSPs. Small CSPs indicate a lack of large structural differences in the assigned regions of PHL4_FL_ and PHL4_effector_. In sum, the solution NMR measurements indicate the effector domain in isolation adopts similar conformations and has similar molecular motions as the effector domain *within* the full-length PHL4.

Secondary chemical shifts are used to predict protein secondary structure from the assigned NMR chemical shifts. Secondary chemical shifts are the difference between the observed NMR chemical shift value for a residue and the random coil conformation NMR chemical shift value for that amino acid type (Kjaergaard et al., 2011a, Kjaergaard et al., 2011b). For PHL4_effector_ and PHL4_FL_, the secondary chemical shifts are represented as the difference between the CA and CB secondary chemical shifts in Figures 5a and 5c. Large positive values over several adjacent residues indicates an α-helical structure, while large negative values for neighboring residues are consistent with a β-strand structure (Wishart, 2011). These plots show that the majority of the secondary chemical shift magnitudes are less than 2, a commonly used cutoff for well-defined structure (Wishart, 2011). The secondary chemical shifts are therefore consistent with the assigned residues in both PHL4_FL_ and PHL4_effector_ lacking well-defined secondary structure.

**Figure 5.**
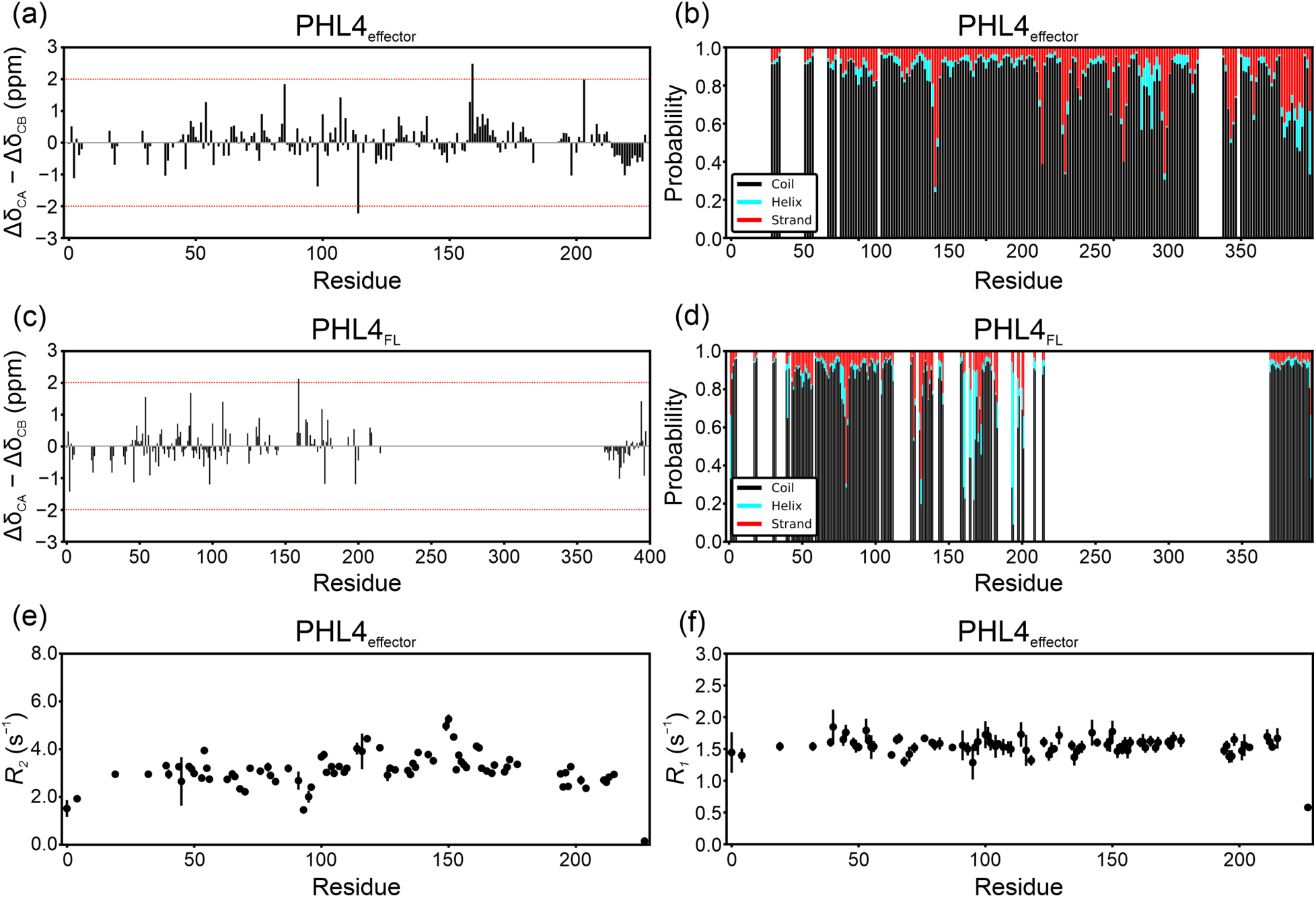
NMR chemical shift assignments and ^15^N relaxation rates show a random coil-like conformation. (a) PHL4_effector_ secondary chemical shift represented as the difference in CA and CB secondary chemical shifts. (b) TALOS-N predicted secondary structure for PHL4_effector_ based on the NMR chemical shift assignments. (c) PHL4_FL_ secondary chemical shifts represented as described for (a). (d) TALOS-N predicted secondary structure for PHL4_FL_ based on the NMR chemical shift assignments. (e) PHL4_effector_ *R_2_* relaxation rates calculated from *R_1_* and *R_1_*ρ measurements. (f) Measured PHL4_effector_ *R_1_* relaxation rates.

Additional structural information was obtained from a TALOS-N analysis of the assigned NMR chemical shifts. TALOS-N uses an artificial neural-network with a database of known protein NMR chemical shifts, structures, and sequences to predict secondary structure and backbone torsion angles based on NMR chemical shift assignments and protein sequence (Bax et al., 2013; Shen et al., 2009). The secondary structure predicted for PHL4_effector_ and PHL4_FL_ are depicted in figure 5b and 5d. Overwhelmingly, random coil is the dominant prediction for PHL4_effector_, which is consistent with our interpretation for the measured R_h_, CD spectra, and Trp fluorescence assays. For PHL4_FL_, the N-terminal effector domain is mostly predicted to be in a random coil arrangement as well, with weak predictions for an α-helix from residues P160–D168. At the C-terminus of PHL4_FL_, in agreement with the secondary chemical shift analysis, the TALOS-N predictions predict a random coil arrangement.

Lastly, NMR relaxation experiments were used to determine the timescales of motion present in PHL4_effector_. Residue-specific longitudinal relaxation (*R_1_*) and longitudinal relaxation in the rotating frame (*R_1ρ_*) measurements were recorded. These data are used to calculate the residue-specific transverse relaxation rate (*R_2_*) (Libich et al., 2015; Palmer, 2004). Figures 5e–f show the *R_1_* and *R_2_* rates with respect to residue number. Supplemental Figures 8a–b shows these relaxation rates as a function of external magnetic field. Supplemental Figure 8c shows heteronuclear ^1^H–^15^N Nuclear Overhauser effect (NOE) enhancements. Relaxation parameters are dependent on both global and local timescales of motion, i.e. the reorientational correlation times, for each residue. Consistent with the lack of both global and local structure in PHL4_effector_, the *R_1_* or *R_2_* values are relatively small across the PHL4_effector_ sequence in Figure 5e–f and there is no strong ^1^H–^15^N NOE enhancement shown in Supplemental Figure 8c. If there were regions transiently adopting a folded conformation, elevated *R_2_* values would be observed. For example, *R_2_* values above 8 s^-1^ were observed for transient helical structure in the TDP-43 protein disordered domain (Conicella et al., 2016). As expected, due to additional freedom for movement for the end-of-chain residues, the N- and C-terminal residues do experience greater molecular motion, i.e. shorter reorientational correlation times producing smaller *R_1_* and *R_2_* values, than residues located in the central portion of the sequence.

Our measurements of intrinsic disorder in the effector domain within PHL4_FL_ are not well predicted by computational algorithms. Supplemental Figure 9 shows the DISOPRED 3.0 (Jones et al., 2015) and Espriv (Walsh et al., 2012) analysis of the PHL4_FL_ sequence do not strongly predict a long continuous disordered domain for the effector domain, instead predicting disorder for shorter segments in the vicinity of residues F175–N225 and the N-terminal region. Supplemental Figure 10 shows the AlphaFold 2.0 (Jumper et al., 2021; Varadi et al., 2022) prediction for PHL4. The prediction for the effector domain is mostly lacking secondary structure and has low confidence scores, consistent with our experimental characterization. Supplemental Figure 10b suggests a slight propensity for helical structure between residues P160–L171, consistent with the weak helical TALOS-N prediction for residues P160– D168, based on our solution NMR assignments. However, residues L161–D168 do not have enhanced relaxation parameters and therefore likely retain significant disorder in our samples. Across DISOPRED, ESPRITZ, and AlphaFold, the C-terminal region of PHL4_FL_ is more confidently predicted to be disordered, in agreement with our solution NMR result for residues G369–E397.

## Discussion

Our measurements indicate the effector domain of PHL4, both in isolation and in the full-length protein, is intrinsically disordered. Intrinsic Trp fluorescence and solution NMR data show that W113 and W149 are solvent accessible and disordered, while W236, conserved in the DNA binding domain of homologues like PHR1, is likely in a well-folded region. W236 is homologous to W230 in the PHR1 homologue, which has been shown to be rigidly folded in a dimeric conformation when bound to DNA from the P1BS promotor (Jiang et al., 2019). Thus, our results are consistent with structure formation in the putative DNA binding region of PHL4.

The solution NMR chemical shift and relaxation measurements are consistent with the effector domain exhibiting significant disorder both in isolation and *within* full-length PHL4. Diffusion NMR and SEC show the effector domain exists as a disordered monomer in solution. The oligomeric state of full-length PHL4 in solution appears to be an even-numbered oligomer from the cross-linking assay, likely between a tetramer and an octamer. Our diffusion NMR and SEC results are supportive of this interpretation but the combination of disordered and ordered segments suggested by CD measurements precludes an exact size determination using these methods. However, the coiled-coil region of the PHR1 homologue was observed to be responsible for tetrameric oligomerization in crystal structures, explaining the binding of phosphate response inhibitors like SPX1 (Jiang et al., 2019; Ried et al., 2021).

In combination with the identification of the disordered effector domain and C-terminal region in full length PHL4 from our multidimensional solution state NMR study, these bulk measurements point to structured oligomerization likely involving the DNA binding and coiled-coil domains of PHL4. The precise nature of the PHL4 oligomer remains to be determined, including the dependence on biological context, such as when interacting with the P1BS DNA binding site or partner proteins like SPX or those involved in cellular transcriptional machinery.

How does an intrinsically disordered effector domain fit in with the current picture of transcriptional activation by eukaryotic specific transcription factors? This question is important in light of results showing that the PHL4 effector domain is a *multi-organism* activating TF (Hummel et al., 2023). It has been suggested that, for mRNA encoding genes in plants, intrinsically disordered regions in transcription factors interact with the mediator complex, facilitating favorable contacts with enhancer sites bound by the DNA-binding domains (Buendía-Monreal et al., 2016). For instance, a recent study showed that the highly acidic repeat DFDLDMLGD in the viral TF VP16 interacts with a component of the native plant mediator complex, Med25 Activation Interaction Domain, potentially explaining the mechanism through which the acidic effector domain exerts its function to increase transcription of mRNA targets (Aguilar et al., 2014). Although the exact VP16 acidic amino acid sequence is not found in the PHL4 effector domain, there is an extreme bias toward aspartic acid residues, with one DDDD repeat and two DDD repeats within the first 35 residues of PHL4. This suggests that PHL4 might attribute some of its gene-activation function through a mechanism similar to the artificial TF VP16, via a highly acidic subregion within its effector domain. It remains to be discovered what accessory transcription components the PHL4 effector domain interact with. Discovery of the interaction network and strengths of interaction of the PHL4 effector domain with the supporting cast of transcriptional machinery could explain why the PHL4 effector domain is more effective than VP16 in activation of transcription for certain genes in some eukaryotes (Hummel et al., 2023).

Acidic regions are more common in activating transcription factors than the general proteome (Kotha et al., 2023; Soto et al., 2022), perhaps in order to make favorable contacts with the preinitiation complex (PIC) to effectively recruit and/or retain PIC binding at a promotor site. In PHL4, the initial acidic region within the N-terminal motif is separated from the predicted DNA binding region by what is shown here to be a long, mostly disordered region. This long, disordered region could provide the necessary conformational flexibility to allow the acidic region of PHL4 to effectively reach the PIC complex. As PHL4 is shown to affect many different genes (Wang et al., 2018) at varying distances from promotor sites like P1BS (Jiang et al. 2019), the disordered effector domain plasticity and dynamics could be a key component to accommodate the different configurations of upregulated genes and the associated promotor regions. One way to visualize this hypothesis is that PHL4 intrinsically disordered effector domain may act like a dynamic extension ladder, able to be shortened and lengthened as needed to increase transcription, whether binding at proximal promotor site or at distant enhancer site.

## Conclusion

PHL4 is a plant TF protein that plays an important role in activating genes in response to low levels of phosphate. In addition, the PHL4 effector domain is a promising genetic tool to up-regulate proteins in multiple organism types. This study provides a structural characterization of PHL4 and its potent effector domain. The 227 amino acid N-terminal effector domain is shown by various techniques to be intrinsically disordered which, as a fragment, is a monomer in solution. This study also demonstrates R_h_ measurements demonstrate that full-length PHL4, although retaining significant disorder in its effector domain and C-terminus, likely forms an oligomer via regions outside of the effector domain. It remains to be shown whether these disordered regions adopt structure when PHL4 binds to DNA or interacts with partner proteins.

## Acknowledgements

Professor Patrick Shih, Niklas Hummel, and Kasey Markel, generously provided the plasmids used in this work, as well as provided insight regarding transcription factor proteins. Dr. Ping Yu and Professor Jim Ames at UC Davis provided technical assistance with solution NMR experiments. Professor David Libich from UT Health San Antonio provided NMR relaxation pulse sequences and provided insight regarding the data analysis. Dr. Upasana Sridharan and Khaled Jami provided technical assistance with the mass spectrometry experiments. The UC Davis NMR Core Facility is partially supported though National Science Foundation award DBI-0722538 and award National Institutes of Health award S10RR013871-01A1. The content is solely the responsibility of the authors and does not necessarily represent the official views of the National Institutes of Health or the National Science Foundation. UC Davis provided financial support for these experiments.

## Materials and Methods

### Protein Expression

The PHL4_FL_ (uniport accession number Q8GXC2) and PHL4_effector_ (N-terminal effector domain, residues 1–227) proteins were expressed with an N-terminal 6xHis tag followed by a TEV cleavage site. These sequences are shown in Supplemental Figure 1. The plasmids, generously provided by the laboratory of Dr. Patrick Shih at UC Berkeley, used pET28 expression vectors containing an inducible lac-operon system and Kanamycin antibiotic resistance. Protein concentrations were quantified using the extinction coefficient calculated using the online ProtParam tool (https://web.expasy.org/protparam/).

Bacterial expression procedures were the same for each construct except where specified. Plasmids were transformed into in-house prepared chemically competent BL21(DE3) E. coli cells using standard heat shock methods, spread onto Luria Broth (LB) agar plates containing 50 µg/mL kanamycin, and incubated overnight at 37 °C. A single isolated colony was used to inoculate 10 mL (PHL4_FL_) or 15 mL (PHL4_effector_), of Luria Broth (LB) media containing 50 µg/mL of kanamycin. The cultures were incubated with shaking at 37 °C and 220 RPM until an optical density at 600 nm (OD_600_) of 1.6–1.7 was reached. Glycerol cell stocks were made with equal parts liquid cell culture and 0.22 µm sterile filtered 50% v/v glycerol in water, inverted gently to mix, flash frozen in liquid nitrogen, and stored at −80 °C.

To start large scale protein expression for unlabeled proteins frozen cells were streaked from the −80 °C glycerol stocks with a sterile 200 µL pipet tip onto LB agar plates containing 50 µg/mL kanamycin and placed into a gravity convection incubator at 37 °C for 16 h. The plate was removed from the incubator and stored at room temperature for ∼3 h until use. Cells from the plate were transferred using a sterile 200 µL pipet tip into a 100–500 mL starter culture, of sterile LB media containing 50 µg/mL kanamycin and 1% w/v glucose added to prevent un-induced protein expression. The culture was incubated at 37 °C in a gravity convection incubator. After incubation for 21 h, 80 mL of the culture is placed into 1 L of sterile LB media containing 1% w/v glucose and 50 µg/mL kanamycin, and incubated with shaking at 37 °C and 220 RPM until the OD_600_ reached 0.7−0.9. Isopropyl-β-D-thiogalactoside (IPTG) was added to 0.5 mM to induce protein expression. After 3 h further growth at 37 °C and 220 RPM, the cells are harvested by centrifugation at 6,000 g for 10 min. The cell pellet was scraped into conical tubes, flash frozen in liquid nitrogen, and stored at −80 °C.

Isotopically labeled protein was expressed identically to the unlabeled protein until the transfer to the 1 L LB media culture. Here, 4 L of LB media split evenly between 4 flasks of 4 L volume containing 50 µg/mL kanamycin and 1% w/v glucose was inoculated with 80 mL of the LB pre-culture per L of LB media. The 1 L cultures were incubated with shaking at 37 °C and 220 RPM until an OD_600_ of 0.7–0.9 was reached. Then the cells were harvested by centrifugation at 6,000 g for 10 m and resuspended in small volumes (∼20 mL) of M9 minimal media using an automatic pipettor and serological pipette, combined, and transferred to a single 1 L of M9 minimal media. The 1 L M9 culture contained 1 g of ^15^N labelled ammonium chloride and 2 g of ^13^C uniformly labelled α-D glucose (Cambridge Isotope Laboratories). The M9 culture was incubated for 30 min with shaking at 37 °C and 220 RPM before adding IPTG to 0.5 mM. After 3 h of additional incubation at 37 °C and 220 RPM, the cells were harvested by centrifugation at 6,000 g for 10 min, scraped into a plastic tube, flash frozen in liquid nitrogen, and stored at −80 °C.

### Purification of Protein Constructs

The cell pellet was thawed on ice for 30 min, then partially resuspended using a serological pipette with ∼25 mL of lysis buffer containing 6 M Gu-HCl, 1% v/v Triton X-100, 500 mM sodium chloride, 50 mM tris(hydroxymethyl)aminomethane (Tris-HCl) pH 7.5, 0.25 mg/mL hen egg white lysozyme (Fisher), and three tablets of EDTA-free Pierce protease inhibitor tablets. The partially resuspended cells were then sonicated in an ice bath with a Branson SFX-250 sonifier pulsed at 0.3 s on, 3 s off, 20 min total “on” time, using a 1/4-inch microtip. The sonicated solution was centrifuged at 4 °C and 75,600 g for 30 min. Protein was purified from the supernatant using a Bio-Rad NGC Quest 10 Plus liquid chromatography instrument and a 5 mL Cytiva HisTrap FF Ni^2+^ immobilized metal affinity chromatography (IMAC) column. The supernatant was loaded onto a column pre-equilibrated in 6 M urea, 20 mM 4-(2-hydroxyethel)-1-piperazineethanesulfonic acid (HEPES) pH 7.5, and 500 mM sodium chloride, then the column was washed with identical buffer containing an additional 20 mM imidazole until the absorbance at 280 nm (A_280_) returned to the baseline value. Then, the protein was eluted with the same buffer containing 200 mM imidazole. Purification buffers initially, for the PHL4_FL_ unlabeled expression that yielded the protein for the SEC measurement as well as the ^15^N labeled PHL4_effector_ in material, also contained 1 mM dithiothreitol, DTT, in the elution buffer. However, DTT was deemed unnecessary for the initial purification steps and therefore DTT was left out in subsequent purifications to no observed detriment, as determined by SDS PAGE gel analyses similar to those in Supplementary Figure 1.

To return the protein to a more native state for each protein preparation, the purified elution fractions were pooled and dialyzed overnight into 20 mM sodium phosphate pH 7.4, 100 mM sodium chloride, 1 mM DTT, so that less than 1% initial buffer remained at the end of the dialysis. The 6xHis purification tag was removed using TEV protease glycerol stocks (25% v/v glycerol, 1 mM EDTA, 5 mM DTT, 250 mM sodium chloride, 10 mM Tris-HCl pH 8.0) prepared in-house. The concentration of TEV in the reaction was changed from preparation to preparation initially, as follows. The PHL4_FL_ sample used in the SEC test used a 1:200 ratio of TEV:protein with an initial TEV stock concentration of 108 µM. The PHL4_FL_ ^13^C- and ^15^N-labeled sample for the NMR assignment experiments used a 1:20 ratio of TEV:protein, with an initial TEV stock concentration of 82 µM. At this point, the reaction was seen to be highly robust at fully cleaving the protein, therefore all subsequent protein preparations used a 1:250 ratio of TEV:protein, with all subsequent TEV stocks at an initial concentration between 6–7 µM.

In the TEV reaction solution, DTT was added to 1 mM initially, and the cleavage reaction was incubated for 19–30 h (PHL4_FL_) and 48 h (PHL4_effector_ domain). Reactions were run until the protein monomer band in an SDS-PAGE gel shifts completely to the tag-cleaved molecular weight. Fresh 1 mM DTT was added after the first 3 h of reaction, and then once every ∼24 h to ensure the TEV protease remained active. Depending on the activity of the TEV stock which is slightly different from prep to prep, this procedure of checking the cleavage reaction before purifying the cleaved protein ensured a minimum amount of uncleaved protein due to differences in activity of the TEV stocks. Supplementary Figure 1 shows SDS-PAGE gels for IMAC purified and TEV-cleaved proteins.

After the cleavage reaction, the solution was passed over a gravity column containing Bio-Rad Nuvia^TM^ IMAC nickel(II) resin equilibrated in 20 mM sodium phosphate pH 7.4, 100 mM sodium chloride, 5 mM imidazole. The initial flow through was collected, and a few CV of the same buffer was used to rinse the column. These protein fractions were stored at 4 °C until used. The protein concentration in all fractions was quantified using absorption at 280 nm with extinction coefficients calculated using the Expasy ProtParam tool (Gasteiger et al., 2005).

### Disorder Predictions

DISOPRED 3.0 was ran using default parameters on the UCL bioinformatics web portal (http://bioinf.cs.ucl.ac.uk/psipred/), accessed July 2023 (Jones and Cozzetto, 2015). ESpriv was ran with standard parameters utilizing the web portal (http://old.protein.bio.unipd.it/espriv/), accessed in December 2023 (Walsh et al., 2012; Walsh et al., 2014).

### Cross-linking Assay

PHL4_FL_ was thawed at room-temperature from −80 °C protein stocks consisting of equal parts TEV-cleaved PHL4 _FL_ from the reverse IMAC purification and 50% v/v glycerol in H_2_O. Then, PHL4 _FL_ was buffer exchanged into 20 mM sodium phosphate pH 7.4 using Amicon-Ultra 0.5 3kDa MWCO centrifuge filters according to the manufacturer’s instructions until the initial buffer remaining is less than 5%. Then, cross-linking reactions were performed with 15.4 µM protein and 0.004–0.0256% glutaraldehyde, with successive concentrations doubling in 20 mM sodium phosphate pH 7.4 buffer at 26 µL total volume, and incubation at 25 °C for 30 min. For glutaraldehyde concentrations above 0.016%, significant aggregation was observed, preventing the samples from entering the SDS-PAGE gels used for analysis. Reactions were quenched by adding 10 µL 4X SDS-PAGE loading buffer, containing 250 mM Tris-HCl pH 6.8, 8% w/v SDS, 0.2% w/v bromophenol blue, and 40% glycerol. 4 µL of 1 M DTT was then added. Samples were heated to 70 °C for 10 min, cooled at room temperature for ∼5 min, then centrifuged at 14,000 g for 15 min, then loaded onto a 4–10% gradient gel and ran with tricine running buffer at 45 mA for ∼20 min.

### Size Exclusion Chromatography

Protein was buffer exchanged and concentrated in an Amicon-Ultra 0.5 mL 3 kDa MWCO centrifuge filter, per manufacturer’s instructions, into SEC buffer (20 mM sodium phosphate pH 6.2, 100 mM sodium chloride, and 1 mM DTT) until the initial buffer remaining was less than 5%. The protein concentration for PHL4 _FL_ was 76.2 µM and the protein concentration for the PHL4_effector_ domain was 96.8 µM. A 200 µL loading loop was pre-loaded with an excess volume of sample, and ran over a Bio-rad ENrich 650 column equilibrated in SEC buffer at a flow rate of 0.75 mL/min. The Bio-Rad gel filtration standards (product #1511901) were resuspended in 0.5 mL of the same SEC buffer, and ran using the same SEC running buffer as PHL4 _FL_, except that a 100 µL loading loop was used. The peak maxima in the chromatogram were determined using in-house Python scripts and the SciPy package “find_peaks” function. The R_h_ values for the standards from the literature (Stetefeld et al., 2016; Talmard et al., 2007; Wilkins et al., 1999) were plotted as a function of the natural logarithm of the retention time and fit to an exponential decay with the SciPy package “curve_fit” function.

### Fluorescence Experiments

Fluorescence experiments all used a 45 µL 10 mm × 10 mm quarv cuvette (Starna). TEV-cleaved and purified PHL4 _FL_ and the PHL4_effector_ domain were diluted to 2.38 µM in either non-denaturing fluorescence buffer (20 mM sodium phosphate pH 7.4, 100 mM sodium chloride, and 1 mM DTT) or denaturing fluorescence buffer (6 M guanidinium-HCl, 20 mM sodium phosphate, 100 mM sodium chloride, and 1 mM DTT). Fluorescence spectra were recorded using a Cary Eclipse fluorescence spectrometer operating in emission mode with excitation fixed at 290 nm and a slit width for both excitation and emission of 5 nm. The excitation was fixed at 290 nm to avoid contribution from Tyr residues so that the resultant emission spectra would be dominated to the greatest possible extent by signal from Trp residues. The spectral range acquired was 300 to 475 nm, scanned at 120 nm/min in 1 nm intervals, with 0.5 s averaging, and the photomultiplier tube (PMT) set to high.

The denaturing condition spectrum in Figure 1 is the average of three scans, with the spectrum normalized so that the maximum intensity was 1.0. The non-denaturing spectrum in Figure 1, is data from one scan with a 5-point window smoothing function applied and normalized such that the maximum intensity was 1.0. An in-house Python script was used to determine λ_max_, using the “curve_fit” function from SciPy to fit a Taylor series expansion for fluorescence intensity to the data

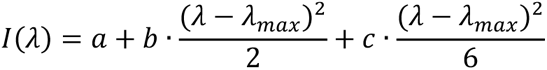

where *I* is the intensity at wavelength λ in nm, *a, b,* and *c* are fitting constants, and λ_max_ is the wavelength of the maximum intensity in the spectra in nm.

### Fluorescence Unfolding Assay

Spectra were recorded of protein diluted to 2.38 µM in a mixture of the non-denaturing and denaturing fluorescence buffers, with final Gu-HCl concentrations from 0 to 6 M; 0, 0.1, 0.2, 0.3, 0.4, 0.5, 0.6, 0.7, 0.8, 0.9, 1.0, 1.1, 1.2, 1.3, 1.4, 1.6, 1.8, 2.0, 2.5, 3.0, 3.5, 4.0, 4.5, 5.0, 5.5, 6.0 M. The samples were allowed to equilibrate overnight at room temperature. Spectra were recoded using a Cary Eclipse fluorescence spectrometer operating in emission mode with excitation fixed at 290 nm and a slit width for both excitation and emission of 5 nm. The excitation was fixed at 290 nm, the acquired spectral range was 300 to 420 nm, scanned at 120 nm/min in 1 nm intervals, with 0.5 s averaging, and the photomultiplier tube (PMT) set to high.

The experimental workflow and equation derivations are described in detail in Monsellier and Bedouelle, 2005, and the analysis here does not deviate substantially. The equations and analysis are restated here for clarity. The overall goal of the procedure is to obtain the mole fraction of unfolded molecular species (*f_U_*) as a function of the Gu-HCl concentration with the observed Gibbs energy of unfolding (ΔG_unfolding_) as a curve-fitting parameter. This analysis assumes a two-state unfolding process, *F* ↔ *U,* and is done via two approaches. In the first approach, the mole fraction of unfolded protein was calculated from λ_max_ using the equation:

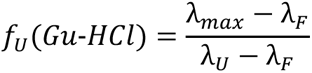

where *f_U_* is the apparent mole fraction of the unfolded molecules for a given Gu-HCl concentration, λ*_F_* and λ*_U_* are the wavelengths of maximum intensity for the 0 M Gu-HCl and 6 M Gu-HCl spectra, and λ*_max_* is the wavelength of maximum intensity for the given concentration of Gu-HCl. λ*_max_* is determined from a third order Taylor series polynomial fit to each spectral curve, according to the equation and procedure given in the *Fluorescence Experiments* section.

In the second approach, the unfolded mole fraction was calculated using the intensity value at the wavelength of maximum intensity difference between the 0 and 6 M spectra, calculated to be 334 nm for this data. A subset of the data, the first five Gu-HCl concentrations for the folded species and the last nine Gu-HCl concentrations for the unfolded species, chosen based on minimal perturbation of the observed spectrum, were independently fit to an equation of the form

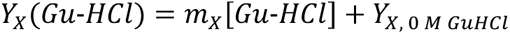

where *Y_X_* is the intensity contribution at 334 nm at each Gu-HCl concentration, [*Gu-HCl*] is the Gu-HCl concentration, *Y_X, 0 M GuHCl_* is the projected intensity at 0 M Gu-HCl, and *m_X_* is the perturbation to the fluorescence intensity at each Gu-HCl concentration. “X” subscripts can be either “_U_” representing the unfolded molecular species or “_F_” representing the folded molecular species. An in-house Python script using the “curve_fit” functionality in SciPy was used to solve for *m_X_* and *Y_x, 0M GuHCl_*.

The measured fluorescence intensity at 334 nm (*Y_meas_*) is then the sum of the contributions from the folded and unfolded molecular species

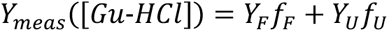

where, *f_F_* is the mole fraction of folded protein and *f_U_* is the mole fraction of unfolded protein. Since the mole fractions of the folded and unfolded molecular species must sum to one, the above equation can be rewritten to obtain the mole fraction of the folded protein as a function of *Y_meas_* and the concentration of Gu-HCl

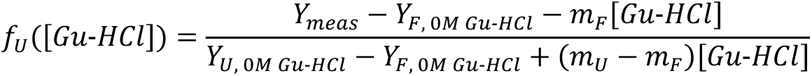

Regardless of the method used to obtain *f_U_*, the Δ*G_unfolding_* is determined from a standard relationship between Δ*G_unfolding_* and the equilibrium constant, *K*, with a linear perturbation due to the Gu-HCl denaturant where ε is a measure of the strength of the perturbation

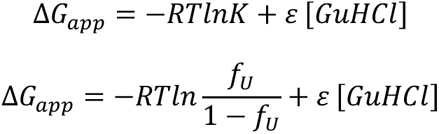

Which can be rearranged to give the fraction of unfolded molecular species as a function of the Gu-HCl denaturant concentration with Δ*G_unfolding_* obtained as a fitting parameter

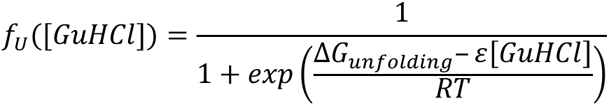

For the mole fractions derived from the first approach, a correction factor more accurately represents Δ*G_unfolding_* due to the non-linearity of the change in λ*_max_* with respect to the protein unfolding process

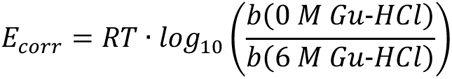

where *b* is the second order constant from the Taylor series fit to the spectrum at the specified Gu-HCl concentration. Supplementary Figure 2 shows the application of these equations to the PHL4_FL_ data.

### Circular Dichroism Spectroscopy

Pure, TEV-cleaved PHL4_effector_ domain and PHL4_FL_ protein was prepared by dialyzing the gravity column flow through fractions into 5 mM sodium phosphate pH 7.4 with 6–8 kDa molecular weight cut-off 0.32 mL/cm dialysis tubing overnight. The protein solutions were then harvested and diluted to 0.3 mg/mL in the same buffer used for the dialysis. The CD spectrum was acquired on a JASCO J-715 CD spectrometer, from 260–190 nm, in 1 nm increments with a 1 mm pathlength cuvette. Three scans were recorded for each sample, and the spectrometer was blanked with the same buffer used for the dialysis prior to running the protein sample. The three scans were averaged in Python prior to plotting. The “Disordered-Ordered Classification” tool from the BESTSEL web portal (Micsonai et al., 2022) was used to identify the general shape of the observed spectrum as disordered (PHL4_effector_) and ordered (PHL4_FL_). BESTSEL was used to determine the structural motifs for PHL4_FL_.

### Mass Spectrometry

Protein for MALDI-MS was prepared by diluting the non-isotopically enriched samples (143 µM PHL4_effector_ and 274 µM PHL4_FL_) from the diffusion NMR experiments into 10 mM Tris-HCl pH 7.5 buffer to a final protein concentration of 5 µM. MALDI-MS was recorded on a Bruker UltraFlextreme MALDI TOF/TOF spectrometer. Bruker Flex Analysis was used to convert the data to text format, then an in-house Python script was used to smooth the spectrum with a 13-point window function, equivalent to smoothing over a 0.5 m/z window. To find the mass spectral peaks, an in-house Python script was used with the “curve_fit” functionality in SciPy.

Protein ESI-MS was prepared by dialyzing 15 µM pure TEV-cleaved protein from the reverse IMAC into 10 mM ammonium carbamate pH 8 with a 6–8 kDa molecular cut-off, 0.32 mL/cm dialysis tubing overnight. The protein was then harvested and diluted to 9 µM with the same buffer used in dialysis. ESI-MS was recorded with a Thermo Q-Exative High-Field Orbitrap ran in full MS mode, from 1200 to 4000 m/z, with 45000 resolution, in positive polarity mode, a spray voltage of 2 kV, a capillary temp of 473 K, and the auxiliary gas heater temperature at 323 K. Spectral analysis was performed first by converting the data with MSConvert (PreoteoWizard 3.0) to UniDec format (Chambers et al., 2012). Then, UniDec was used similar to the method of Marty et al., 2015 to fit the data with 20-point gaussian smoothing, normalized data, and charge search regions of 4–70 (PHL4_effector_) and 4–80 (PHL4_FL_), sampled every 100 Da.

### Diffusion NMR Experiments

Horse Myoglobin protein was purchased from Sigma Aldrich (M1882) and 22.6 mg was resuspended with 700 µL of NMR buffer (20 mM MES pH 6.2 buffer, 25 mM NaCl, and 5 mM TCEP). The tube was gently mixed on a rotary mixer for 1 h and left on the benchtop at ambient temperature for ∼14 h. The Horse Myoglobin NMR sample was then 0.22 µm sterile filtered. The concentration was determined by absorbance at 280 nm and a calculated extinction coefficient to be 3.1 mM. Then 495 µL of the protein sample was mixed with 55 µL 99.8% deuterated water (Sigma-Aldrich). The diffusion NMR NMR samples for the PHL4_FL_ and PHL4_effector_ domain were prepared by concentrating pure non-isotopically labeled, TEV-cleaved protein from the reverse IMAC and buffer exchanging with Amicon Ultra 3 kDa molecular weight cut-off centrifuge filters according to manufacturer recommendations, using 3 to 5 repetitions of diluting the concentrated protein into NMR buffer. The number of repetitions was dependent on ensuring the initial buffer remaining was calculated to be less than 5%. Finally, 55 µL of 99.8% deuterated water (Sigma-Aldrich) was added to 495 µL of each protein solution, giving a final concentration of 246 µM for PHL4_effector_ and 115 µM for PHL4_FL_. All spectra are recorded with 0.1% v/v 1,4 dioxane as an internal diffusion standard.

Spectra were recorded on a 300 MHz Avance Neo NMR spectrometer, using the ledbpgppr2s pulse sequence with water pre-saturation added to reduce baseline distortion from the water resonance (Wu et al., 1995). The bipolar gradient powers range from 2–95% in fixed increments of 6.2% (Wu et al., 1995). For each gradient value, 8 scans were added for PHL4_effector_, and 40 scans were added for PHL4_FL_. The spectrometer frequency was set to the center of the water resonance and the acquisition time was 3.375 s. For PHL4_effector_, the gradient pulse length, δ, was 3200 μs, while for PHL4_FL_, due to slower diffusion, 4000 μs was used. All experiments used a diffusion time, Δ, of 150 ms and an eddy current delay, *T_e_*, of 5 ms. Reference horse myoglobin diffusion NMR spectra were recorded at both 3200 μs and 4000 μs gradient pulse lengths to ensure the calibration was performed with the same experimental parameters for both PHL4_effector_ and PHL4_FL_.

Exponential adiposation functions (1 Hz) were applied and time domain data were Fourier transformed using TopSpin 4.1.4. The spectra were referenced to the 1,4-dioxane resonance to 3.75 ppm. The peak areas were integrated using an in-house Python script and a custom midpoint integral formula for dioxane (3.7525–3.7475 ppm) and the protein (horse myoglobin, 6.0000–9.5000 ppm; PHL4_effector_, 1.0800–0.7500 ppm; PHL4_FL_, 1.0800–0.7500 ppm). The decay of the signals as a function of gradient power were fit using in-house Python scripts utilizing the SciPy “curve_fit” functionality with a Gaussian decay function, as described previously (Wilkins et al., 1999)

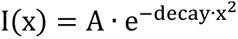

where *I(x)* is the integrated intensity as a function of gradient power *x*, *A* is an amplitude factor, and *decay* describes the decrease in signal with increasing gradient coil powers in the NMR experiment. This decay value was directly used to calculate the radius of hydration. Using dioxane as an internal standard with R_h_ = 2.12 Å, the R_h_ for a protein of unknown size was determined via (Wilkins et al., 1999)

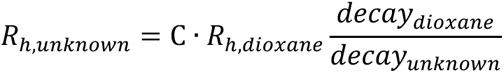

where *C* is an experimentally derived conversion factor. To determine the *C* values, R_h_ was measured in triplicate for horse myoglobin at each gradient length with *C* set to 1. Comparison with the accepted R_h_ value for horse myoglobin (20.4 Å) (Zhou et al., 2002) allows the calculation of the C value for a given gradient length. The R_h_ value measured with a 3200 µs gradient length was 18.5 Å, requiring a C factor of 1.105. For the 4000 µs gradient length the calculated C factor was 1.045. These calibrated C factors were used to calculate the R_h_ values for PHL_effector_ domain and PHL4_FL_ in Table 1.

### NMR Chemical Shift Measurements

NMR experiments were performed on protein in 90% NMR Buffer and 10% deuterated water (99.8%, Sigma-Aldrich). The PHL4_effector_ domain concentration was 73.2 µM for the ^13^C and ^15^N labeled sample and 98.2 µM for the ^15^N labeled sample. The PHL4_FL_ concentration for the ^13^C and ^15^N labeled sample was 114.9 µM.

The NMR samples for PHL4_FL_ and PHL4_effector_ were prepared by concentrating the TEV-cleaved pure protein obtained from the reverse IMAC and buffer exchanging, by 3 to 5 repetitions of concentrating and diluting the protein, into NMR buffer using an Amicon Ultra 3 kDa molecular weight cut-off centrifuge filter according to manufacturer recommendations. The number of repetitions was dependent on ensuring the initial buffer remaining was calculated to be less than 5%. The final NMR sample was obtained by adding 55 µL 99.8% deuterated water (Sigma-Aldrich) to 495 µL of the protein solutions. The data acquisition parameters for the NMR experiments are in Supplemental Tables 1 and 2.

All NMR spectra used for the NMR chemical shift assignments were processed in NMRPipe (Delaglio et al., 1995). Data was converted for peak picking in NMRFAM SPARKY (Lee et al., 2015). Supplementary Tables 1 and 2 contain the data processing parameters.

Sequence specific assignment of the observed NMR chemical shifts for the effector domain were made manually with signal tables consisting of the “*i”* and “*i – 1*” spin clusters resolved in the 3D and 2D spectra according to standard solution NMR assignment strategies. The procedure involves matching the CA and CB NMR chemical shifts from the “*i – 1”* signal table to the same NMR chemical shifts in the “*i“* signal table to determine the “*i“* and “*i – 1”* amide ^15^N and ^1^H NMR chemical shift values. This information is used in combination with the amino acid sequence and the known ranges of NMR chemical shifts for each amino acid type to make unambiguous sequence specific assignment of the observed NMR chemical shifts. Assignments for the PHL4_FL_ C-terminal domain were made according to the same procedure. Due to significant similarity between the spectra of PHL4_effector_ and PHL4_FL_, sequence specific assignment for much of the PHL4_FL_ N-terminal domain were transferable from PHL4_effector_. To make assignments on the PHL4_FL_ N-terminal domain, the peaks observed in PHL4_effector_ spectra are filtered by removing peaks without intensity in the PHL4_FL_ spectra. The assignments were then checked manually. The assigned and unassigned peaks are listed in Supplemental Tables 3 and 4 for PHL4_effector_ and in Supplemental Tables 5 and 6 for PHL4_FL_.

Despite our best effort, including measuring HA and HB NMR chemical shifts for PHL4_effector_ with an additional 3D experiment, the sequence specific assignments were unable to be extended further for PHL4_effector_ and PHL4_FL_. For PHL4_effector_, the inability to extend assignments was primarily due to the high redundancy of the sequence, as some remaining unassigned peaks were of the same signal-to-noise as the assigned peaks but could not be placed on the sequence unambiguously. The N-terminal region of PHL4_effector_ and PHL4_FL_ were especially difficult due to pseudo-repetitive amino acid sequences, unique N and H_N_ but non-unique CA, CB NMR chemical shifts, and in some cases, non-unique HA and HB NMR chemical shifts. Additionally, for PHL4_FL_, sequence specific assignments were also hampered by lower signal-to-noise likely due to the large size of the protein, resulting in less disordered regions of the protein having unfavorable reorientational correlation times for liquid state NMR experiments. This interpretation is also consistent with the smaller number of signals observed in the PHL4_FL_ spectra, and the lack of any signal assignment in the coiled-coil and DNA binding regions of PHL4_FL_.

BMRB accession numbers for the assignments are 52524 (PHL4_FL_) and 52525 (PHL4_effector_).

Secondary chemical shifts are calculated as the difference between the observed chemical shift and the random coil chemical shift from the Poulson IDP calculator. The chemical shift calculation uses a pH of 6.2 and includes nearest neighbor effects (Kjaergaard et al., 2011b). Secondary chemical shift plots were made by plotting the difference of CA and CB secondary chemical shifts (Δ*δ*_*CA*_ − Δ*δ*_*CB*_).

### NMR Relaxation Measurements

For all relaxation experiments, the 93.2 µM ^15^N-only labeled PHL4_effector_ sample was used. Data acquisition and processing parameters are listed in Supplementary Table 1. The data were processed and signal intensities determined by integration using NMRPipe (Delaglio et al., 1995). Only non-overlapped peaks in the ^1^H–^15^N HSQC experiment were integrated and the bounds of integration were +/− 1 point in both spectra dimensions for the NOE experiment and +/− 2 points for the *R_1_*, *R_2_*, and *R_1ρ_* experiment. integrals were fit using the “curve_fit” SciPy function to an equation of the form (Palmer et al., 1991)

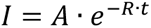

where *A* is an arbitrary scaling factor, *t* is the time in the pulse sequence during which the relaxation process occurred, and *R* is the relaxation time constant (*R_1_*, *R_2_*, or *R_1_*ρ). The uncertainties in Figure 5e–f and Supplementary Figure 8a–b are the standard deviations from the curve-fitting procedure. The ^1^H–^15^N NOE value calculated in Supplementary Figure 8c is the enhancement, defined as (Kharchenko et al., 2020)

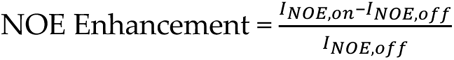

where *I_NOE_* represents the integrated intensity and the “on” and “off” subscript indicating if the ^1^H–^15^N NOE interaction was present during the experiment.

The *R_2_* values in Figure 5e are calculated from *R_1_* and *R_1_*ρ experiments described in Supplementary Table 1 using the equations described in (Lakomek et al., 2012; Libich et al., 2015)

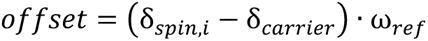

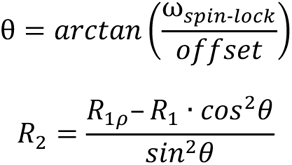

where δ_spin,i_ is the ^15^N chemical shift in ppm units, δ_carrier_ is the ^15^N carrier frequency in ppm units, ω_ref_ is the 0.0 ppm ^15^N chemical shift in MHz units, ω_spin-lock_ was set to 2,000 Hz, and R_1_ρ and R_1_ are the values from the relaxation experiments. ß

