## Supplemental Information for "The Potent PHL4 Transcription Factor Effector Domain Contains Significant Disorder"

### Supplementary Information

(a)

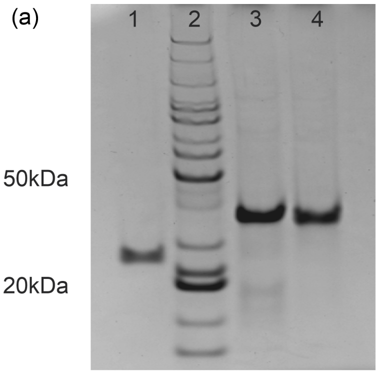

(b)

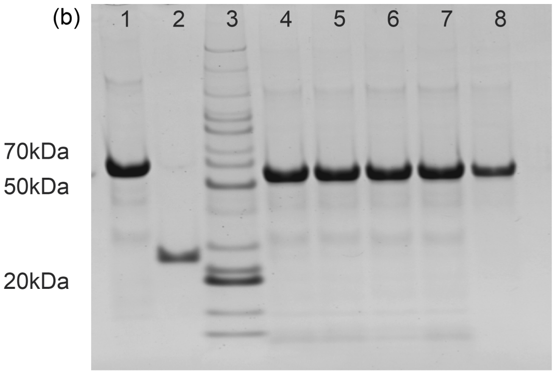

(c)

```
Tag:        MGSSHHHHHHSSGLVPRGSHMENLYFQG

        10         20         30         40
MIPNDDDDAN SMKNYPLNDD DANSMKNYPL NDDDANSMEN
        50         60         70         80
YPLRSIPTEL SHTCSLIPPS LPNPSEAAAD MSFNSELNQI
        90        100        110        120
MARPCDMLPA NGGAVGHNPF LEPGFNCPET TDWIPSPLPH
       130        140        150        160
IYFPSGSPNL IMEDGVIDEI HKQSDLPLWY DDLITTDEDP
       170        180        190        200
LMSSILGDLL LDTNFNSASK VQQPSMQSQI QQPQAVLQQP
       210        220
SSCVELRPLD RTVSSNSNNN SNSNNAA
```

(d)

```
Tag:        MGSSHHHHHHSSGLVPRGSHMENLYFQG

        10         20         30         40
MIPNDDDDAN SMKNYPLNDD DANSMKNYPL NDDDANSMEN
        50         60         70         80
YPLRSIPTEL SHTCSLIPPS LPNPSEAAAD MSFNSELNQI
        90        100        110        120
MARPCDMLPA NGGAVGHNPF LEPGFNCPET TDWIPSPLPH
       130        140        150        160
IYFPSGSPNL IMEDGVIDEI HKQSDLPLWY DDLITTDEDP
       170        180        190        200
LMSSILGDLL LDTNFNSASK VQQPSMQSQI QQPQAVLQQP
       210        220        230        240
SSCVELRPLD RTVSSNSNNN SNSNNAAAAA KGRMRWTPEL
       250        260        270        280
HEVFVDAVNQ LGGSNEATPK GVLKHMKVEG LTIFHVKSHL
       290        300        310        320
QKYRTAKYIP VPSEGSPEAR LTPLEQITSD DTKRGIDITE
       330        340        350        360
TLRIQMEHQK KLHEQLESLR TMQLRIEEQG KALLMMIEKQ
       370        380        390
NMGFGGPEQG EKTSAKTPEN GSEESESPRP KRPRNEE
```

**Supplementary Figure 1. Preparation of PHL4effector and PHL4FL.** (a) Denaturing, non-reducing, 4–10% bis-acrylamide gradient gel ran with tricine running buffer. Lane 1 is the Tobacco Etch Virus protein (TEV) used to cleave off the purification tag. Lane 2 is the protein standard ladder (Invitrogen™ BenchMark™, 10747012), dark bands indicate 50 and 20 kDa molecular weights. Lane 3 is the IMAC purified PHL4effector sample before cleaving with TEV. Lane 4 is the purified PHL4effector used for NMR studies, after incubation with TEV protease and reverse phase nickel affinity chromatography purification. (b) Denaturing, non-reducing, 4–10% bis-acrylamide gradient gel ran with tricine running buffer. Lane 1 is the IMAC PHL4FL purification elution fraction before TEV cleavage. Lane 2 is the TEV stock used to cleave the protein. Lane 3 is the same protein standard ladder used in (a). Lanes 4–7 are fractions taken during the PHL4FL TEV cleavage reaction at 7.5 h, 3.7 h, 27.5 h, and 29.5 h. The downward shift of the dark band in lanes 4–7 to ~56 kDa from ~60 kDa in lane 1 demonstrates that the TEV cleaves the tag off most of the protein. Lane 8 is purified PHL4FL after reverse phase IMAC purification, and shows the removal of the lower MW impurities from the sample. (c) PHL4effector sequence with 6x His-tag and TEV-cleavage site; following cleavage, the only non-native residue left is an N-terminal Gly residue. (d) PHL4FL sequence with 6x His-tag and TEV-cleavage site; following cleavage, the only non-native residue left is an N-terminal Gly residue. In (c) and (d) the coloring follows the cartoon in Figure 1 and the N-terminal purification tag is in green.

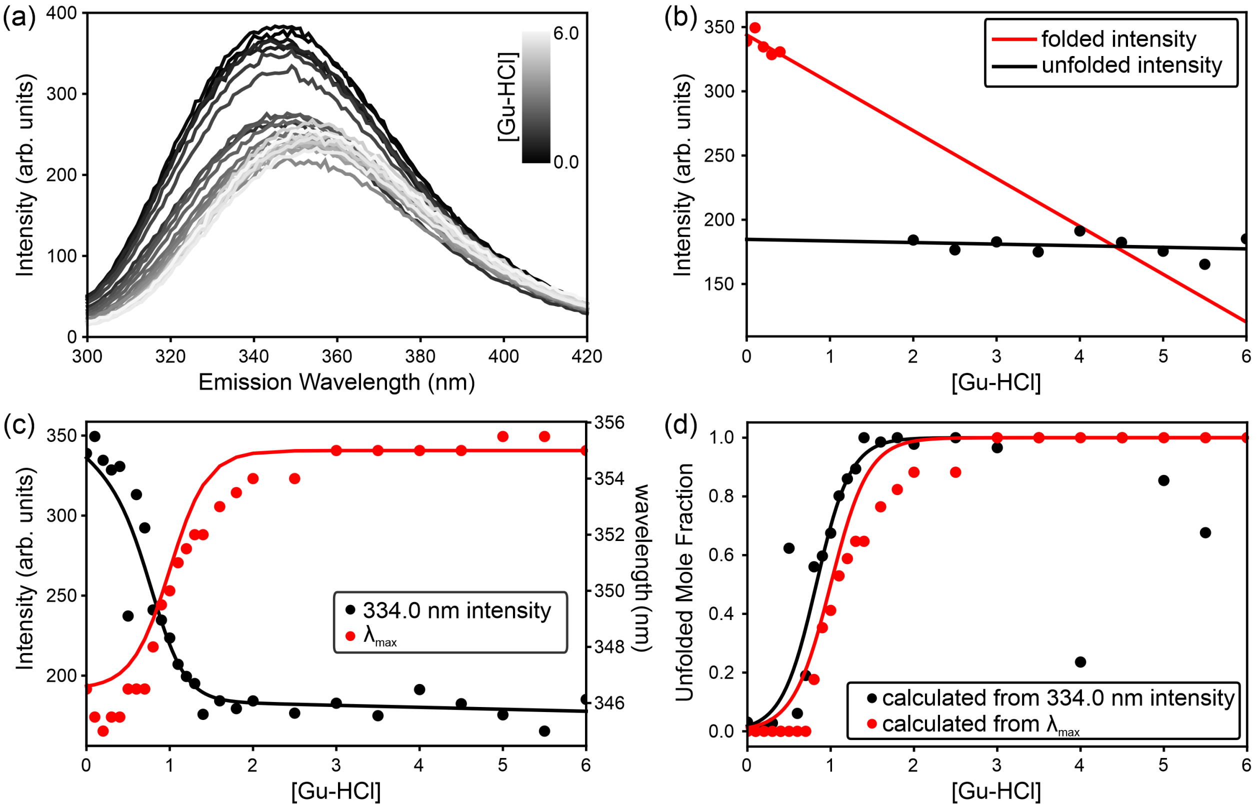

**Supplementary Figure 2. Fluorescence unfolding assay of PHL4.** (a) A series of intrinsic Trp fluorescence spectra of $PHL4_{FL}$ under increasing concentrations of Gu-HCl, with black to grey representing the 0 to 6 M denaturant. The analysis in panels (b)–(d) follows Monsellier and Bedouelle, 2005. (b) Linear plots used to determine the effect of Gu-HCl on the fluorescence intensity independent of the protein unfolding process. (c) The left vertical axis and black line and dots are the fluorescence intensity values at 334 nm versus Gu-HCl concentration. The right vertical axis red line and dots are the wavelength of maximum intensity ($\lambda_{max}$) versus Gu-HCl. (d) Plots of the mole fraction of unfolded $PHL4_{FL}$ versus Gu-HCl concentration, calculated from intensity (black dots) and $\lambda_{max}$ (red dots). The solid lines are a sigmoidal fit to the data, used to obtain the apparent Gibbs energy of folding ($\Delta G_{folding}$). The analysis of the intensity values at 334.0 nm yields a $\Delta G_{folding}$ of –2.5 ± 0.3 kcal/mol and the analysis of the $\lambda_{max}$ yields a $\Delta G_{folding}$ of –2.3 ± 0.8 kcal/mol. As described in the methods section, the slight difference appearance of the sigmoidal curves in (d) does not lead to a large difference in the $\Delta G_{folding}$ fitting parameter due to a correction factor. See the methods section for more details.

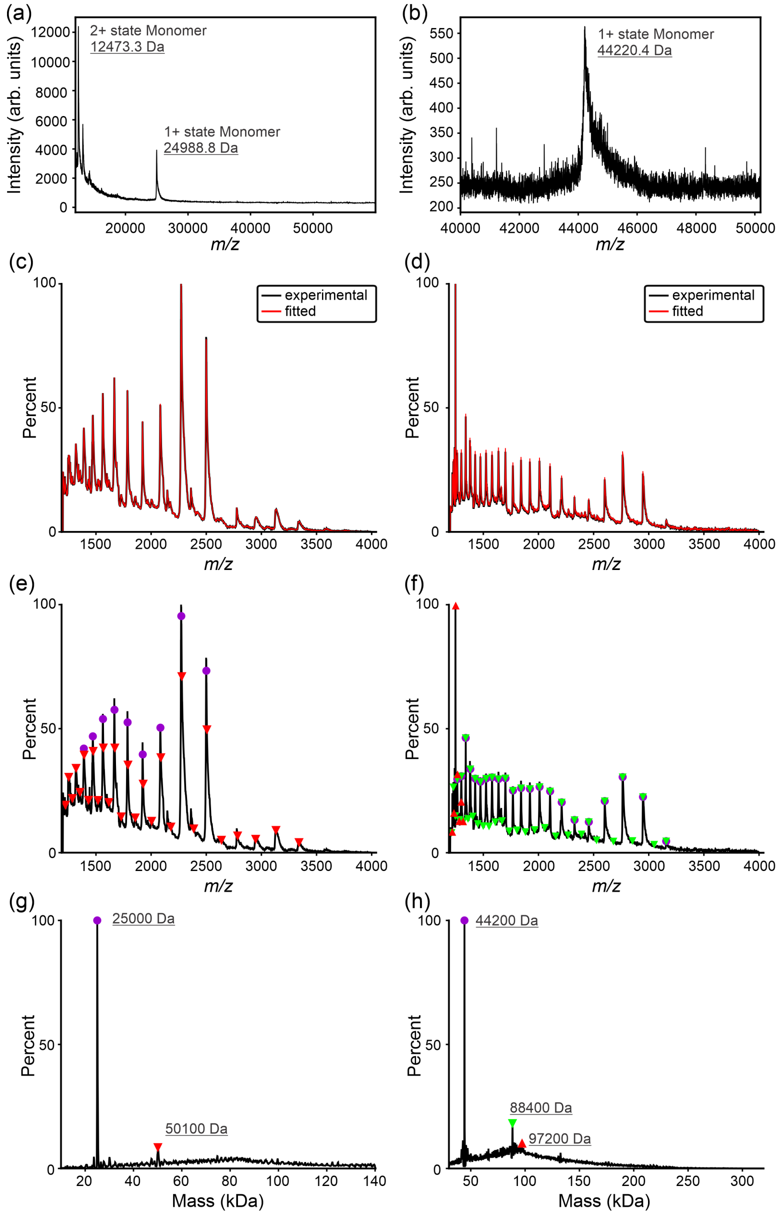

**Supplementary Figure 3. Mass spectrum analysis from MALDI-TOF and ESI-Orbitrap shows primarily monomeric protein.** (a) MALDI-TOF spectrum of the PHL4$_{effector}$ shows mass peaks consistent with the 24963.5 Da mass calculated from the amino acid sequence. (b) MALDI-TOF spectrum of PHL4$_{FL}$ shows a single mass peak consistent with the 44169.3 Da mass calculated from the amino acid sequence. Panels (c), (e), and (g) are an ESI-Orbitrap analysis of the PHL4$_{effector}$. (c) The data calculated from the UniDec program (Marty et al., 2015) in red overlayed on the experimental data in black shows a good fit to the data. Panel (e) shows mass to charge ratio (m/z) peaks derived from the molecular weight species in panel (g) used in the fitting calculation. The symbols in panel (e) correspond to the same molecular species indicated in paned (g). Panels (d), (f), and (h) are an ESI-Orbitrap analysis of PHL4$_{FL}$. The data analysis and plot descriptions are the same as for panels (c), (e), and (g) and also show a good fit to the data.

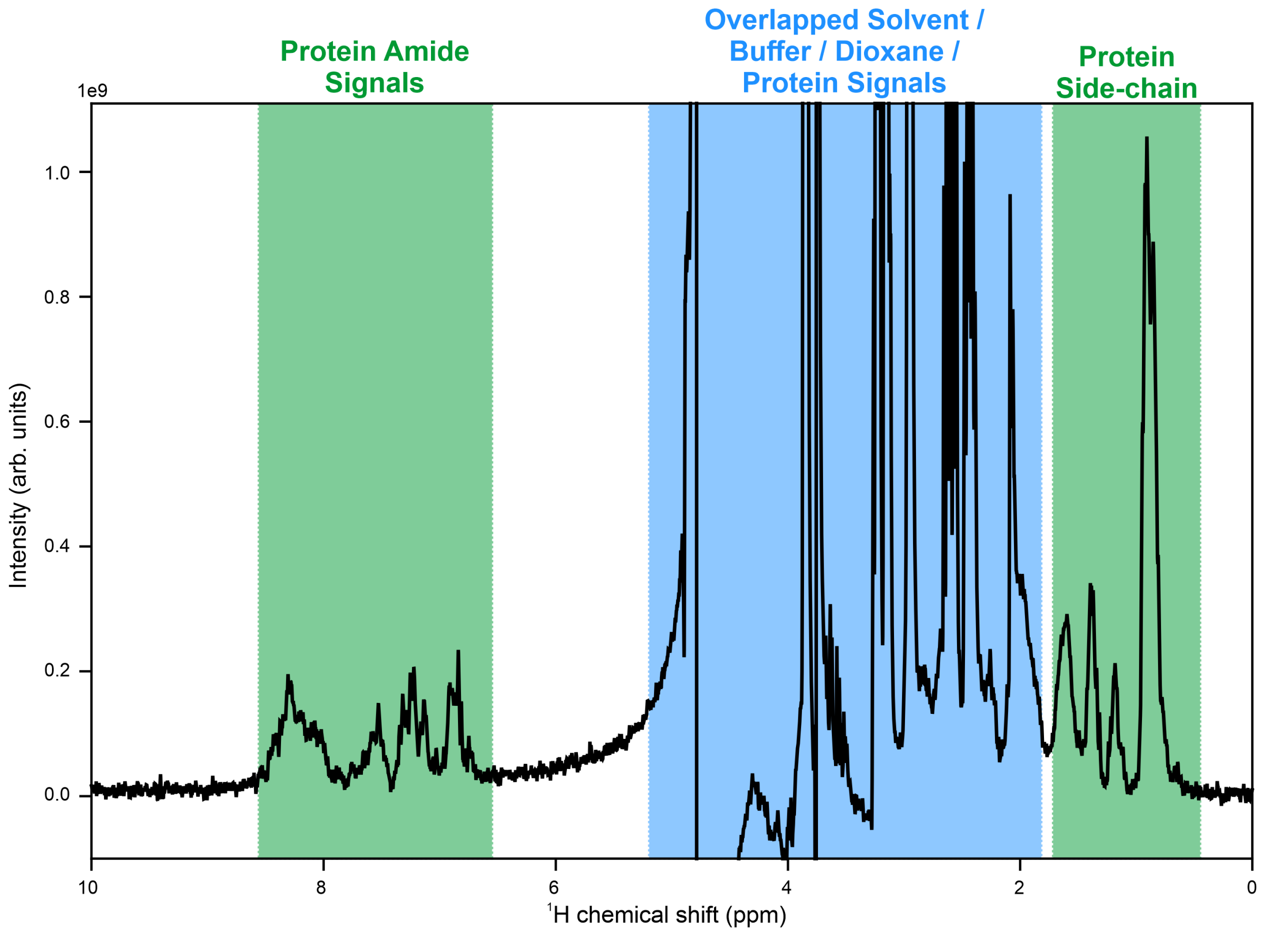

**Supplementary Figure 4. Diffusion NMR spectrum.** 1D $^{1}H$ NMR spectrum of $PHL4_{effector}$ with the weakest applied gradient power from a 2D diffusion experiment at 300 MHz $^{1}H$ frequency. The region labeled "overlapped" contains a mixture of signals from protein, solvent, and other molecules. The protein side chain NMR chemical shifts near 1 ppm are more intense than the amide backbone for the $PHL4_{effector}$, therefore the integrated signal intensity from these side chain peaks is used for the diffusion determination of $R_h$.

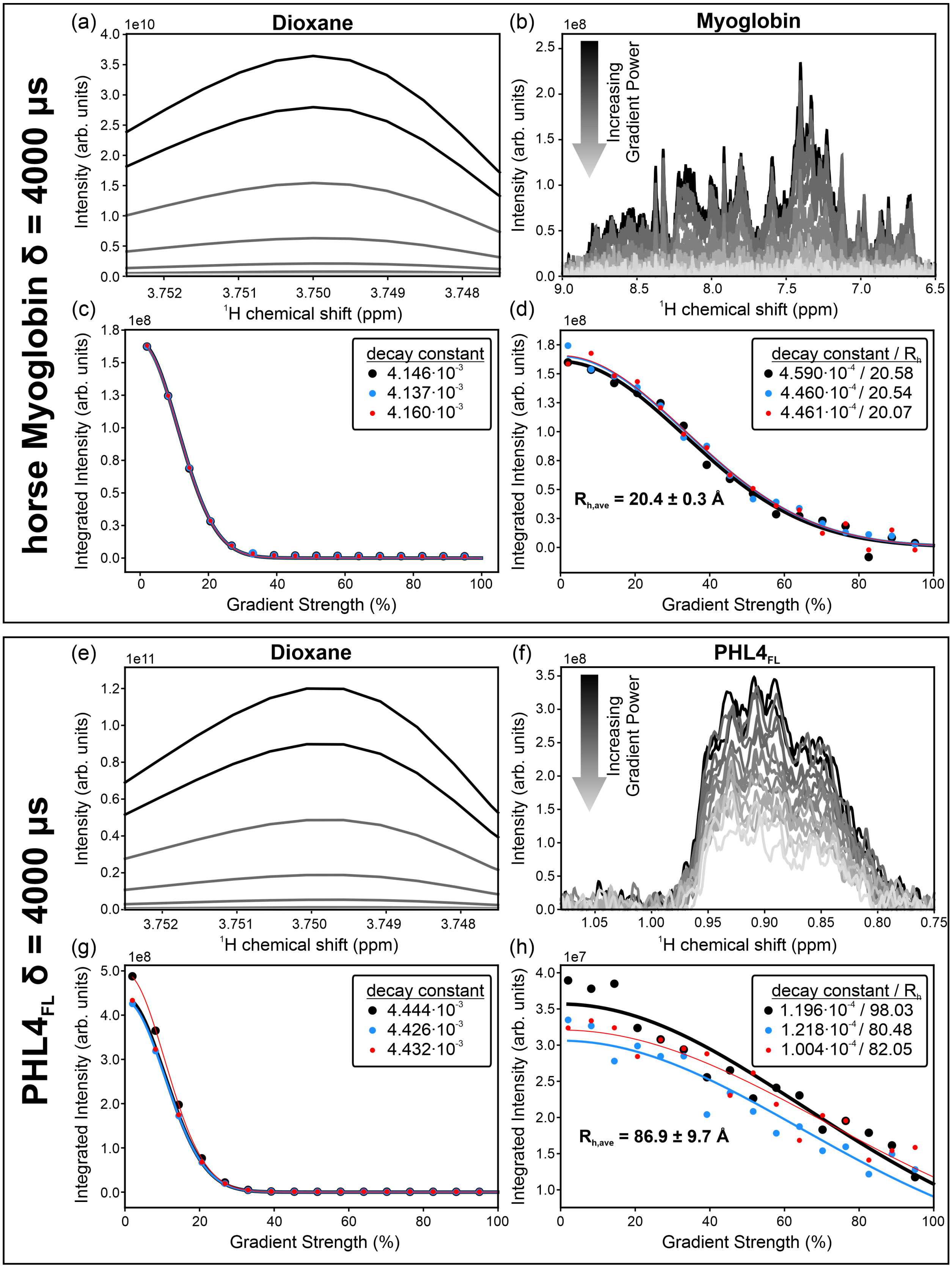
horse Myoglobin δ = 4000 μs
(a)
Dioxane
1e10
Intensity (arb. units)
¹H chemical shift (ppm)
(b)
Myoglobin
1e8
Increasing Gradient Power
Intensity (arb. units)
¹H chemical shift (ppm)
(c)
1e8
decay constant
4.146·10⁻³
4.137·10⁻³
4.160·10⁻³
Integrated Intensity (arb. units)
Gradient Strength (%)
(d)
1e8
decay constant / Rh
4.590·10⁻⁴ / 20.58
4.460·10⁻⁴ / 20.54
4.461·10⁻⁴ / 20.07
Rh,ave = 20.4 ± 0.3 Å
Integrated Intensity (arb. units)
Gradient Strength (%)
PHL4FL δ = 4000 μs
(e)
Dioxane
1e11
Intensity (arb. units)
¹H chemical shift (ppm)
(f)
PHL4FL
1e8
Increasing Gradient Power
Intensity (arb. units)
¹H chemical shift (ppm)
(g)
1e8
decay constant
4.444·10⁻³
4.426·10⁻³
4.432·10⁻³
Integrated Intensity (arb. units)
Gradient Strength (%)
(h)
1e7
decay constant / Rh
1.196·10⁻⁴ / 98.03
1.218·10⁻⁴ / 80.48
1.004·10⁻⁴ / 82.05
Rh,ave = 86.9 ± 9.7 Å
Integrated Intensity (arb. units)
Gradient Strength (%)

**Supplementary Figure 5. Diffusion-NMR measurements for $PHL4_{FL}$ and the horse Myoglobin standard.** Panels (a)–(d) depict the diffusion NMR analysis on the horse Myoglobin standard sample, and (e)–(h) depict the diffusion NMR analysis on the $PHL4_{FL}$ sample. Each sample contains 1,4-dioxane as an internal reference to calibrate diffusion rates. (a) Superposition of spectra from the internal 1,4-dioxane reference covering all 16 1D $^1$H gradient powers for a single diffusion experiment, illustrating the diffusion-based relaxation of the signal due to increasing gradient power linearly in steps of 6.2% from 2% to 95% power. (b) Superposition of spectra from the Horse Myoglobin covering all sixteen 1D $^1$H gradient powers for a single diffusion experiment. For (a) and (b), the black to grey shading represents increasing gradient power. Panels (c) and (d) are plots of integrated intensities as a function of gradient power for three independent measurements. The solid lines are gaussian fits to the integrated intensities. (c) is for the 1,4-dioxane internal reference and (d) is for the horse Myoglobin standard. In panel (d), $R_h$ is the radius of hydration for horse Myoglobin calculated as described in the methods section of this manuscript. $R_{h,ave}$ is the average $R_h$ from the three independent measurements. Panels (e)–(h) are the same as (a)–(d) with horse Myoglobin replaced by $PHL4_{FL}$. In these experiments the diffusion NMR gradient pulse length, $\delta$, is 4.0 ms.

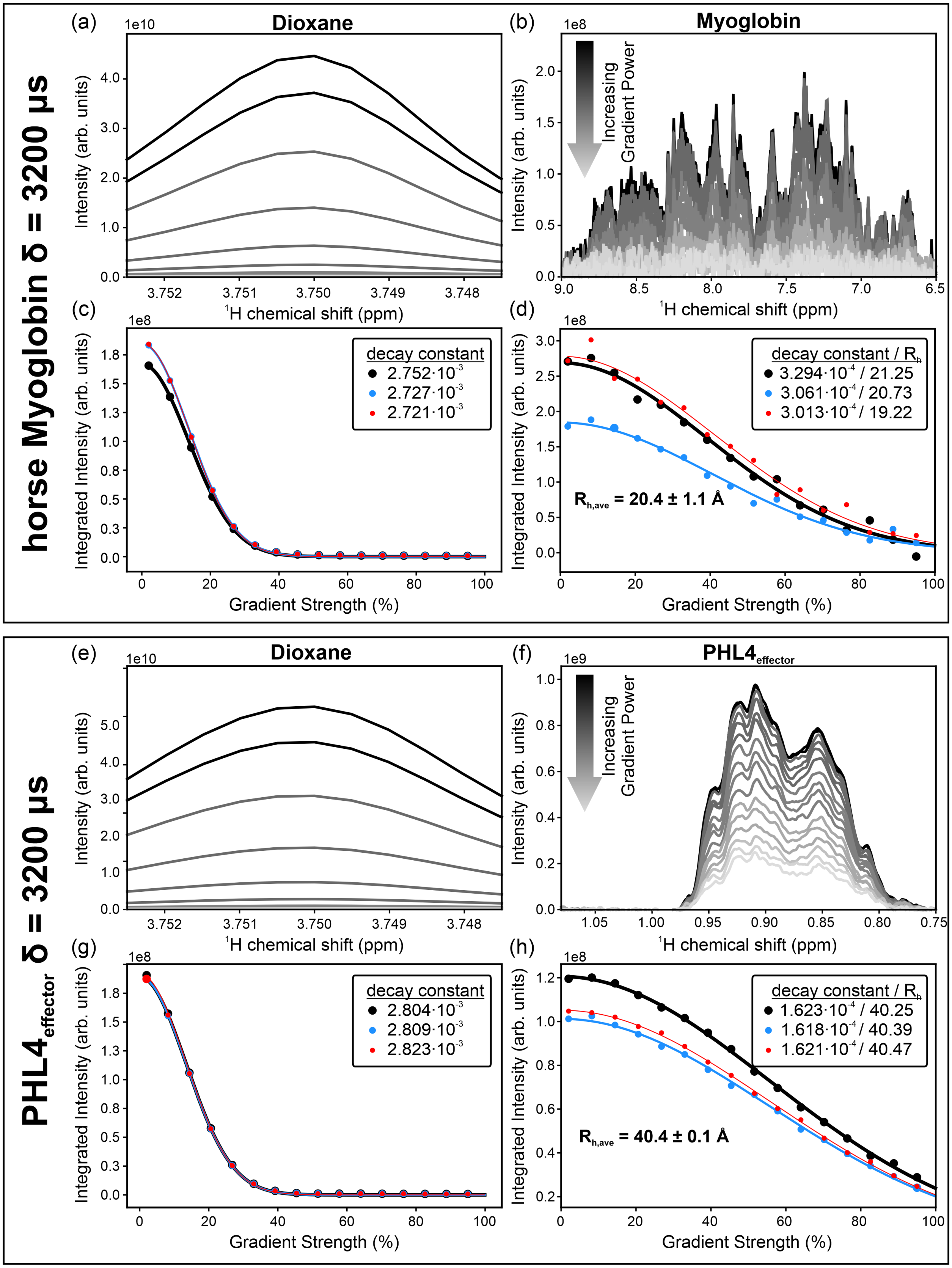
horse Myoglobin δ = 3200 µs
(a)
Dioxane
1e10
Intensity (arb. units)
¹H chemical shift (ppm)
(b)
Myoglobin
1e8
Increasing Gradient Power
Intensity (arb. units)
¹H chemical shift (ppm)
(c)
1e8
Integrated Intensity (arb. units)
Gradient Strength (%)
decay constant
2.752·10⁻³
2.727·10⁻³
2.721·10⁻³
(d)
1e8
Integrated Intensity (arb. units)
Gradient Strength (%)
decay constant / Rh
3.294·10⁻⁴ / 21.25
3.061·10⁻⁴ / 20.73
3.013·10⁻⁴ / 19.22
Rh,ave = 20.4 ± 1.1 Å
PHL4effector δ = 3200 µs
(e)
Dioxane
1e10
Intensity (arb. units)
¹H chemical shift (ppm)
(f)
PHL4effector
1e9
Increasing Gradient Power
Intensity (arb. units)
¹H chemical shift (ppm)
(g)
1e8
Integrated Intensity (arb. units)
Gradient Strength (%)
decay constant
2.804·10⁻³
2.809·10⁻³
2.823·10⁻³
(h)
1e8
Integrated Intensity (arb. units)
Gradient Strength (%)
decay constant / Rh
1.623·10⁻⁴ / 40.25
1.618·10⁻⁴ / 40.39
1.621·10⁻⁴ / 40.47
Rh,ave = 40.4 ± 0.1 Å

**Supplementary Figure 6. Diffusion NMR measurements for $PHL4_{effector}$ and horse Myoglobin standard.** Panels (a)–(d) depict the diffusion NMR analysis on the horse Myoglobin standard sample, and (e)–(h) depict the diffusion NMR analysis on $PHL4_{effector}$ sample. Each sample contains 1,4-dioxane as an internal reference to calibrate diffusion rates. (a) Superposition of spectra from the internal 1,4-dioxane reference covering all 16 1D $^{1}H$ gradient powers for a single diffusion experiment, illustrating the diffusion-based relaxation of the signal due to increasing gradient power linearly in steps of 6.2% from 2% to 95% power. (b) Superposition of spectra from the horse Myoglobin covering all sixteen 1D $^{1}H$ gradient powers for a single diffusion experiment. For (a) and (b), the black to grey shading represents increasing gradient power. Panels (c) and (d) are plots of integrated intensities as a function of gradient power for three independent measurements. The solid lines are gaussian fits to the integrated intensities. (c) is for the 1,4-dioxane internal reference and (d) is for the horse Myoglobin standard. In panel (d), $R_h$ is the radius of hydration for horse Myoglobin calculated as described in the methods section of this manuscript. $R_{h,ave}$ is the average $R_h$ from the three independent measurements. Panels (e)–(h) are the same as (a)–(d) with horse Myoglobin replaced by $PHL4_{effector}$. In these experiments the diffusion NMR gradient pulse length, $\delta$, is 3.2 ms.

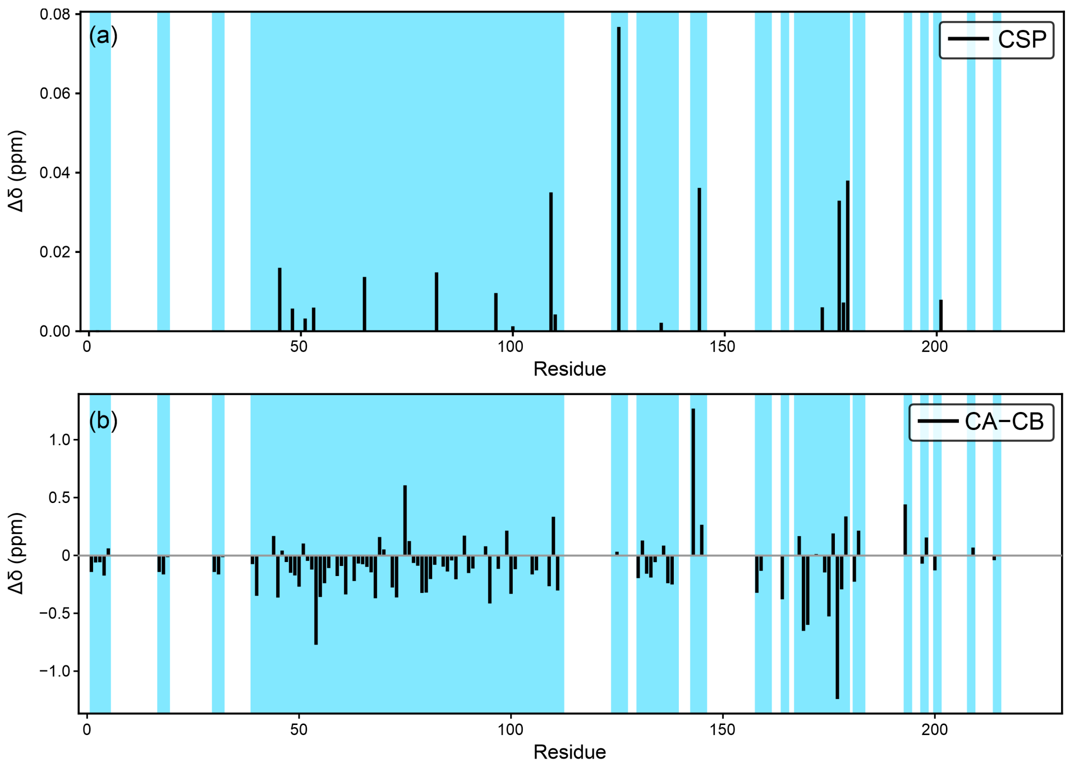

**Supplementary Figure 7. Comparison of $PHL4_{FL}$ and $PHL4_{effector}$ NMR assignments.** (a) $^1H$–$^{15}N$ chemical shift perturbations calculated between $PHL4_{FL}$ and $PHL4_{effector}$ via the formula $CSP = \sqrt{(\Delta\delta_H)^2 + 0.14(\Delta\delta_N)^2}$ (Mureddu et al., 2019), where $\Delta\delta_H$ and $\Delta\delta_N$ represent the ppm difference between the assigned NMR chemical shift values for $^1H$ and $^{15}N$. (b) The difference between $PHL4_{FL}$ and $PHL4_{effector}$ CA and CB NMR chemical shifts for all non-glycine residues where both CA and CB assigned are assigned for both $PHL4_{FL}$ and $PHL4_{effector}$. Light blue shading in both plots indicates sites where chemical shifts are assigned in both $PHL4_{FL}$ and $PHL4_{effector}$, highlighting sites with negligible differences. For specific numbers, compare Supplementary Tables 3 and 5.

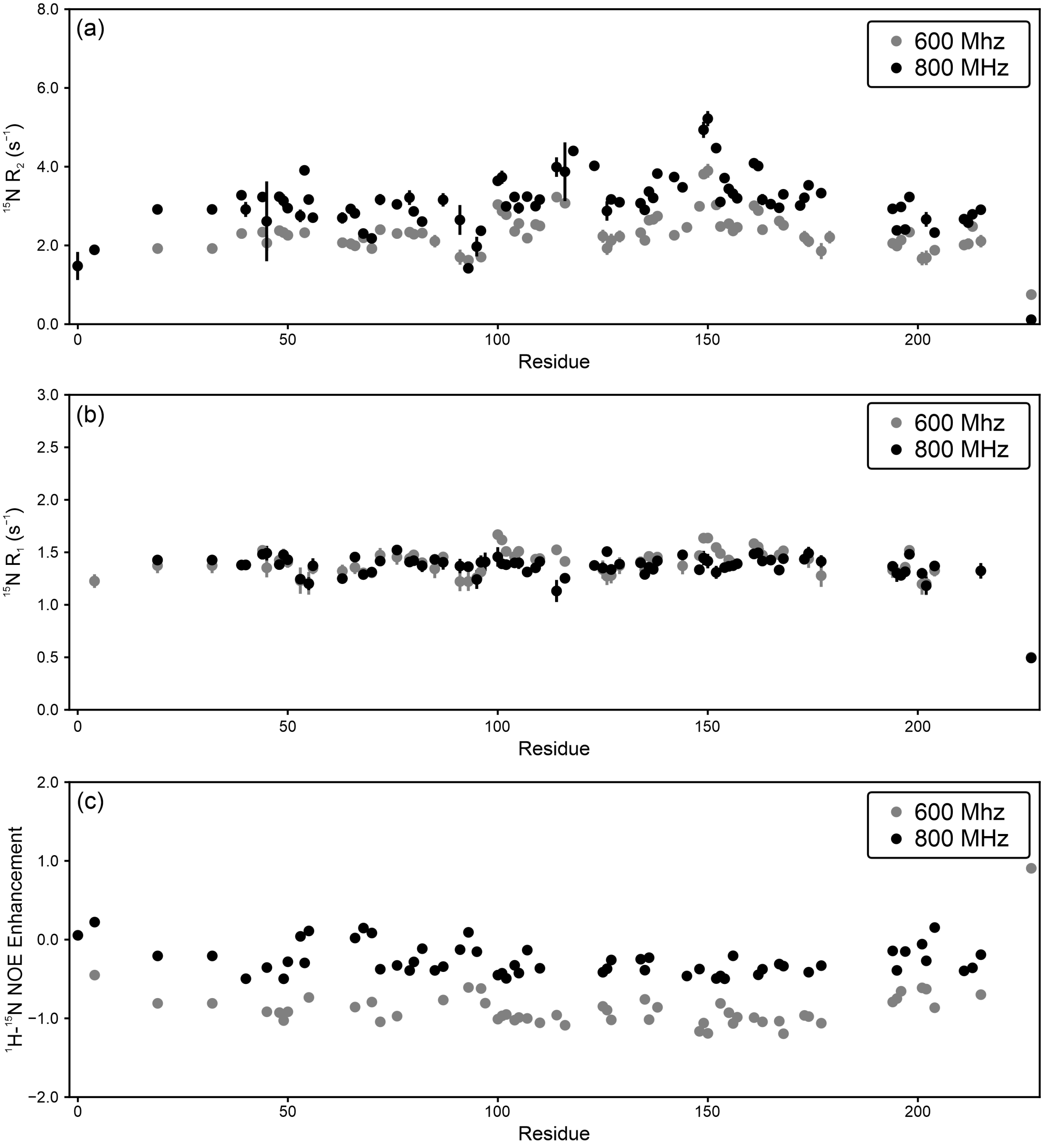

**Supplementary Figure 8. Additional relaxation rates and Nuclear Overhauser Effects (NOE) for PHL4$_{effector}$.** Residue-specific comparison of (a) $^{15}$N R$_2$ relaxation rates, (b) $^{15}$N R$_1$ relaxation rates, and (c) backbone amide $^1$H–$^{15}$H heteronuclear NOE enhancements at 800 MHz and 600 MHz $^1$H NMR frequencies.

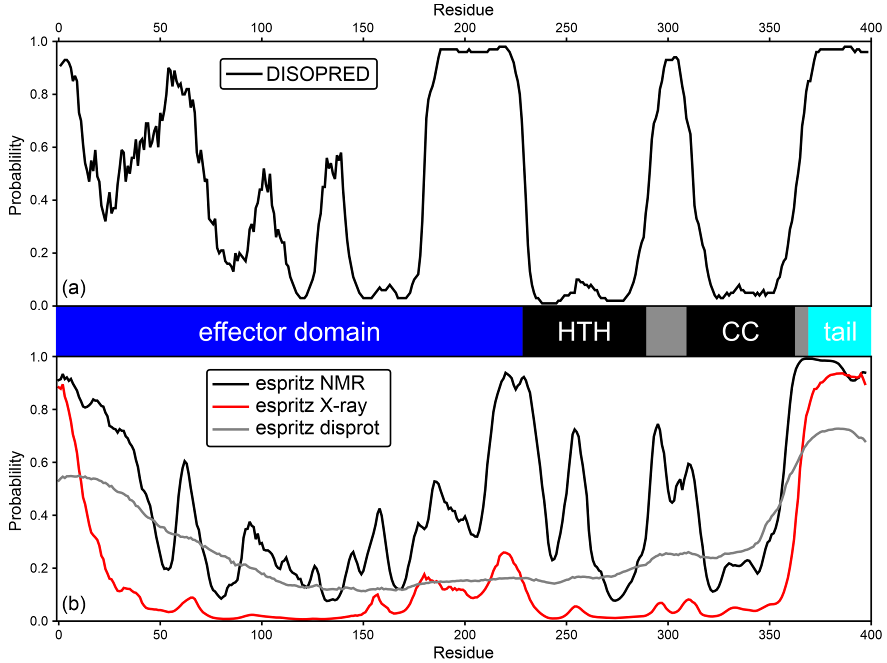

**Supplementary Figure 9. Disorder predictions for PHL4FL.** (a) DISOPRED 3.0 (Jones et al., 2015) prediction for PHL4FL. (b) ESpritz (Walsh et al., 2012; Walsh et al., 2014) predictions of protein disorder for PHL4FL, using the databases listed in the figure legend. For both plots higher probability indicates more confident predictions of disorder. In the cartoon in the middle highlighting the domain structure of PHL4FL, HTH is the helix-turn-helix DNA binding domain and CC is the coiled-coil oligomerization domain.

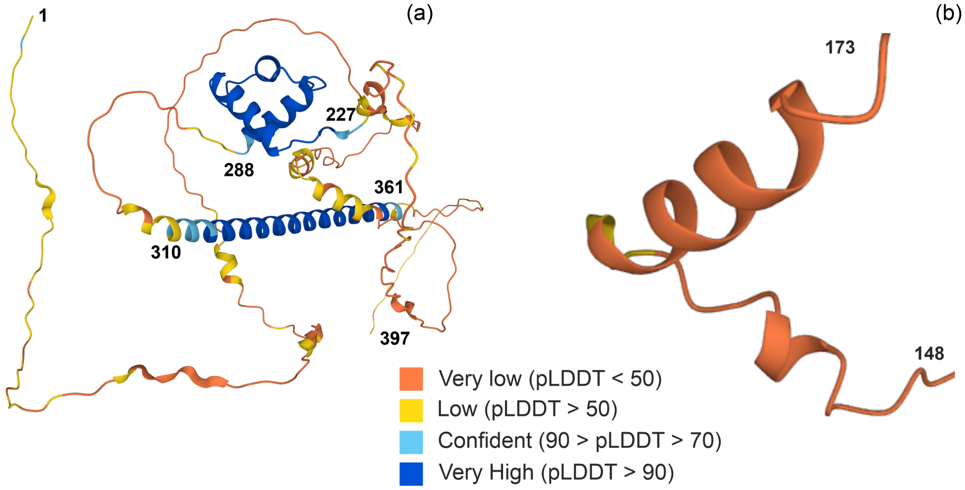

**Supplementary Figure 10. PHL4 structure prediction.** (a) AlphaFold 2.0 prediction for PHL4. The N-terminal effector domain (1–227), the Helix-Turn-Helix DNA binding region (228–288), the coiled-coil domain (310–361), and the C-terminal domain (361–397) are delineated by residues numbers. (b) Expanded view of the low-confidence helical region predicted for residues 160–171 in the Effector Domain. The coloring of the models indicates the confidence of the structure prediction and is described in the figure legend.

**Supplementary Table 1. NMR Experimental Parameters for $PHL4_{effector}$.**

| Spectrum | Acquisition Parameters[a] | Processing Parameters[b] |
|---|---|---|
| **Assignment Experiments** | | |
| 2D $^{1}$H–$^{15}$N HSQC NUS<br>(fhsqcf3gpph) | Spectrometer: 800 MHz Bruker AVANCE III; ns = 32; d1 = 1 s; $\nu_{1H\text{-}carr}$= 4.718 ppm; $\nu_{15N\text{-}carr}$= 117.022 ppm; $\tau_{t1}$ = 90.2 ms; $\tau_{aq}$ = 79.87 ms with 2048 × 512 pts ($^{1}$H, $^{15}$N, indirect dim. w/ States-TPPI); Temp = 296.5 K; 50% NUS, random seed, automatic points | IST NMRPipe processing using ist2D.com, with standard parameters[c].<br>Re-reference in Sparky:<br>$^{15}$N: 0.00 ppm<br>$^{1}$H: 0.061 ppm |
| 2D $^{1}$H–$^{15}$N HSQC<br>(fhsqcf3gpph) | Spectrometer: 800 MHz Bruker AVANCE III; ns = 16; d1 = 1 s; $\nu_{1H\text{-}carr}$= 4.718 ppm; $\nu_{15N\text{-}carr}$= 117.022 ppm; $\tau_{t1}$ = 45.1 ms; $\tau_{aq}$ = 79.87 ms with 2048 × 256 pts ($^{1}$H, $^{15}$N, indirect dim. w/ States-TPPI); Temp = 295.8 K | $SP_{t1}$ -off 0.5 -end 1 -pow 1 -c 1<br>$SP_{t2}$ -off 0.5 -end 1 -pow 1 -c 0.5<br>Re-reference in Sparky:<br>$^{15}$N: –0.07 ppm<br>$^{1}$H: –0.009 ppm |
| 3D CBCACONH<br>(cbcaconhgpwg3d) | Spectrometer: 800 MHz Bruker AVANCE III; ns = 8; d1 = 1 s; $\nu_{1H\text{-}carr}$= 4.788 ppm; $\nu_{15N\text{-}carr}$= 118.093 ppm; $\nu_{13C\text{-}carr}$= 41.75 ppm; $\tau_{t1\,N}$ = 12.33 ms; $\tau_{t2\,C}$ = 4.54 ms; $\tau_{aq}$ = 91.75 ms with 2048 × 60 × 128 pts ($^{1}$H, $^{15}$N, $^{13}$C, indirect dim. w/ States-TPPI); Temp = 296.5 K | SOL<br>$SP_{t1}$ -off 0.5 -end 1 -pow 1 -c 1<br>$SP_{t2}$ -off 0.5 -end 1 -pow 1 -c 0.5<br>$SP_{t3}$ -off 0.5 -end 1 -pow 1 -c 0.5<br>Re-reference in Sparky:<br>$^{15}$N: 0 ppm<br>$^{13}$C: 0 ppm<br>$^{1}$H: 0 ppm |
| 3D CBCANH<br>(hncacbgpwg3d) | Spectrometer: 800 MHz Bruker AVANCE III; ns = 8; d1 = 1 s; $\nu_{1H\text{-}carr}$= 4.788 ppm; $\nu_{15N\text{-}carr}$= 118.093 ppm; $\nu_{13C\text{-}carr}$= 41.75 ppm; $\tau_{t1\,N}$ = 12.3 ms; $\tau_{t2\,C}$ = 4.26 ms; $\tau_{aq}$ = 91.75 ms with 2048 × 60 × 120 pts ($^{1}$H, $^{15}$N, $^{13}$C, indirect dim. w/ States-TPPI); Temp = 296.5 K | SOL<br>$SP_{t1}$ -off 0.5 -end 1 -pow 1 -c 1<br>$SP_{t2}$ -off 0.5 -end 1 -pow 1 -c 0.5<br>$SP_{t3}$ -off 0.5 -end 1 -pow 1 -c 0.5<br>Re-reference in Sparky:<br>$^{15}$N: 0 ppm<br>$^{13}$C: 0 ppm<br>$^{1}$H: 0 ppm |
| 3D HNCO<br>(hncogpwg3d) | Spectrometer: 800 MHz Bruker AVANCE III; ns = 8; d1 = 1 s; $\nu_{1H\text{-}carr}$= 4.788 ppm; $\nu_{15N\text{-}carr}$= 118.089 ppm; $\nu_{13C\text{-}carr}$= 175.724 ppm; $\tau_{t1\,N}$ = 12.33 ms; $\tau_{t2\,C}$ = 22.72 ms; $\tau_{aq}$ = 91.75 ms with 2048 × 60 × 128 pts ($^{1}$H, $^{15}$N, $^{13}$C, indirect dim. w/ States-TPPI); Temp = 296.5 K | SOL<br>$SP_{t1}$ -off 0.5 -end 1 -pow 1 -c 1<br>$SP_{t2}$ -off 0.5 -end 1 -pow 1 -c 0.5<br>POLY -auto<br>$SP_{t3}$ -off 0.5 -end 1 -pow 2 -c 0.5<br>Re-reference in Sparky:<br>$^{15}$N: 0 ppm<br>$^{13}$C: 0 ppm<br>$^{1}$H: 0 ppm |
| 3D HNCA<br>(hncagpwg3d) | Spectrometer: 800 MHz Bruker AVANCE III; ns = 8; d1 = 1 s; $\nu_{1H\text{-}carr}$= 4.788 ppm; $\nu_{15N\text{-}carr}$= 117.089 ppm; $\nu_{13C\text{-}carr}$= 56.751 ppm; $\tau_{t1\,N}$ = 12.33 ms; $\tau_{t2\,C}$ = 10.6 ms; $\tau_{aq}$ = 91.75 ms with 2048 × 60 × 128 pts ($^{1}$H, $^{15}$N, $^{13}$C, indirect dim. w/ States-TPPI); Temp = 296.5 K | SOL<br>$SP_{t1}$ -off 0.5 -end 1 -pow 1 -c 1<br>$SP_{t2}$ -off 0.5 -end 1 -pow 1 -c 0.5<br>$SP_{t3}$ -off 0.5 -end 1 -pow 1 -c 0.5<br>Re-reference in Sparky:<br>$^{15}$N: 0 ppm<br>$^{13}$C: 0 ppm |

| | | |
|---|---|---|
| | | $^{1}H$: 0.0ppm |
| 3D HBHA(CO)NH (hbhaconhgpwg3d) | Spectrometer: 600 MHz Bruker AVANCE III; ns = 4; d1 = 1 s; $\nu_{1H\text{-}carr}$= 4.771 ppm; $\nu_{15N\text{-}carr}$= 117.0 ppm; $\nu_{1H\text{-}carr}$= 3.084 ppm; $\tau_{t1\ N}$ = 16.45 ms; $\tau_{t2\ H}$ = 21 ms; $\tau_{aq}$ = 137.6 ms with 3072 × 60 × 170 pts ($^{1}H$, $^{15}N$, $^{13}C$, indirect dim. w/ States-TPPI); Temp = 298.2 K | SOL<br>$SP_{t1}$ -off 0.5 -end 1 -pow 1 -c 1<br>$SP_{t2}$ -off 0.5 -end 1 -pow 1 -c 0.5<br>$SP_{t3}$ -off 0.5 -end 1 -pow 1 -c 0.5<br>Re-reference in Sparky:<br>$^{15}N$: 0.16ppm<br>$^{1}H$ Direct: 0.004 ppm<br>$^{1}H$: 0.004 ppm |
| 3D HBHANH (hbhanhgpwg3d) | Spectrometer: 600 MHz Bruker AVANCE III; ns = 4; d1 = 1 s; $\nu_{1H\text{-}carr}$= 4.771 ppm; $\nu_{15N\text{-}carr}$= 117.07 ppm; $\nu_{1H\text{-}carr}$= 3.104 ppm; $\tau_{t1\ N}$ = 16.45 ms; $\tau_{t2\ H}$ = 15 ms; $\tau_{aq}$ = 170.4 ms with 3072 × 60 × 120 pts ($^{1}H$, $^{15}N$, $^{13}C$, indirect dim. w/ States-TPPI); Temp = 298.2 K; 50% NUS, random seed, automatic points | NMRPipe IST with ist3d.com and standard parameters[c].<br>Re-reference in Sparky:<br>$^{15}N$: 0.000 ppm<br>$^{1}H$ Direct: 0.011 ppm<br>$^{1}H$: 0.011 ppm |
| **Relaxation Experiments** | | |
| NOE $^{1}H$–$^{15}N$ (hsqcnoef3gpsi) | Spectrometer: 600 MHz Bruker AVANCE III; ns = 10; $\nu_{1H\text{-}carr}$= 4.778 ppm; $\nu_{15N\text{-}carr}$= 117.076 ppm; $\tau_{t1}$ = 45.1 ms; $\tau_{aq}$ = 213 ms with 4096 × 256 pts ($^{1}H$, $^{15}N$, indirect dim. w/ Echo-Antiecho); d1 = 5 s; Temp = 297.5 K | -COADD used to separate $NOE_{on}$/$NOE_{off}$ data<br>-MAC bruk_ranceY.M<br>-$GM_{t1}$ -g1 5 -g2 20 -c 1<br>-$SP_{t2}$ -off 0.5 -end 1 -pow 1 -c 0.5<br>Re-reference in Sparky:<br>$^{15}N$: 0.000 ppm<br>$^{1}H$: 0.004 ppm |
| NOE $^{1}H$–$^{15}N$ (hsqcnoef3gpsi) | Spectrometer: 800 MHz Bruker AVANCE III; ns = 56; $\nu_{1H\text{-}carr}$= 4.718 ppm; $\nu_{15N\text{-}carr}$= 117.02 ppm; $\tau_{t1}$ = 45.1 ms; $\tau_{aq}$ = 159.7 ms with 4096 × 256 pts ($^{1}H$, $^{15}N$, indirect dim. w/ Echo-Antiecho); d1 = 5 s; Temp = 296.5 K | -COADD used to separate NOE/NONOE data<br>-MAC bruk_ranceY.M<br>-$GM_{t1}$ -g1 5 -g2 10 -c 1<br>-$SP_{t2}$ -off 0.5 -end 1 -pow 1 -c 0.5<br>Re-reference in Sparky:<br>$^{15}N$: 0.000 ppm<br>$^{1}H$: 0.067 ppm |
| $^{15}N$ $T_1$ (hsqc1etf3gpsi3d) | Spectrometer: 600 MHz Bruker AVANCE III; ns = 16; $\nu_{1H\text{-}carr}$= 4.778 ppm; $\nu_{15N\text{-}carr}$= 117.084 ppm; $\tau_{t1}$ = 45.1 ms; $\tau_{aq}$ = 106.5 ms with 2048 × 256 pts ($^{1}H$, $^{15}N$, indirect dim. w/ Echo-Antiecho); d1 = 1 s; Temp = 297.5 K; $T_1$ delay = 0.1 ,0.2 ,0.3 ,0.4 ,0.6 ,0.8 ,1.0 s | TP, ZTP, TP used to rearrange the data for FT.<br>$SP_{t1}$ -off 0.5 -end 1 -pow 1 -c 1<br>$SP_{t2}$ -off -0.5 -end 1 -pow 2 -c 1<br>Re-reference in Sparky:<br>$^{15}N$: –0.08 ppm<br>$^{1}H$: 0.005 ppm |
| $^{15}N$ $T_1$ (hsqc1etf3gpsi3d) | Spectrometer: 800 MHz Bruker AVANCE III; ns = 8; $\nu_{1H\text{-}carr}$= 4.794 ppm; $\nu_{15N\text{-}carr}$= 117.1 ppm; $\tau_{t1}$ = 31.65 ms; $\tau_{aq}$ = 91.75 ms with 2048 × 154 pts ($^{1}H$, $^{15}N$, indirect dim. w/ Echo-Antiecho); d1 = 1 s; Temp = 295.8 K; $T_1$ delay = 0.1, 0.2, 0.3, 0.4, 0.6, 0.8, 1.0 s | TP, ZTP, TP used to rearrange the data for FT.<br>$SP_{t1}$ -off 0.5 -end 1 -pow 1 -c 1<br>$SP_{t2}$ -off -0.5 -end 1 -pow 2 -c 1<br>Re-reference in Sparky:<br>$^{15}N$: –0.09 ppm<br>$^{1}H$: 0.01 ppm |

| | | |
|---|---|---|
| $^{15}$N $T_2$ (hsqc2etf3gpsi3d) | Spectrometer: 600 MHz Bruker AVANCE III; ns = 16; $\nu_{1H\text{-}carr}$= 4.778 ppm; $\nu_{15N\text{-}carr}$= 117.084 ppm; $\tau_{t1}$ = 45.1 ms; $\tau_{aq}$ = 106.5 ms with 2048 × 256 pts ($^1$H, $^{15}$N, indirect dim. w/ Echo-Antiecho); d1 = 1 s; Temp = 297.5 K; $T_2$ loop counter = 0, 9, 27, 46, 64, 91, 119; d31=0.01696 s | TP, ZTP, TP used to rearrange the data for FT.<br>$SP_{t1}$ -off 0.5 -end 1 -pow 1 -c 1<br>$SP_{t2}$ -off -0.5 -end 1 -pow 2 -c 1<br>Re-reference in Sparky:<br>$^{15}$N: –0.07 ppm<br>$^1$H: 0.004 ppm |
| $^{15}$N-$T_{1rho}$ with sensitivity enhanced (Rance-Kay) read-out (Lakomek et al., 2012) | Spectrometer: 800 MHz Bruker AVANCE III; ns = 16; $\nu_{1H\text{-}carr}$= 4.788 ppm; $\nu_{15N\text{-}carr}$= 117.092 ppm; $\tau_{t1}$ = 45.06 ms; $\tau_{aq}$ = 79.87 ms with 2048 × 256 pts ($^1$H, $^{15}$N, indirect dim. w/ Echo-Antiecho); d1 = 3 s; Temp = 296.5 K; spin-lock $T_{1rho}$ period = 0.071, 0.001, 0.125, 0.21, 0.101, 0.041 s; 2 kHz spin-lock pulse | Custom c script to separate the interleaved 2D planes before nmrPipe processing.<br>In fid.com, -yMODE Echo-Antiecho<br>$SP_{t1}$ -off 0.5 -end 1 -pow 1 -c 1<br>$SP_{t2}$ -off -0.5 -end 1 -pow 2 -c 0.5<br>Re-reference in Sparky:<br>$^{15}$N: –0.11 ppm<br>$^1$H: –0.007 ppm |
| $^{15}$N-$T_1$ with sensitivity enhanced (Rance-Kay) read-out (Lakomek, 2012) | Spectrometer: 800 MHz Bruker AVANCE III; ns = 16; $\nu_{1H\text{-}carr}$= 4.788 ppm; $\nu_{15N\text{-}carr}$= 117.092 ppm; $\tau_{t1}$ = 45.06 ms; $\tau_{aq}$ = 79.87 ms with 2048 × 256 pts ($^1$H, $^{15}$N, indirect dim. w/ Echo-Antiecho); d1 = 3 s, Temp = 296.5 K; $T_1$ period = 0.04, 1, 0.24, 0.68, 0.12, 0.4 s; Heat compensation spin-lock pulse, 2 kHz, 71 ms | Custom c script to separate the interleaved 2D planes before nmrPipe processing.<br>In fid.com, -yMODE Complex -MAC bruk_ranceY.M<br>$SP_{t1}$ -off 0.5 -end 1 -pow 1 -c 1<br>$SP_{t2}$ -off -0.5 -end 1 -pow 2 -c 0.5<br>Re-reference in Sparky:<br>$^{15}$N: –0.095ppm<br>$^1$H: –0.008ppm |

[a] ns is the number of scans averaged; $\tau_{aq}$ is the total acquisition time in the directly detected dimension; $\nu_{X\text{-}carr}$ is the center frequency for nucleus x; $\tau_{tX}$ is the total acquisition time for the indirect dimension x. The proton pulse length was optimized and used from the value given by the bruker "pulsecal" command with a 1d proton experiment, zggpwg.

[b] Spectra are processed with NMRPipe (Delaglio et al., 1995). Sinebell apodization was applied according to the parameters specified, and Gaussian line broadening is applied where specified. Zero filling was applied twice before applying each Fourier transform. All spectra were indirectly referenced by utilizing the H2O peak and the D2O lock signal, with respect to a calibrated temperature value in NMRPipe in the 'bruker' GUI. After NMRPipe processing, sparky re-referencing was applied align the spectra to the peaks in $PHL4_{effector}$ HSQC (Lee et al., 2015). See methods for more information.

[c] (Delaglio et al., 1995; Hyberts et al., 2012)

**Supplementary Table 2. NMR Experimental Parameters for PHL4.**

| Spectrum | Acquisition Parameters[a] | Processing Parameters[b] |
|---|---|---|

| | | |
|---|---|---|
| 2D $^{1}$H–$^{15}$N HSQC<br>(fhsqcf3gpph) | Spectrometer: 600 MHz Bruker AVANCE III; ns = 16; d1 = 1 s; $\nu_{1H\text{-}carr}$= 4.7 ppm; $\nu_{15N\text{-}carr}$= 118 ppm; $\tau_{t1}$ = 65.8 ms; $\tau_{aq}$ = 113.6 ms with 2048 × 256 pts ($^{1}$H, $^{15}$N, indirect dim. w/ States-TPPI); Temp = 297.5 K | $SP_{t1}$ -off 0.5 -end 1 -pow 1 -c 1<br>$SP_{t2}$ -off 0.5 -end 1 -pow 1 -c 0.5<br>Re-reference in Sparky:<br>$^{15}$N: 0.08 ppm<br>$^{1}$H: 0.082 ppm |
| 3D CBCACONH<br>(cbcaconhgpwg3d) | Spectrometer: 600 MHz Bruker AVANCE III; ns = 8; d1 = 1 s; $\nu_{1H\text{-}carr}$= 4.7 ppm; $\nu_{15N\text{-}carr}$= 118.007 ppm; $\nu_{13C\text{-}carr}$= 41.663 ppm; $\tau_{t1\ N}$ = 16.5 ms; $\tau_{t2\ C}$ = 6.06 ms; $\tau_{aq}$ = 11.36 ms with 2048 × 64 x 128 pts ($^{1}$H, $^{15}$N, $^{13}$C, indirect dim. w/ States-TPPI); Temp = 297.5 K | SOL<br>$SP_{t1}$ -off 0.5 -end 1 -pow 1 -c 1<br>$SP_{t2}$ -off 0.5 -end 1 -pow 1 -c 0.5<br>$SP_{t3}$ -off 0.5 -end 1 -pow 2 -c 0.5<br>CS -ls 1ppm (due to cnst)<br>POLY -auto<br>Re-reference in Sparky:<br>$^{15}$N: 0.09 ppm<br>$^{13}$C: –0.05 ppm<br>$^{1}$H: 0.088 ppm |
| 3D CBCANH<br>(hncacbgpwg3d) | Spectrometer: 600 MHz Bruker AVANCE III; ns = 8; d1 = 1 s; $\nu_{1H\text{-}carr}$= 4.7 ppm; $\nu_{15N\text{-}carr}$= 118.007 ppm; $\nu_{13C\text{-}carr}$= 41.663 ppm; $\tau_{t1\ N}$ = 16.5 ms; $\tau_{t2\ C}$ = 6.06 ms; $\tau_{aq}$ = 113.6 ms with 2048 × 64 × 128 pts ($^{1}$H, $^{15}$N, $^{13}$C, indirect dim. w/ States-TPPI); Temp = 297.5 K | SOL<br>$SP_{t1}$ -off 0.5 -end 1 -pow 1 -c 1<br>$SP_{t2}$ -off 0.5 -end 1 -pow 1 -c 0.5<br>POLY -auto<br>$SP_{t3}$ -off 0.5 -end 1 -pow 2 -c 0.5<br>Re-reference in Sparky:<br>$^{15}$N: 0.03 ppm<br>$^{13}$C: 0.05 ppm<br>$^{1}$H: 0.088 ppm |
| 3D HNCO<br>(hncogpwg3d) | Spectrometer: 600 MHz Bruker AVANCE III; ns = 8; d1 = 1 s; $\nu_{1H\text{-}carr}$= 4.7 ppm; $\nu_{15N\text{-}carr}$= 118.007 ppm; $\nu_{13C\text{-}carr}$= 177.634 ppm; $\tau_{t1\ N}$ = 16.5 ms; $\tau_{t2\ C}$ = 19.3 ms; $\tau_{aq}$ = 113.6 ms with 2048 × 64 × 128 pts ($^{1}$H, $^{15}$N, $^{13}$C, indirect dim. w/ States-TPPI); Temp = 297.5 K | SOL<br>$SP_{t1}$ -off 0.5 -end 1 -pow 1 -c 1<br>$SP_{t2}$ -off 0.5 -end 1 -pow 1 -c 0.5<br>POLY -auto<br>$SP_{t3}$ -off 0.5 -end 1 -pow 2 -c 0.5<br>Re-reference in Sparky:<br>$^{15}$N: 0.15 ppm<br>$^{13}$C: 0.1 ppm<br>$^{1}$H: 0.088 ppm |
| 3D HNCA<br>(hncagpwg3d) | Spectrometer: 600 MHz Bruker AVANCE III; ns = 16; d1 = 1 s; $\nu_{1H\text{-}carr}$= 4.771 ppm; $\nu_{15N\text{-}carr}$= 117.077 ppm; $\nu_{13C\text{-}carr}$= 55.933 ppm; $\tau_{t1\ N}$ = 11.28 ms; $\tau_{t2\ C}$ = 14.14 ms; $\tau_{aq}$ = 121.65 ms with 2048 × 48 × 128 pts ($^{1}$H, $^{15}$N, $^{13}$C, indirect dim. w/ States-TPPI); Temp = 298.2 K | SOL<br>$SP_{t1}$ -off 0.5 -end 1 -pow 1 -c 1<br>$SP_{t2}$ -off 0.5 -end 1 -pow 1 -c 0.5<br>POLY -auto<br>$SP_{t3}$ -off 0.5 -end 1 -pow 1 -c 0.5<br>Re-reference in Sparky:<br>$^{15}$N: 0 ppm<br>$^{13}$C: 0 ppm<br>$^{1}$H: 0.011 ppm |

[a] ns is the number of scans averaged; $\tau_{aq}$ is the total acquisition time in the directly detected dimension; $\nu_{X\text{-}carr}$ is the center frequency for nucleus x; $\tau_{tX}$ is the total acquisition time for the indirect dimension x. The proton pulse length was optimized and used from the value given by the bruker "pulsecal" command with a 1d proton experiment, zggpwg.

[b] Spectra are processed with NMRPipe (Delaglio, et al., 1995). Sinebell apodization was applied according to the parameters specified, and Gaussian line broadening is applied where specified. Zero filling was applied twice before applying each Fourier transform. All spectra were indirectly referenced by utilizing the H2O peak and the D2O lock signal, with respect to a calibrated temperature value in NMRPipe in the 'bruker' GUI. After NMRPipe processing, sparky re-referencing was applied align the spectra to the peaks in the effector domain HSQC (Lee, et al., 2015). See methods for more information.

**Supplementary Table 3. $PHL4_{effector}$ domain NMR chemical shift assignments.** Column 1 is the residue number, SN represents the peak intensity calculated by NMRFAM-SPARKY (Lee, et al., 2015) for the non-NUS $^{1}$H-$^{15}$N HSQC for $PHL4_{effector}$. For the SN Type, "G" stands for non-overlapped peaks in the HSQC spectrum, whereas "B" represents overlapping peaks in the HSQC spectrum. The assigned NMR chemical shift values come from the HNCACB, HNCA, HN(CO)CACB, HNCO, HNHAHB, HN(CO)HAHB, and NUS-HSQC experiments. In cases where overlapping peaks were observed, the values are taken from the spectrum without any peak overlap. Peaks without N or $H_N$ shifts, but having other nuclei with assigned chemical shifts, come from experiments reporting on the chemical shifts for "*i* – *1*" residues. Black bars represent long amino acid sequences without assignments.

| | AA Type | CA | CB | CO | N | $H_N$ | HA | HB | HB2 | SN | SN Type* |
|---|---|---|---|---|---|---|---|---|---|---|---|
| 0 | G | 45.258 | | | 108.758 | 8.416 | 3.95 | | | 27 | G |
| 1 | M | 55.929 | 33.189 | 175.485 | | | 4.472 | 2.029 | | | |
| 2 | I | 58.625 | 38.651 | | 123.537 | 8.307 | | | | 293 | B |
| 3 | P | 63.328 | 32.212 | 174.949 | | | 4.389 | 2.285 | 1.928 | | |
| 4 | N | 53.341 | 39.199 | 176.421 | 119.125 | 8.541 | 4.672 | 2.808 | | 269 | G |
| 5 | D | 54.649 | 41.104 | | 121.04 | 8.349 | 4.602 | 2.703 | | 397 | B |
| 6 | D | | | | | | | | | | |
| 15 | Y | | | | | | | | | | |
| 16 | P | 63.323 | 32.108 | 174.381 | | | | | | | |
| 17 | L | 55.377 | 42.345 | 174.318 | 121.999 | 8.363 | 4.388 | 1.62 | | 482 | B |
| 18 | N | 53.213 | 39.289 | 176.544 | 119.276 | 8.424 | 4.727 | 2.812 | 2.778 | 520 | B |
| 19 | D | 54.669 | 41.052 | | 121.195 | 8.386 | 4.605 | 2.682 | | 534 | G |
| 20 | D | | | | | | | | | | |
| 28 | Y | | | | | | | | | | |
| 29 | P | 63.323 | 32.108 | 174.381 | | | | | | | |
| 30 | L | 55.377 | 42.345 | 174.318 | 121.999 | 8.363 | 4.388 | 1.62 | | 482 | B |
| 31 | N | 53.213 | 39.289 | 176.544 | 119.276 | 8.424 | 4.727 | 2.812 | 2.778 | 520 | B |
| 32 | D | 54.669 | 41.052 | | 121.195 | 8.386 | 4.605 | 2.682 | | 534 | G |
| 33 | D | | | | | | | | | | |

| | | | | | | | | | | | |
|---|---|---|---|---|---|---|---|---|---|---|---|
| 37 | S | | | | | | | | | | |
| 38 | M | 55.065 | 32.995 | 175.405 | | | 4.525 | | | | |
| 39 | E | 56.801 | 30.323 | 175.344 | 122.578 | 8.461 | 4.291 | 2.1 | 1.954 | 299 | G |
| 40 | N | 53.361 | 38.986 | 177.253 | 118.523 | 8.316 | 4.672 | 2.704 | 2.633 | 64 | G |
| 41 | Y | 56.063 | 38.346 | | 121.724 | 8.025 | 4.657 | 2.876 | | 464 | B |
| 42 | P | | | | | | | | | | |
| 43 | L | 55.316 | 42.03 | 173.988 | | | 4.314 | | | | |
| 44 | R | 56.017 | 30.861 | 175.389 | 121.34 | 8.201 | 4.36 | 1.848 | | 141 | G |
| 45 | S | 58.157 | 63.862 | 177.544 | 116.968 | 8.25 | 4.471 | 3.81 | | 160 | G |
| 46 | I | 58.815 | 38.642 | | 124.233 | 8.174 | | | | 213 | B |
| 47 | P | 63.378 | 32.232 | 174.27 | | | 4.47 | 2.323 | 1.929 | | |
| 48 | T | 62.498 | 69.754 | 176.558 | 114.899 | 8.212 | 4.264 | 4.221 | | 227 | G |
| 49 | E | 56.957 | 30.072 | 174.981 | 123.107 | 8.524 | 4.311 | 2.1 | | 292 | G |
| 50 | L | 55.491 | 42.351 | 173.982 | 122.998 | 8.228 | 4.25 | 1.605 | | 269 | G |
| 51 | S | 58.579 | 63.786 | 176.917 | 115.8 | 8.183 | 4.397 | 3.881 | | 249 | B |
| 52 | H | 55.872 | 29.437 | 176.538 | 121.999 | 8.363 | | 3.323 | | 482 | B |
| 53 | T | 62.157 | 69.814 | 174.974 | 115.191 | 8.182 | | | | 113 | G |
| 54 | C | 58.657 | 28.643 | 177.037 | 121.676 | 8.425 | 4.671 | | | 283 | G |
| 55 | S | 58.566 | 63.888 | 177.347 | 118.595 | 8.423 | | | | 123 | G |
| 56 | L | 55.289 | 42.353 | 174.643 | 124.476 | 8.263 | | | | 217 | G |
| 57 | I | 58.517 | 38.61 | | 123.975 | 8.137 | 4.285 | | | 269 | B |
| 58 | P | | | | | | | | | | |
| 59 | P | 62.942 | 32.185 | 174.651 | | | | | | | |
| 60 | S | 57.993 | 63.829 | 177.41 | 116.059 | 8.308 | 4.427 | 3.87 | | 520 | B |
| 61 | L | 53.072 | 42.064 | | 125.427 | 8.285 | 4.342 | | | 274 | B |
| 62 | P | 63.153 | 32.106 | 175.092 | | | | | | | |
| 63 | N | 51.409 | 38.71 | | 120.024 | 8.533 | 4.94 | 2.818 | | 223 | G |
| 64 | P | 63.854 | 32.158 | 174.171 | | | 4.408 | 2.311 | | | |
| 65 | S | 58.968 | 63.595 | 176.665 | 115.134 | 8.311 | 4.388 | 3.887 | | 246 | G |
| 66 | E | 56.78 | 30.244 | 175.212 | 122.507 | 8.194 | 4.282 | 2.086 | 1.949 | 270 | G |
| 67 | A | 52.578 | 19.201 | 173.759 | 124.742 | 8.168 | 4.316 | 1.385 | | 481 | B |
| 68 | A | 52.889 | 19.234 | 173.943 | 123.163 | 8.202 | 4.284 | 1.392 | | 232 | G |
| 69 | A | 52.659 | 19.121 | 173.924 | 123.25 | 8.174 | 4.282 | 1.389 | | 255 | B |
| 70 | D | 54.487 | 41.067 | 174.89 | 119.083 | 8.223 | 4.568 | 2.716 | 2.646 | 348 | G |
| 71 | M | 55.691 | 32.574 | 174.763 | 121.381 | 8.29 | 4.446 | | | 273 | B |
| 72 | S | 59.09 | 63.719 | 177.078 | 115.705 | 8.141 | 4.401 | 3.89 | | 241 | G |
| 73 | F | 58.356 | 39.453 | 176.085 | 121.818 | 8.14 | 4.593 | 3.099 | | 321 | B |
| 74 | N | 53.179 | 39.039 | | 120.863 | 8.232 | 4.584 | | | 305 | B |
| 75 | S | 58.386 | 63.719 | 176.448 | | | 4.424 | 3.876 | | | |
| 76 | E | 57.342 | 29.612 | 174.45 | 122.298 | 8.439 | 4.26 | 2.07 | 1.941 | 421 | G |

| | | | | | | | | | | | |
|---|---|---|---|---|---|---|---|---|---|---|---|
| 77 | L | 55.776 | 42.191 | 174.082 | 121.062 | 7.977 | 4.249 | 1.595 | | 252 | B |
| 78 | N | 53.552 | 38.858 | 176.224 | 118.277 | 8.211 | 4.645 | 2.716 | | 448 | B |
| 79 | Q | 56.213 | 29.539 | 175.555 | 119.761 | 8.127 | 4.312 | | | 241 | G |
| 80 | I | 61.255 | 38.615 | 175.471 | 121.617 | 8.09 | 4.212 | 1.854 | | 319 | G |
| 81 | M | 55.182 | 33.08 | 175.862 | 124.919 | 8.413 | | | | 239 | G |
| 82 | A | 52.457 | 19.365 | 174.3 | 125.304 | 8.19 | | | | 183 | G |
| 83 | R | 53.966 | 30.275 | | 121.552 | 8.352 | | | | 226 | B |
| 84 | P | 63.766 | 32.069 | 174.197 | | | | | | | |
| 85 | C | 58.62 | 27.954 | 177.069 | 117.834 | 8.374 | | | | 169 | G |
| 86 | D | 54.638 | 41.051 | 175.562 | 122.545 | 8.291 | | | | 562 | B |
| 87 | M | 55.35 | 32.988 | 175.719 | 119.766 | 8.049 | | | | 218 | G |
| 88 | L | 53.307 | 41.742 | | 124.61 | 8.197 | | | | 253 | B |
| 89 | P | 62.621 | 32.119 | 174.518 | | | 4.389 | | | | |
| 90 | A | 52.86 | 19.034 | 173.623 | 124.364 | 8.439 | 4.261 | 1.374 | | 202 | B |
| 91 | N | 53.184 | 38.918 | 175.596 | 117.305 | 8.411 | 4.692 | 2.834 | | 179 | G |
| 92 | G | 45.761 | | 176.77 | 109.025 | 8.355 | 3.943 | | | 106 | B |
| 93 | G | 45.261 | | 177.738 | 108.378 | 8.221 | 3.937 | | | 152 | G |
| 94 | A | 52.461 | 19.326 | 173.587 | 123.724 | 8.136 | 4.359 | 1.36 | | 185 | B |
| 95 | V | 62.715 | 32.621 | 174.726 | 119.02 | 8.068 | 4.086 | 2.061 | | 58 | G |
| 96 | G | 45.138 | | 177.831 | 112.161 | 8.431 | 3.911 | | | 149 | G |
| 97 | H | 55.524 | 29.823 | 177.497 | 118.33 | 8.243 | 4.476 | 3.169 | | 122 | G |
| 98 | N | 51.013 | 38.942 | | 121.924 | 8.507 | | | | 111 | B |
| 99 | P | 63.584 | 31.953 | 174.906 | | | 4.283 | 2.07 | | | |
| 100 | F | 57.952 | 38.874 | 175.925 | 118.281 | 8.001 | | | | 167 | G |
| 101 | L | 54.662 | 42.72 | 174.907 | 121.858 | 7.68 | | | | 114 | G |
| 102 | E | 54.442 | 29.827 | | 122.834 | 8.095 | | | | 228 | G |
| 103 | P | 63.709 | 32.01 | 173.857 | | | | | | | |
| 104 | G | 45.268 | | 177.471 | 109.024 | 8.462 | 3.905 | | | 163 | G |
| 105 | F | 58.182 | 39.609 | 176.095 | 120.045 | 7.987 | 4.567 | 2.708 | 2.632 | 246 | G |
| 106 | N | 53.103 | 39.178 | 177.356 | 120.744 | 8.327 | 4.569 | 2.675 | | 199 | B |
| 107 | C | 56.608 | 27.495 | | 120.843 | 8.182 | | | | 166 | G |
| 108 | P | 63.465 | 32.106 | 174.642 | | | | | | | |
| 109 | E | 57.122 | 30.056 | 174.707 | 120.447 | 8.536 | 4.306 | | | 275 | G |
| 110 | T | 61.627 | 69.896 | 176.821 | 114.554 | 8.127 | 4.39 | | | 154 | G |
| 111 | T | 61.971 | 69.795 | 177.197 | 115.751 | 8.168 | 4.397 | 4.177 | | 213 | B |
| 112 | D | 55.089 | 40.987 | 176.783 | 122.486 | 8.291 | | | | 562 | B |
| 113 | W | 57.682 | 29.654 | 176.109 | 119.975 | 7.796 | | | | 22 | G |
| 114 | I | 58.137 | 39.248 | | 125.128 | 7.759 | | | | 98 | G |
| 115 | P | 62.875 | 32.065 | 174.91 | | | 4.152 | 2.208 | | | |
| 116 | S | 56.327 | 63.667 | | 117.295 | 8.216 | | | | 150 | G |
| 117 | P | 63.029 | 32.079 | 174.832 | | | | | | | |

| | | | | | | | | | | | |
|---|---|---|---|---|---|---|---|---|---|---|---|
| 118 | L | 53.024 | 41.691 | | 123.506 | 8.211 | | | | 148 | G |
| 119 | P | 62.971 | 32.149 | 174.963 | | | | | | | |
| 120 | H | 55.56 | 29.652 | 177.269 | 118.869 | 8.436 | | | | 97 | B |
| 121 | I | 60.938 | 38.948 | 176.391 | 122.219 | 7.952 | | | | 99 | B |
| 122 | Y | 57.606 | 39.194 | 176.967 | 124.661 | 8.205 | 4.57 | 2.714 | | 159 | B |
| 123 | F | 55.256 | 39.499 | | 123.719 | 8.156 | | | | 132 | G |
| 124 | P | 63.229 | 31.957 | 174.518 | | | 4.367 | 2.274 | 1.958 | | |
| 125 | S | 58.676 | 63.974 | 176.323 | 115.637 | 8.36 | 4.422 | 3.901 | | 165 | G |
| 126 | G | 45.187 | | 177.682 | 110.807 | 8.395 | 4.003 | | | 138 | G |
| 127 | S | 56.444 | 63.729 | | 116.62 | 8.148 | 4.741 | 3.841 | | 188 | G |
| 128 | P | 63.389 | 32.006 | 175.01 | | | 4.365 | 2.207 | 1.881 | | |
| 129 | N | 53.315 | 38.706 | | 117.996 | 8.323 | 4.601 | 2.641 | | 200 | G |
| 130 | L | 55.988 | 42.25 | | | | | | | | |
| 131 | I | 61.612 | 38.417 | 175.122 | 121.474 | 8.115 | 4.291 | | | 499 | B |
| 132 | M | 55.696 | 33.034 | 175.463 | 123.762 | 8.306 | 4.497 | 2.103 | | 196 | B |
| 133 | E | 56.962 | 30.225 | 175.382 | 120.66 | 8.26 | 4.193 | 2.039 | 1.909 | 334 | B |
| 134 | D | 54.713 | 41.159 | 174.714 | 121.386 | 8.417 | 4.597 | 2.688 | | 321 | G |
| 135 | G | 45.574 | | 177.459 | 109.028 | 8.328 | 3.943 | | | 238 | G |
| 136 | V | 62.719 | 32.647 | 175.226 | 120.104 | 7.922 | 4.095 | 2.059 | | 287 | G |
| 137 | I | 61.166 | 38.784 | 175.673 | 124.971 | 8.267 | 4.17 | 1.844 | | 233 | G |
| 138 | D | 54.381 | 41.338 | 175.402 | 124.674 | 8.357 | 4.606 | 2.7 | | 256 | G |
| 139 | E | 56.986 | 30.187 | 174.751 | 121.484 | 8.338 | 4.213 | 2.051 | 1.93 | 404 | B |
| 140 | I | 61.776 | 38.871 | | 120.583 | 8.09 | | | | 508 | B |
| 141 | H | 55.498 | 28.885 | 176.964 | | | 4.693 | 3.273 | 3.133 | | |
| 142 | K | 56.728 | 33.246 | 174.893 | 122.729 | 8.237 | 4.31 | 1.81 | 1.749 | 123 | G |
| 143 | Q | 56.221 | 29.427 | 174.939 | 122.015 | 8.495 | 4.31 | 2.027 | | 238 | B |
| 144 | S | 58.499 | 63.863 | 177.37 | 116.62 | 8.321 | 4.419 | 3.87 | | 467 | G |
| 145 | D | 54.37 | 40.927 | 175.732 | 122.348 | 8.346 | 4.624 | 2.696 | 2.609 | 243 | G |
| 146 | L | 53.166 | 41.607 | | 122.983 | 7.995 | 4.583 | | | 249 | B |
| 147 | P | 62.85 | 31.772 | 174.798 | | | 4.306 | 2.06 | | | |
| 148 | L | 55.533 | 42.31 | 174.449 | 121.818 | 8.178 | 4.216 | 1.491 | | 232 | G |
| 149 | W | 57.05 | 29.649 | 175.927 | 119.666 | 7.755 | 4.62 | 3.218 | | 168 | G |
| 150 | Y | 57.825 | 39.031 | 177.025 | 121.478 | 7.549 | 4.375 | | | 178 | G |
| 151 | D | 54.546 | 41.258 | 175.682 | 121.554 | 8.02 | 4.494 | 2.688 | 2.542 | 491 | B |
| 152 | D | 54.503 | 41.069 | 174.818 | 120.288 | 8.127 | 4.551 | 2.681 | 2.624 | 209 | G |
| 153 | L | 55.77 | 42.227 | 174.26 | 122.011 | 8.08 | 4.324 | 1.687 | | 353 | G |
| 154 | I | 61.216 | 38.625 | 175.042 | 122.028 | 8.051 | 4.214 | 1.911 | | 231 | G |
| 155 | T | 61.484 | 69.913 | 176.926 | 118.792 | 8.275 | 4.477 | 4.25 | | 200 | G |
| 156 | T | 61.551 | 69.947 | 177.326 | 115.999 | 8.216 | 4.415 | 4.27 | | 244 | G |
| 157 | D | 54.554 | 41.109 | 175.998 | 122.631 | 8.349 | 4.637 | 2.648 | 2.625 | 284 | G |
| 158 | E | 57.457 | 29.671 | 175.36 | 121.391 | 8.013 | 4.259 | 1.936 | | 424 | B |

| | | | | | | | | | | | |
|---|---|---|---|---|---|---|---|---|---|---|---|
| 159 | D | 53.552 | 38.858 | | 118.258 | 8.209 | 4.645 | 2.716 | | 448 | B |
| 160 | P | 63.833 | 32.244 | 173.833 | | | 4.417 | 2.315 | 1.941 | | |
| 161 | L | 55.833 | 41.615 | 173.283 | 120.101 | 8.356 | 4.286 | 1.595 | | 279 | G |
| 162 | M | 55.911 | 32.543 | 174.763 | 119.616 | 8.025 | 4.446 | 2.101 | | 222 | G |
| 163 | S | 59.09 | 63.719 | 176.394 | 115.705 | 8.141 | 4.401 | 3.89 | | 241 | G |
| 164 | S | 58.965 | 63.806 | 176.648 | 117.789 | 8.264 | 4.476 | 3.905 | | 201 | B |
| 165 | I | 61.839 | 38.471 | 174.897 | 121.947 | 8.001 | 4.208 | 1.588 | | 248 | G |
| 166 | L | 55.644 | 42.276 | 173.43 | 124.265 | 8.161 | 4.303 | 1.674 | 1.588 | 255 | B |
| 167 | G | 45.753 | | 177.313 | 108.695 | 8.194 | 3.918 | | | 193 | G |
| 168 | D | 54.721 | 41.08 | 175.431 | 120.527 | 8.159 | 4.572 | 2.658 | 2.63 | 307 | G |
| 169 | L | 55.522 | 42.275 | 174.214 | 121.471 | 8.059 | 4.212 | 1.687 | 1.585 | 349 | B |
| 170 | L | 55.336 | 42.409 | 174.214 | 121.959 | 7.952 | 4.327 | 1.619 | 1.585 | 269 | B |
| 171 | L | 55.336 | 42.409 | 174.358 | 121.959 | 7.952 | 4.327 | 1.619 | | 269 | B |
| 172 | D | 54.375 | 41.232 | 174.641 | 121.048 | 8.239 | 4.566 | 2.698 | 2.663 | 395 | G |
| 173 | T | 62.598 | 69.51 | 176.698 | 114.364 | 8.086 | 4.214 | | | 287 | G |
| 174 | N | 53.651 | 38.814 | 176.203 | 120.259 | 8.409 | 4.712 | 2.747 | | 259 | G |
| 175 | F | 58.643 | 39.139 | 175.807 | 120.583 | 8.09 | 4.517 | 3.095 | | 508 | B |
| 176 | N | 53.264 | 38.958 | 176.083 | 120.151 | 8.295 | 4.569 | 2.816 | 2.705 | 86 | B |
| 177 | S | 59.044 | 63.733 | 176.73 | 116.497 | 8.183 | 4.33 | 3.881 | | 190 | G |
| 178 | A | 53.086 | 18.975 | 173.45 | 125.403 | 8.273 | 4.329 | 1.406 | | 250 | B |
| 179 | S | 58.715 | 63.756 | 176.797 | 114.087 | 8.079 | 4.363 | 3.854 | | 215 | G |
| 180 | K | 56.374 | 32.95 | 174.976 | 123.004 | 8.173 | 4.351 | 1.857 | 1.753 | 138 | B |
| 181 | V | 62.504 | 32.751 | 175.426 | 120.729 | 8.012 | 4.091 | 2.047 | | 296 | B |
| 182 | Q | 55.643 | 29.458 | 175.892 | 124.48 | 8.435 | 4.328 | 2.046 | 1.951 | 312 | B |
| 183 | Q | 53.747 | 29.083 | | 122.992 | 8.438 | 4.302 | 2.085 | 1.938 | 265 | B |
| 184 | P | | | | | | | | | | |
| 192 | Q | | | | | | | | | | |
| 193 | P | 63.26 | 32.136 | 174.563 | | | 4.404 | 2.3 | 1.9 | | |
| 194 | Q | 55.941 | 29.563 | 175.67 | 120.526 | 8.489 | 4.263 | 2.104 | 1.971 | 304 | G |
| 195 | A | 52.776 | 19.218 | 173.861 | 125.483 | 8.324 | 4.309 | 1.371 | | 191 | G |
| 196 | V | 62.529 | 32.676 | 175.371 | 119.612 | 8.095 | 4.084 | 2.051 | | 257 | G |
| 197 | L | 55.171 | 42.246 | 173.988 | 126.056 | 8.273 | 4.32 | | | 254 | G |
| 198 | Q | 56.017 | 30.861 | | 121.34 | 8.201 | | | | 141 | G |
| 199 | Q | | | | | | | | | | |
| 200 | P | 63.248 | 32.19 | 174.42 | | | 4.464 | 2.315 | 1.933 | | |
| 201 | S | 58.66 | 63.97 | 176.723 | 116.068 | 8.478 | 4.455 | 3.884 | | 183 | G |
| 202 | S | 58.59 | 63.897 | 177.072 | 117.452 | 8.348 | 4.498 | | | 163 | G |
| 203 | C | 58.732 | 27.951 | 176.934 | 121.048 | 8.37 | 4.605 | | | 376 | B |
| 204 | V | 62.683 | 32.712 | 175.556 | 122.364 | 8.156 | 4.121 | | | 286 | G |
| 205 | E | 56.372 | 30.149 | 175.493 | 124.652 | 8.423 | 4.329 | 2.015 | | 493 | B |

| | | | | | | | | | | | |
|---|---|---|---|---|---|---|---|---|---|---|---|
| 206 | L | 55.108 | 42.376 | 174.509 | 124.399 | 8.293 | 4.371 | | | 295 | B |
| 207 | R | 53.982 | 30.257 | | 123.193 | 8.367 | | | | 215 | B |
| 208 | P | 63.46 | 31.949 | 174.596 | | | 4.289 | 1.634 | 1.575 | | |
| 209 | L | 55.317 | 42.164 | | 121.112 | 8.093 | 4.312 | 1.629 | | 181 | B |
| 210 | D | 54.322 | 41.108 | 175.117 | 121.138 | 8.262 | 4.566 | 2.698 | 2.639 | 325 | G |
| 211 | R | 56.101 | 30.59 | 174.946 | 121.71 | 8.308 | 4.408 | 1.96 | | 128 | G |
| 212 | T | 62.595 | 69.789 | 176.699 | 115.429 | 8.314 | 4.307 | 4.204 | | 187 | G |
| 213 | V | 62.538 | 32.807 | 175.256 | 122.24 | 8.095 | 4.173 | 2.107 | | 288 | G |
| 214 | S | 58.441 | 64.025 | 176.743 | 119.48 | 8.406 | 4.509 | 3.878 | | 222 | B |
| 215 | S | 58.605 | 63.873 | | 117.992 | 8.391 | 4.479 | 3.888 | | 170 | G |
| 216 | N | 53.4 | 38.9 | | | | | | | | |
| 217 | S | 58.73 | 63.86 | | | | | | | | |
| 218 | N | 53.4 | 38.9 | | | | | | | | |
| 219 | N | 53.36 | 38.93 | | | | | | | | |
| 220 | N | 53.36 | 38.93 | | | | | | | | |
| 221 | S | 58.73 | 63.86 | | | | | | | | |
| 222 | N | 53.4 | 38.9 | | | | | | | | |
| 223 | S | 58.73 | 63.86 | | | | | | | | |
| 224 | N | 53.4 | 38.9 | | | | | | | | |
| 225 | N | 53.36 | 38.93 | 176.833 | | | 4.687 | 2.834 | 2.738 | | |
| 226 | A | 52.499 | 19.235 | 175.171 | 124.742 | 8.168 | 4.316 | 1.385 | | 481 | B |
| 227 | A | 53.91 | 20.079 | | 129.201 | 7.879 | 4.114 | 1.34 | | 268 | G |

**Supplementary Table 4. Unassigned PHL4$_{effector}$ signals.** Signals labeled n# and s# in the first column and in Figure 4 were ambigously assigned to the C-terminal NS-rich region. The "Comments" column indicates likely assignments. Note that some peaks likely represent alternative conformations, such as cis-proline pairs, as determined by typical chemical shift values for cis- and trans-proline residues. SN is the signal-to-noise output from analysis of the non-NUS $^{1}H$–$^{15}N$ HSCQ in NMRFAM-Sparky (Lee et al., 2015).

| Name in Figure | N | CA | CB | $H_N$ | H | HB | HB2 | CA (*i* – 1) | CB (*i* – 1) | CO (*i* – 1) | HA (*i* – 1) | HB (*i* – 1) | HB2 (*i* – 1) | SN | Comments |
|---|---|---|---|---|---|---|---|---|---|---|---|---|---|---|---|
| n1 | 120.35 | 53.41 | 38.84 | 8.45 | 4.75 | 2.82 | – | 58.70 | 63.74 | 177.08 | 4.43 | 3.88 | – | 360 | Used at Cterm |
| n2 | 120.57 | 53.45 | 38.93 | 8.46 | 4.75 | 2.82 | – | 58.72 | 63.77 | 177.02 | 4.43 | 3.88 | – | 251 | Used at Cterm |
| n3 | 120.69 | 53.48 | 38.93 | 8.45 | 4.75 | 2.82 | – | 58.70 | 63.84 | 177.07 | 4.43 | 3.88 | – | 219 | Used at Cterm |
| n4 | 118.96 | 53.48 | 38.76 | 8.35 | 4.71 | 2.82 | – | 53.35 | 38.77 | 176.48 | 4.73 | 2.86 | 2.77 | 189 | Likely at Cterm LC region |
| n5 | 119.17 | 53.31 | 38.97 | 8.32 | 4.70 | 2.79 | – | 53.34 | 38.75 | 176.53 | 4.72 | 2.86 | 2.76 | 244 | Likely at Cterm LC region |
| n6 | 119.15 | 53.30 | 39.07 | 8.41 | 4.73 | 2.81 | – | 53.40 | 38.72 | 176.41 | 4.71 | 2.86 | 2.76 | 374 | Likely at Cterm LC region |
| s1 | 116.12 | 58.73 | 63.86 | 8.29 | 4.43 | 3.88 | – | 53.46 | 38.95 | 176.08 | 4.78 | 2.87 | 2.79 | 428 | Likely Cterm LC region |
| s2 | 115.93 | 58.73 | 63.84 | 8.25 | 4.43 | 3.88 | – | 53.36 | 38.91 | 176.06 | 4.77 | 2.88 | 2.79 | 211 | Likely Cterm LC region |
| s3 | 115.98 | – | – | 8.31 | – | – | – | 53.55 | 38.86 | 176.01 | 4.75 | 2.87 | 2.79 | 194 | unknown before a N, |

| | | | | | | | | | | | | | | | |
|---|---|---|---|---|---|---|---|---|---|---|---|---|---|---|---|
| | | | | | | | | | | | | | | | perhaps ser at C-term? |
| | 120.45 | 52.93 | – | 7.80 | – | – | – | 55.20 | 42.76 | 175.78 | – | – | – | 18 | weak, potentially alt conformer for DA |
| | 120.41 | 56.36 | 39.93 | 7.86 | 4.43 | 2.92 | – | 53.05 | 39.09 | 178.04 | 4.71 | 2.80 | 2.68 | 144 | Likely Cterm LC region |
| | 120.51 | 56.38 | 40.06 | 7.88 | – | – | – | 53.12 | 39.21 | 178.01 | – | – | – | 74 | Likely Cterm LC region |
| | 123.36 | 54.87 | 41.13 | 7.97 | – | – | – | 52.45 | 19.45 | 174.80 | 4.29 | 1.37 | – | 29 | weak AD alternative conformer at D70 |
| | 122.96 | 52.84 | – | 7.98 | – | – | – | 58.14 | 63.98 | 178.49 | – | – | – | 49 | weak alt conformer before Ser |
| | 121.47 | 56.42 | 38.37 | 8.02 | 4.80 | 3.06 | – | 58.41 | – | – | – | – | – | 445 | overlapped! |
| | 115.86 | 59.31 | 63.73 | 8.03 | 4.37 | 3.93 | – | 53.68 | 38.83 | 175.62 | 4.70 | 2.88 | 2.78 | 489 | likely N term Asn in LC region |
| | 115.86 | 59.31 | 63.73 | 8.03 | 4.37 | 3.93 | – | 53.68 | 38.83 | 175.62 | – | – | – | 489 | likely N term Asn |
| | 122.56 | 55.31 | 42.48 | 8.04 | – | – | – | 53.32 | 38.62 | 176.71 | – | – | – | 198 | unknown |
| | 121.42 | 56.35 | 32.77 | 8.07 | – | – | – | 55.94 | 32.25 | 175.10 | – | – | – | 159 | unknown |
| | 115.79 | 58.95 | 63.79 | 8.08 | 4.41 | 3.93 | – | 53.51 | 38.92 | 175.83 | 4.71 | 2.87 | 2.77 | 267 | likely N term Asn in LC region |
| | 121.47 | 55.29 | 42.07 | 8.12 | 4.29 | – | – | 55.99 | 29.59 | – | – | – | – | 499 | unknown, Bad CO |
| | 121.73 | 56.22 | 32.95 | 8.13 | 4.29 | 3.08 | – | 55.71 | 32.34 | 175.17 | 4.48 | – | – | 257 | Likely N term LC domain |
| | 118.67 | 53.07 | 39.13 | 8.18 | 4.67 | 2.68 | – | 56.42 | 32.79 | 175.49 | 4.23 | 1.70 | 1.39 | 376 | Likely N term LC region |
| | 124.07 | 53.44 | 18.91 | 8.20 | 4.21 | 1.40 | – | 54.86 | 41.09 | 174.74 | 4.56 | 2.70 | – | 214 | Likely N term 'NA' pair |
| | 120.78 | 54.82 | 41.16 | 8.21 | 4.58 | 2.68 | – | 54.65 | 41.10 | 175.20 | 4.59 | 2.71 | 2.64 | 364 | Likely Cterm LC region in Asp repeat |
| | 124.13 | 58.73 | 38.85 | 8.23 | – | – | – | 57.96 | 63.86 | 177.54 | 4.46 | 3.81 | – | 107 | SIP' alt conformer, or truncation artifact? |
| | 121.40 | 55.84 | 32.45 | 8.24 | 4.38 | 2.06 | – | 59.15 | 63.67 | 176.55 | 4.37 | 3.93 | – | 297 | Likely N term LC region SM |
| | 120.71 | 54.88 | 41.11 | 8.25 | 4.57 | 2.70 | – | 54.97 | 41.08 | 174.94 | 4.56 | 2.70 | 2.63 | 582 | Likely Cterm LC region in Asp repeat |
| | 121.42 | 55.79 | 32.40 | 8.26 | 4.39 | 2.05 | – | 59.17 | 63.65 | 176.52 | 4.38 | 3.93 | – | 303 | Likely N term LC region SM |
| | 124.06 | 53.82 | 18.77 | 8.26 | 4.20 | 1.42 | – | 54.86 | 41.15 | 174.48 | 4.55 | 2.70 | – | 223 | Likely N term 'NA' pair |
| | 120.85 | 54.82 | 41.13 | 8.27 | 4.57 | 2.70 | – | 55.48 | 41.12 | – | 4.56 | 2.70 | 2.65 | 630 | Bad CO, overlapped CA / CB |
| | 120.18 | 54.71 | 41.18 | 8.28 | 4.57 | 2.70 | – | 54.71 | 41.18 | 175.36 | 4.58 | 2.72 | 2.63 | 451 | Likely Cterm LC region in Asp repeat |
| | 121.12 | 54.92 | 41.11 | 8.29 | 4.57 | 2.70 | – | 54.65 | 41.13 | 174.91 | 4.57 | 2.71 | 2.65 | 389 | Likely Cterm LC region in Asp repeat |
| | 122.37 | 55.09 | 42.50 | 8.29 | 4.38 | 1.62 | – | 63.09 | 32.03 | 174.80 | 4.43 | 2.25 | 1.92 | 676 | Multiple 'PL' choices |
| | 116.53 | 53.58 | 38.85 | 8.29 | – | – | – | 53.59 | 18.96 | – | 4.32 | 1.40 | – | 312 | Bad CO, likely N term LC region |
| | 116.12 | 53.99 | 38.88 | 8.30 | 4.71 | 2.83 | 2.83 | 53.60 | 18.88 | 172.96 | 4.21 | 1.41 | – | 718 | Likely N term LC region |
| | 124.15 | 53.81 | 18.75 | 8.30 | 4.21 | 1.42 | – | 54.78 | 41.12 | 174.50 | 4.55 | 2.71 | – | 231 | Likely N term 'NA' pair |
| | 117.29 | 59.23 | 63.78 | 8.30 | 4.60 | – | – | 53.13 | 38.93 | 176.19 | 4.68 | 2.82 | 2.69 | 242 | Could be Cterm, SN |
| | 120.54 | 54.88 | 41.06 | 8.30 | 4.57 | 2.68 | – | 54.66 | 41.09 | 175.27 | 4.60 | 2.70 | 2.63 | 631 | Likely Cterm LC region in Asp repeat |

| | | | | | | | | | | | | | | | |
|---|---|---|---|---|---|---|---|---|---|---|---|---|---|---|---|
| | 118.90 | 52.91 | 39.00 | 8.33 | 4.70 | 2.79 | – | 56.53 | 32.96 | 175.33 | 4.26 | 1.75 | – | 143 | Likely N term LC region |
| | 121.67 | 58.88 | 32.63 | 8.34 | – | – | – | 54.95 | 42.35 | 174.40 | 4.39 | 1.60 | – | 508 | unknown |
| | 121.07 | 56.29 | 29.42 | 8.36 | – | – | – | 55.77 | 39.26 | 175.04 | 4.67 | 2.79 | – | 527 | unknown |
| | 122.00 | 55.38 | 42.35 | 8.36 | 4.39 | – | – | 63.32 | 32.11 | 174.38 | – | – | – | 482 | Questionable CB; multiple 'PL' choices |
| | 123.06 | 53.88 | 30.26 | 8.36 | – | – | – | 62.71 | 34.07 | 175.66 | – | – | – | 114 | unknown, not too intense |
| | 109.15 | 45.70 | – | 8.40 | – | – | – | 53.17 | 38.83 | 175.60 | – | – | – | 19 | weak NG at G92 |
| | 121.96 | 55.85 | 29.26 | 8.40 | 4.37 | – | – | 55.68 | 41.95 | 173.97 | – | – | – | 67 | unknown |
| | 123.17 | 52.19 | 41.47 | 8.40 | – | – | – | 56.24 | 30.58 | 175.62 | – | – | – | 284 | unknown |
| | 114.23 | 45.85 | – | 8.42 | – | – | – | 54.83 | 41.08 | 174.90 | – | – | – | 34 | weak, alt G135 signal? |
| | 115.74 | 58.44 | 63.95 | 8.43 | 4.43 | 3.89 | – | 63.32 | 32.20 | 174.50 | 4.44 | 2.31 | 1.93 | 193 | 125 or 185 or 201 (PS) |
| | 110.61 | 45.29 | – | 8.44 | – | – | – | 58.90 | 63.93 | 176.43 | 4.38 | 3.91 | – | 14 | weak, alt G126 conformer |
| | 123.20 | 53.85 | 28.92 | 8.45 | 4.61 | – | – | 55.65 | 29.61 | 175.89 | 4.34 | 2.06 | 1.96 | 457 | One of the two QQP pairs |
| | 122.77 | 55.54 | 42.20 | 8.49 | – | – | – | 62.65 | 34.16 | 175.67 | 3.93 | 1.94 | – | 156 | Multiple 'PL' choices |
| | 117.76 | 53.23 | 38.84 | 8.50 | – | – | – | 52.93 | 19.11 | 173.75 | – | – | – | 17 | weak NA alternative conformer |
| | 117.10 | 58.99 | 63.85 | 8.50 | – | – | – | 63.07 | 34.26 | 175.14 | – | – | – | 34 | weak Cis Proline conformer |
| | 116.30 | 58.43 | 63.85 | 8.51 | – | – | – | 62.84 | 34.34 | 175.36 | – | – | – | 25 | weak Cis Proline conformer PS |
| | 116.49 | 58.61 | 63.96 | 8.54 | – | – | – | 62.85 | 34.29 | 175.16 | – | – | – | 21 | weak Cis Proline conformer PS |
| | 115.19 | 58.71 | 64.12 | 8.56 | 4.50 | – | – | 63.18 | 33.33 | 175.29 | – | – | – | 18 | weak Cis Proline conformer PS |
| | 119.44 | 53.48 | 39.40 | 8.64 | 4.71 | 2.84 | – | 63.21 | 34.34 | 175.81 | 4.75 | – | – | 46 | Likely minor proline Cis species, perhaps for a PN pair |
| | 122.83 | – | – | 8.66 | – | – | – | – | – | – | – | – | – | 12 | unknown |
| | 124.83 | 52.85 | 19.17 | 8.70 | – | – | – | 62.54 | – | 175.39 | – | – | – | 18 | weak PA alternative conformer, at A90 |
| | 121.70 | 57.29 | – | 8.72 | – | – | – | 63.09 | – | 175.53 | – | – | – | 19 | weak Cis Proline conformer |
| | 120.60 | 51.61 | 39.16 | 8.83 | – | – | – | 62.48 | 34.48 | 175.66 | – | – | – | 23 | weak Cis Proline conformer |

**Supplementary Table 5. Assigned NMR chemical shifts for PHL4$_{FL}$.** Column 1 is the residue number, SN represents the peak intensity calculated by NMRFAM-SPARKY for the $^{1}$H-$^{15}$N HSQC for PHL4$_{FL}$. For the SN Type, "G" stands for non-overlapped peaks in the HSQC spectrum, whereas "B" represents overlapping peaks in the HSQC spectrum. The assigned NMR chemical shift values come from the NCACB, HNCA, HN(CO)CACB, HNCO spectra. In cases where overlapping peaks were observed, the values are taken from the spectrum without any peak overlap. Peaks without N or H$_N$ shifts, but having other nuclei with assigned chemical shifts, come from experiments reporting on the chemical shifts for "$i-1$" residues. Black bars represent long amino acid sequences without assignments.

| | AA Type | CA | CB | CO | N | $H_N$ | SN | SN Type* |
|---|---|---|---|---|---|---|---|---|
| 0 | G | | | | | | | |
| 1 | M | 55.835 | 33.14 | 175.465 | | | | |
| 2 | I | 58.441 | 38.774 | | 123.537 | 8.307 | 182 | B |
| 3 | P | 63.282 | 32.199 | 174.929 | | | | |
| 4 | N | 53.241 | 39.125 | 176.401 | 119.125 | 8.541 | 170 | G |
| 5 | D | 54.649 | 41.165 | | 121.04 | 8.349 | 374 | B |
| 6 | D | | | | | | | |
| [redacted] | | | | | | | | |
| 16 | P | | | | | | | |
| 17 | L | 55.174 | 42.405 | 174.298 | | | | |
| 18 | N | 53.066 | 39.272 | 176.524 | 119.276 | 8.424 | 297 | B |
| 19 | D | 54.567 | 41.139 | | 121.195 | 8.386 | 352 | G |
| 20 | D | | | | | | | |
| [redacted] | | | | | | | | |
| 29 | P | | | | | | | |
| 30 | L | 55.174 | 42.405 | 174.298 | | | | |
| 31 | N | 53.066 | 39.272 | 176.524 | 119.276 | 8.424 | 297 | B |
| 32 | D | 54.567 | 41.139 | | 121.195 | 8.386 | 352 | G |
| 33 | D | | | | | | | |
| [redacted] | | | | | | | | |
| 38 | M | | | | | | | |
| 39 | E | 56.888 | 30.161 | 175.324 | | | | |
| 40 | N | 52.889 | 39.109 | 177.174 | 118.523 | 8.316 | 49 | B |
| 41 | Y | 56.063 | 38.346 | | 121.724 | 8.025 | 260 | B |
| 42 | P | | | | | | | |
| 43 | L | 55.316 | 42.03 | 173.968 | | | | |
| 44 | R | 56.085 | 30.961 | 175.352 | 121.34 | 8.201 | 82 | B |
| 45 | S | 57.945 | 63.71 | 177.524 | 117.01 | 8.253 | 64 | G |
| 46 | I | 58.688 | 38.811 | | 124.233 | 8.174 | 85 | B |
| 47 | P | 63.321 | 32.232 | 174.25 | | | | |
| 48 | T | 62.411 | 69.691 | 176.521 | 114.886 | 8.209 | 92 | G |
| 49 | E | 56.7 | 30.156 | 174.944 | 123.107 | 8.524 | 95 | G |
| 50 | L | 55.304 | 42.268 | 173.944 | 122.998 | 8.228 | 98 | G |
| 51 | S | 58.683 | 63.785 | 176.897 | 115.803 | 8.186 | 102 | B |
| 52 | H | 55.732 | 29.53 | 176.549 | 121.999 | 8.363 | 57 | B |
| 53 | T | 62.046 | 69.803 | 174.701 | 115.206 | 8.184 | 145 | B |
| 54 | C | 58.41 | 28.118 | 177.017 | 121.676 | 8.425 | 93 | B |
| 55 | S | 58.322 | 63.773 | 177.298 | 118.595 | 8.423 | 57 | G |
| 56 | L | 55.153 | 42.249 | 174.598 | 124.476 | 8.263 | 63 | G |
| 57 | I | 58.357 | 38.661 | | 123.975 | 8.137 | 77 | B |

| | | | | | | | | |
|---|---|---|---|---|---|---|---|---|
| 58 | P | | | | | | | |
| 59 | P | 62.832 | 32.117 | 174.631 | | | | |
| 60 | S | 57.912 | 63.819 | 177.39 | 116.059 | 8.308 | 274 | B |
| 61 | L | 52.864 | 41.935 | | 125.427 | 8.285 | 75 | B |
| 62 | P | 63.153 | 32.106 | 175.072 | | | | |
| 63 | N | 51.188 | 38.71 | | 120.024 | 8.533 | 76 | G |
| 64 | P | 63.762 | 32.18 | 174.129 | | | | |
| 65 | S | 58.934 | 63.553 | 176.626 | 115.099 | 8.307 | 95 | G |
| 66 | E | 56.599 | 30.326 | 175.164 | 122.507 | 8.194 | 103 | G |
| 67 | A | 52.402 | 19.231 | 173.72 | 124.742 | 8.168 | 90 | G |
| 68 | A | 52.531 | 19.222 | 173.879 | 123.163 | 8.202 | 105 | B |
| 69 | A | 52.625 | 19.314 | 173.879 | 123.25 | 8.174 | 94 | B |
| 70 | D | 54.467 | 41.139 | 174.87 | 119.083 | 8.223 | 93 | G |
| 71 | M | 55.691 | 32.574 | | 121.381 | 8.29 | 64 | B |
| 72 | S | 58.893 | 63.639 | 177.033 | | | | |
| 73 | F | 57.964 | 39.482 | 176.065 | 121.818 | 8.14 | 81 | B |
| 74 | N | 53.073 | 39.148 | | 120.863 | 8.232 | 143 | B |
| 75 | S | 59.092 | 63.619 | 176.396 | | | | |
| 76 | E | 57.31 | 29.768 | 174.408 | 122.298 | 8.439 | 96 | G |
| 77 | L | 55.701 | 42.201 | 174.062 | 121.062 | 7.977 | 58 | G |
| 78 | N | 53.646 | 38.675 | 176.184 | 118.277 | 8.211 | 114 | B |
| 79 | Q | 55.898 | 29.53 | 175.448 | 119.761 | 8.127 | 46 | G |
| 80 | I | 60.974 | 38.575 | 175.403 | 121.617 | 8.09 | 55 | B |
| 81 | M | 55.156 | 32.902 | 175.842 | 124.919 | 8.413 | 20 | B |
| 82 | A | 52.38 | 19.361 | 174.231 | 125.269 | 8.183 | 34 | G |
| 83 | R | 53.966 | 30.275 | | 121.552 | 8.352 | 90 | B |
| 84 | P | 63.746 | 31.992 | 174.157 | | | | |
| 85 | C | 58.473 | 27.962 | 177.049 | 117.834 | 8.374 | 48 | B |
| 86 | D | 54.584 | 41.062 | 175.512 | 122.545 | 8.291 | 160 | B |
| 87 | M | 55.201 | 32.931 | 175.699 | 119.766 | 8.049 | 43 | G |
| 88 | L | 53.307 | 41.742 | | 124.61 | 8.197 | 63 | G |
| 89 | P | 62.785 | 32.126 | 174.498 | | | | |
| 90 | A | 52.7 | 19.042 | 173.581 | 124.364 | 8.439 | 45 | B |
| 91 | N | 53.123 | 38.866 | 175.576 | 117.305 | 8.411 | 41 | G |
| 92 | G | 45.707 | | 176.727 | 109.025 | 8.355 | 44 | B |
| 93 | G | 45.115 | | 177.679 | 108.378 | 8.221 | 43 | G |
| 94 | A | 52.32 | 19.546 | 173.567 | 123.724 | 8.136 | 73 | B |
| 95 | V | 62.355 | 32.566 | 174.679 | 119.02 | 8.068 | 20 | B |
| 96 | G | 45.094 | | 177.774 | 112.139 | 8.426 | 33 | G |
| 97 | H | 55.322 | 29.909 | 177.437 | 118.33 | 8.243 | 47 | B |
| 98 | N | | 39.138 | | 121.924 | 8.507 | 22 | B |

| | | | | | | | | |
|---|---|---|---|---|---|---|---|---|
| 99 | P | 63.645 | 32.106 | 174.866 | | | | |
| 100 | F | 57.699 | 38.795 | 175.889 | 118.283 | 8.002 | 20 | G |
| 101 | L | 54.663 | 42.6 | 174.887 | 121.858 | 7.68 | 15 | G |
| 102 | E | 54.442 | 29.827 | | 122.834 | 8.095 | 28 | B |
| 103 | P | | | | | | | |
| 104 | G | 45.163 | | 177.417 | | | | |
| 105 | F | 57.789 | 39.838 | | 120.045 | 7.987 | 32 | G |
| 106 | N | 53.027 | 39.125 | 177.336 | | | | |
| 107 | C | | 27.517 | | 120.843 | 8.182 | 40 | B |
| 108 | P | 63.465 | 32.106 | 174.585 | | | | |
| 109 | E | 56.88 | 30.032 | 174.659 | 120.54 | 8.532 | 41 | G |
| 110 | T | 61.627 | 70.23 | | 114.565 | 8.126 | 19 | G |
| 111 | T | 61.728 | 69.735 | 177.177 | | | | |
| 112 | D | 55.089 | 40.987 | | 122.486 | 8.291 | 212 | B |
| 113 | W | | | | | | | |
| [redacted] | | | | | | | | |
| 123 | F | | | | | | | |
| 124 | P | 63.219 | 31.961 | 174.486 | | | | |
| 125 | S | 58.687 | 63.995 | | 115.842 | 8.357 | 34 | G |
| 126 | G | 45.15 | | 177.631 | | | | |
| 127 | S | | | | 116.62 | 8.148 | 15 | B |
| 128 | P | | | | | | | |
| 129 | N | | | | | | | |
| 130 | L | 55.788 | 42.254 | 175.398 | | | | |
| 131 | I | 61.563 | 38.595 | 175.102 | 121.474 | 8.115 | 88 | B |
| 132 | M | 55.98 | 32.592 | 175.312 | 123.762 | 8.306 | 43 | B |
| 133 | E | 56.607 | 30.389 | 175.323 | 120.66 | 8.26 | 335 | B |
| 134 | D | 54.664 | 41.15 | 174.677 | 121.386 | 8.417 | 60 | B |
| 135 | G | 45.545 | | 177.439 | 109.03 | 8.33 | 32 | B |
| 136 | V | 62.654 | 32.797 | | 120.104 | 7.922 | 23 | G |
| 137 | I | 61.007 | 38.703 | 175.623 | | | | |
| 138 | D | 54.213 | 41.255 | 175.382 | 124.674 | 8.357 | 18 | G |
| 139 | E | 56.986 | 30.187 | | 121.484 | 8.338 | 77 | B |
| 140 | I | | | | | | | |
| 141 | H | | | | | | | |
| 142 | K | | | | | | | |
| 143 | Q | 56.758 | 30.159 | 174.751 | | | | |
| 144 | S | 58.465 | 63.893 | 177.35 | 116.575 | 8.289 | 334 | B |
| 145 | D | 54.368 | 41.194 | 175.712 | 122.348 | 8.346 | 52 | B |
| 146 | L | | | | 122.983 | 7.995 | 12 | B |
| 147 | P | | | | | | | |

| | | | | | | | | |
|---|---|---|---|---|---|---|---|---|
| 157 | D | | | | | | | |
| 158 | E | 56.872 | 29.933 | 175.34 | | | | |
| 159 | D | 53.308 | 38.969 | | 118.258 | 8.209 | 124 | B |
| 160 | P | 63.576 | | 173.813 | | | | |
| 161 | L | | | | 120.101 | 8.356 | 27 | G |
| 162 | M | | | | | | | |
| 163 | S | | | | | | | |
| 164 | S | 58.919 | 63.473 | 176.628 | | | | |
| 165 | I | 61.839 | 38.471 | | 121.947 | 8.001 | 18 | B |
| 166 | L | | | | | | | |
| 167 | G | 45.714 | | 177.293 | | | | |
| 168 | D | 54.751 | 41.217 | | 120.527 | 8.159 | 20 | B |
| 169 | L | 55.307 | 41.838 | 174.194 | | | | |
| 170 | L | 55.307 | 41.838 | 174.194 | 121.959 | 7.952 | 14 | B |
| 171 | L | | | | 121.959 | 7.952 | 14 | B |
| 172 | D | 54.389 | 41.23 | 174.62 | | | | |
| 173 | T | | | | 114.378 | 8.089 | 17 | B |
| 174 | N | 53.534 | 38.784 | 176.148 | | | | |
| 175 | F | 58.643 | 38.612 | | 120.583 | 8.09 | 37 | G |
| 176 | N | 53.549 | 38.863 | 176.027 | | | | |
| 177 | S | 57.812 | 63.724 | 176.71 | 116.41 | 8.188 | 23 | G |
| 178 | A | 52.84 | 18.928 | 173.409 | 125.398 | 8.28 | 80 | B |
| 179 | S | 59.052 | 63.756 | | 114.188 | 8.083 | 24 | B |
| 180 | K | | | | | | | |
| 181 | V | 62.471 | 32.557 | 175.371 | | | | |
| 182 | Q | 55.647 | 29.668 | 175.872 | 124.48 | 8.435 | 39 | B |
| 183 | Q | | | | 122.992 | 8.438 | 30 | B |
| 184 | P | | | | | | | |
| 192 | Q | | | | | | | |
| 193 | P | 63.613 | 32.224 | 174.543 | | | | |
| 194 | Q | | | | 120.526 | 8.489 | 40 | B |
| 195 | A | | | | | | | |
| 196 | V | | | | | | | |
| 197 | L | 55.316 | 42.03 | 173.968 | | | | |
| 198 | Q | 56.017 | 31.016 | | 121.34 | 8.201 | 82 | B |
| 199 | Q | | | | | | | |
| 200 | P | 63.119 | 32.19 | 174.377 | | | | |
| 201 | S | | | | 116.058 | 8.485 | 12 | G |
| 202 | S | | | | | | | |

| | | | | | | | | |
|---|---|---|---|---|---|---|---|---|
| 207 | R | | | | | | | |
| 208 | P | 63.46 | 31.949 | 174.576 | | | | |
| 209 | L | 55.432 | 42.117 | | 121.112 | 8.093 | 78 | G |
| 210 | D | | | | | | | |
| 213 | V | | | | | | | |
| 214 | S | 58.507 | 63.918 | 176.723 | | | | |
| 215 | S | 58.421 | | | 117.992 | 8.391 | 30 | B |
| 216 | N | | | | | | | |
| 368 | E | | | | | | | |
| 369 | Q | 56.009 | 29.487 | 174.934 | | | | |
| 370 | G | 45.232 | | 177.392 | 109.656 | 8.364 | 89 | G |
| 371 | E | 56.569 | 30.357 | | 120.763 | 8.327 | 197 | B |
| 372 | K | 56.12 | 32.864 | 174.474 | | | | |
| 373 | T | 61.77 | 69.781 | 176.84 | 115.193 | 8.194 | 114 | B |
| 374 | S | 58.189 | 63.909 | 177.291 | 118.165 | 8.334 | 124 | G |
| 375 | A | 52.463 | 19.381 | 173.945 | 126.115 | 8.319 | 123 | G |
| 376 | K | 55.978 | 33.14 | 174.928 | 121.095 | 8.294 | 498 | G |
| 377 | T | 59.981 | 69.626 | | 118.516 | 8.278 | 126 | G |
| 378 | P | 62.877 | 32.279 | 174.684 | | | | |
| 379 | E | 56.356 | 30.832 | 175.274 | 121.039 | 8.456 | 156 | G |
| 380 | N | 53.334 | 38.984 | 175.613 | 119.294 | 8.498 | 188 | B |
| 381 | G | 45.571 | | 177.105 | 109.679 | 8.457 | 145 | G |
| 382 | S | 58.433 | 63.975 | 176.678 | 115.572 | 8.232 | 195 | G |
| 383 | E | 56.912 | 30.058 | 174.666 | 122.781 | 8.611 | 205 | G |
| 384 | E | 56.912 | 30.2 | | 121.64 | 8.451 | 243 | G |
| 385 | S | 58.61 | 63.847 | 176.914 | | | | |
| 386 | E | 56.381 | 30.393 | 175.134 | 122.289 | 8.367 | 193 | B |
| 387 | S | 56.566 | 63.379 | | 118.147 | 8.288 | 154 | G |
| 388 | P | 63.003 | 32.06 | 174.668 | | | | |
| 389 | R | 53.987 | 30.37 | | 122.536 | 8.378 | 147 | B |
| 390 | P | 62.898 | 32.192 | 174.641 | | | | |
| 391 | K | 56.123 | 33.073 | 174.986 | 122.163 | 8.471 | 141 | G |
| 392 | R | 53.904 | 30.448 | | 123.374 | 8.356 | 120 | G |
| 393 | P | 63.292 | 32.17 | 174.318 | | | | |
| 394 | R | 56.94 | 30.085 | 174.94 | 121.338 | 8.601 | 169 | G |
| 395 | N | 53.543 | 38.758 | 176.515 | 119.469 | 8.502 | 162 | B |
| 396 | E | 56.45 | 30.462 | 176.065 | 121.179 | 8.309 | 253 | B |
| 397 | E | 58.172 | 31.049 | | 126.716 | 7.955 | 270 | G |

**Supplementary Table 6. Unassigned PHL4FL signals.** SN is the value signal-to-noise output from analysis of the $^{1}H$–$^{15}N$ HSQC in NMRFAM-Sparky (Lee, et al., 2015).

| N | CA | CB | HN | CA (i – 1) | CB (i – 1) | CO (i – 1) | SN |
|---|---|---|---|---|---|---|---|
| 115.841 | – | – | 8.426 | 63.441 | 32.201 | 174.437 | 17 |
| 115.926 | 58.727 | 63.84 | 8.25 | 53.495 | 38.803 | 175.967 | 59 |
| 115.793 | 58.951 | 63.787 | 8.079 | 53.507 | 38.856 | 175.782 | 164 |
| 115.863 | 59.307 | 63.729 | 8.033 | 53.681 | 38.827 | 175.599 | 337 |
| 115.863 | 59.307 | 63.729 | 8.033 | 53.681 | 38.827 | 175.599 | 337 |
| 115.797 | 61.971 | 69.795 | 8.173 | – | – | 176.801 | 69 |
| 116.118 | 53.988 | 38.879 | 8.295 | 53.637 | 18.833 | 172.932 | 417 |
| 116.526 | 53.582 | 38.853 | 8.294 | 53.587 | 18.956 | 173.269 | 318 |
| 116.056 | 58.73 | 63.861 | 8.276 | 53.463 | 38.916 | 176.014 | 152 |
| 117.288 | 59.233 | 63.776 | 8.303 | 52.974 | 38.876 | 176.149 | 57 |
| 117.789 | 58.965 | 63.426 | 8.264 | – | – | 176.36 | 17 |
| 117.992 | 58.421 | – | 8.391 | 58.507 | 63.918 | 176.723 | 30 |
| 118.665 | 52.905 | 39.134 | 8.18 | 56.417 | 32.822 | 175.436 | 196 |
| 119.277 | 62.496 | 32.739 | 8.112 | 52.31 | 19.375 | 173.636 | 45 |
| 118.901 | 52.92 | 39.063 | 8.327 | 56.555 | 33.031 | 175.311 | 101 |
| 118.96 | 53.151 | 38.761 | 8.351 | 53.303 | 38.652 | 176.413 | 47 |
| 119.081 | 53.302 | 39.074 | 8.401 | 53.426 | 38.704 | 176.36 | 95 |
| 119.293 | – | – | 8.4 | – | – | 175.278 | 105 |
| 119.245 | – | – | 8.095 | – | – | 173.68 | 29 |
| 120.408 | 56.318 | 40.082 | 7.86 | 52.91 | 39.035 | 178.023 | 95 |
| 120.508 | 56.379 | 40.057 | 7.878 | 52.986 | 39.063 | 177.99 | 44 |
| 120.288 | – | – | 8.127 | – | – | 175.662 | 13 |
| 120.202 | 54.71 | 41.181 | 8.271 | 54.71 | 41.181 | 175.312 | 283 |
| 119.421 | 53.437 | 39.449 | 8.641 | 63.212 | 34.455 | 175.766 | 33 |
| 120.349 | 53.657 | 38.959 | 8.448 | 58.645 | 63.742 | 177.061 | 64 |
| 120.566 | 53.589 | 38.93 | 8.457 | 58.718 | 63.767 | 177.004 | 68 |
| 120.691 | 53.477 | 38.927 | 8.445 | 58.695 | 63.836 | 177.051 | 58 |
| 120.541 | 54.884 | 41.056 | 8.304 | 54.657 | 41.093 | 175.25 | 478 |
| 120.744 | – | – | 8.327 | – | – | 176.075 | 197 |
| 121.075 | 54.916 | 41.114 | 8.294 | 54.65 | 41.129 | 174.895 | 498 |
| 120.775 | 54.823 | 41.16 | 8.213 | 54.533 | 41.102 | 175.152 | 301 |
| 120.713 | 54.879 | 41.114 | 8.249 | 54.973 | 41.084 | 174.916 | 448 |
| 121.048 | 54.375 | 41.232 | 8.239 | – | – | 174.338 | 46 |
| 120.85 | 54.817 | 41.13 | 8.268 | 55.483 | 41.122 | 174.843 | 372 |
| 121.421 | 55.788 | 32.403 | 8.257 | 59.17 | 63.652 | 176.499 | 227 |
| 121.401 | 55.844 | 32.525 | 8.239 | 59.15 | 63.666 | 176.533 | 198 |
| 122.015 | – | – | 8.495 | – | – | 174.873 | 34 |

| | | | | | | | |
|---|---|---|---|---|---|---|---|
| 121.391 | – | – | 8.013 | 55.243 | 41.095 | 175.978 | 111 |
| 121.423 | 56.349 | 32.828 | 8.073 | 55.696 | 32.203 | 175.053 | 175 |
| 121.474 | – | – | 8.115 | 55.812 | 29.594 | 173.936 | 88 |
| 121.729 | 56.435 | 32.954 | 8.13 | 55.744 | 32.225 | 175.146 | 116 |
| 122.028 | – | – | 8.051 | – | – | 174.24 | 31 |
| 121.947 | 61.839 | 38.471 | 8.001 | 58.919 | 63.473 | 176.628 | 18 |
| 122.368 | 55.089 | 42.504 | 8.292 | 62.98 | 32.014 | 174.778 | 222 |
| 122.768 | 55.539 | 42.204 | 8.491 | 62.654 | 34.16 | 175.63 | 102 |
| 123.2 | 53.505 | 28.915 | 8.452 | 55.648 | 29.608 | 175.834 | 45 |
| 123.17 | – | – | 8.399 | – | – | 175.591 | 23 |
| 123.004 | – | – | 8.173 | – | – | 176.777 | 61 |
| 124.067 | 53.439 | 19.006 | 8.204 | 54.732 | 41.069 | 174.717 | 169 |
| 124.13 | 58.699 | 38.897 | 8.226 | 57.967 | 63.851 | 177.522 | 77 |
| 124.064 | 53.783 | 18.844 | 8.257 | 54.864 | 41.121 | 174.456 | 169 |
| 124.151 | 53.895 | 18.836 | 8.296 | 54.776 | 41.119 | 174.478 | 198 |
| 124.652 | – | – | 8.423 | – | – | 175.371 | 23 |
| 121.964 | 56.136 | – | 8.397 | 55.477 | 42.153 | 173.951 | 24 |
| 120.576 | – | – | 8.826 | – | – | 175.595 | 8 |
| 122.825 | – | – | 8.662 | – | – | – | 21 |
| 115.232 | – | – | 8.551 | – | – | 175.27 | 7 |
| 120.451 | – | – | 7.801 | – | – | 175.755 | 10 |
| 122.963 | – | – | 7.975 | – | – | 178.479 | 8 |
| 108.758 | – | – | 8.416 | 45.745 | – | 176.258 | 10 |
| 109.149 | 45.559 | – | 8.395 | – | – | 175.577 | 7 |
| 115.98 | 53.782 | 38.852 | 8.313 | 53.546 | 38.762 | 175.932 | 149 |
| 108.665 | 44.698 | – | 7.976 | – | – | 177.222 | 28 |
| 114.28 | 45.445 | – | 8.37 | 54.681 | 40.932 | 175.101 | 22 |
| 116.336 | 55.584 | – | 8.077 | 56.323 | 30.687 | 176.168 | 41 |
| 120.896 | 56.989 | 29.802 | 8.674 | 63.393 | 32.303 | 174.013 | 70 |
| 122.269 | – | – | 8.857 | – | – | 175.083 | 15 |
| 124.22 | 52.779 | 30.062 | 8.44 | 62.877 | 27.555 | 176.349 | 53 |
| 124.327 | 54.821 | 40.994 | 8.026 | 56.536 | 21.942 | 176.026 | 17 |
| 115.823 | 42.951 | – | 7.984 | 56.073 | 38.826 | 175.852 | 25 |
| 122.688 | – | – | 8.576 | 62.694 | – | 175.411 | 32 |
| 121.491 | – | – | 8.519 | 63.134 | 32.343 | 174.605 | 26 |

##### *References*

Delaglio, F., Grzesiek, S., Vuister, G., Zhu, G., Pfeifer, J., Bax, A. NMRPipe: A multidimensional spectral processing system based on UNIX pipes. *J. Biomol. NMR* 1995; **6**: 315–327.

Hyberts, S.G., Milbradt, A.G., Wagner, A.B., Arthanari, H., Wagner, G. Application of iterative soft thresholding for fast reconstruction of NMR data non-uniformly sampled with multidimensional Poisson Gap scheduling. J. Biomol. NMR 2012; **52**: 315–327.

Lakomek, N.-A., Ying, J., Bax, A. Measurement of 15N relaxation rates in perdeuterated proteins by TROSY-based methods. *J. Biomol. NMR* 2012; **53**: 209–221.

Lee, W., Tonelli, M., Markley, J.L. NMRFAM-SPARKY: Enhanced software for biomolecular NMR spectroscopy. *Bioinformatics* 2015; **31**: 1325–1327.

Marty, M.T., Baldwin, A.J., Marklund, E.G., Hochberg, G.K.A., Benesch, J.L.P., Robinson, C. V. Bayesian deconvolution of mass and ion mobility spectra: From binary interactions to polydisperse ensembles. *Anal. Chem.* 2015; **87**: 4370–4376.

Mureddu, L., Vuister, G.W. Simple high-resolution NMR spectroscopy as a tool in molecular biology. *FEBS J.* 2019; **286**: 2035–2042.

Walsh, I., Martin, A.J.M., Di Domenico, T., Tosatto, S.C.E. ESpritz: accurate and fast prediction of protein disorder. *Bioinformatics* 2012; **28**: 503–509.

Walsh, I., Seno, F., Tosatto, S.C.E., Trovato, A. PASTA 2.0: An improved server for protein aggregation prediction. *Nucleic Acids Res.* 2014; **42**: 301–307.
